# The Safety and Efficacy of Transdermal Auricular Vagal Nerve Stimulation Earbud Electrodes for Modulating Autonomic Arousal, Attention, Sensory Gating, and Cortical Brain Plasticity in Humans

**DOI:** 10.1101/732529

**Authors:** William J. Tyler, Sarah Wyckoff, Taylor Hearn, Nicholas Hool

## Abstract

Our work was motivated by the goal of developing a Targeted Neuroplasticity Training (TNT) method for enhancing foreign language learning. To this end, our primary effort was to evaluate new and optimized approaches to noninvasive vagal nerve stimulation (VNS). We considered several Human Factors Dimensions to develop methods that would be amenable to comfortable, everyday use in common training environments or contexts. Several approaches to noninvasive or external vagal nerve stimulation have been described. Transcutaneous modulation of the left cervical branch of the vagus nerve can be uncomfortable for users resulting in a distracting experience, which may not be ideal for augmenting plasticity during training. Transdermal auricular vagal nerve stimulation (taVNS) offers another approach by targeting nerve fibers innervating the external ear. Prior methods have described many different approached using electrode clips on the ear or stainless-steel ball electrodes, which can respectively result in mechanical discomfort and electrical stimulus discomfort due to high current densities. Other approaches use carbon-doped or conductive rubbers, which require wetting. This is problematic since small degrees of dehydration cause significant changes in the electrical impedance of the skin-electrode interface. Detailed human cadaveric studies have shown the external auditory meatus or ear canal is highly innervated by branches of the auricular vagus nerve. Therefore, we designed taVNS electrodes that were fabricated as a biocompatible, hydrogel earbud electrodes for unilateral or bilateral use. We then evaluated the safety and efficacy of these approaches across a range of stimulus frequencies and intensities. We further evaluated the influence of this approach on autonomic physiology by recording heart rate, heart rate variability, skin conductance, skin temperature, and respiration rate. We investigated attention using simultaneous EEG and pupillometry during auditory stimulation tasks. We further studied the effects on sensory gating and plasticity by examining EEG brain activity patterns obtained during auditory mismatch negativity tasks. Finally, we investigated the basic safety and tolerability of the methods and approaches. We found that a simple, dry (hydrogel), earbud electrode design is a safe and effective method for achieving taVNS. Given the safety, preliminary efficacy, and comfort outcomes observed, we conclude taVNS approaches using earbud electrodes warrant further development and investigation as a TNT tool, to mediate human-computer interactions, for brain-computer interfaces, and as medical devices for the treatment of pervasive health disorders.

## Introduction

### Neuromodulation-based approached to enhancing foreign language learning and skill training outcomes

Studies examining the neurobiology of language learning have shown diverse brain regions are involved. Functional plasticity in these brain areas appears to be a key determinant in the ability of individuals to learn languages from early childhood through adulthood. From a classical perspective, the activity and plasticity of two well characterized language-related brain regions have long been known to play major roles in our ability to learn, communicate with, and comprehend spoken languages. Broca’s area is necessary for the planning and execution of speech while Wernicke’s area is required for the analysis and identification of speech [1, 2]. Stemming from several decades of neurophysiological and neuroimaging studies however, we have expanded our understanding of language learning to include many sparsely distributed anatomical regions. For example the perisylvian cortex, superior and middle temporal gyri, the lingual and fusiform gyri, inferior temporal cortex, dorsolateral prefrontal cortex, insula, central operculum, anterior cingulate, and basal ganglia have been shown to play roles in learning, listening, speaking, repeating, and comprehension of spoken language [1, 2]. Further, primary sensory brain regions like the auditory cortex and memory-encoding or information consolidation regions like the hippocampus are also required for the acquisition and proficient use of language. Therefore neuromodulation-based strategies designed to enhance language learning might benefit from influencing the activity and plasticity of multiple linguistically-relevant anatomical loci, spatially distributed cognitive control networks, and sensory processing regions in a simultaneous or coordinated manner.

Several methods of enhancing language learning have been described in the literature. Major advances have been made over the past decade in the design and use of autonomous learning methods, such as those currently incorporated in mobile-based language training applications or computer-assisted language learning programs [3–5]. More recent advances in virtually immersive or virtual reality (VR) technologies have also proven useful for enhancing foreign language learning [6, 7]. While these VR and computer-assisted cognitive training methods can enhance language acquisition by facilitating natural brain activity patterns arising from repeated training sessions or sensory stimulation, they do not directly augment brain activity or plasticity to facilitate learning *per se*. Sleep has also been shown to play a critical role in the consolidation learned information including spoken-language learning [8]. Recent efforts to influence the consolidation phases of language learning during sleep have also shown promise by using targeted memory reactivation (TMR). These TMR investigations demonstrated word cuing during certain stages of sleep can enhance vocabulary learning [9–11]. These TMR methods require the implementation of electroencephalography (EEG) to identify certain stages of sleep, such as slow wave sleep during which sensory information or linguistic cues are presented to enhance language training outcomes. Using EEG during sleep poses several technical challenges that can interfere with broad adoption for enhancing language learning. Other noninvasive neuromodulation methods, which have been developed to directly augment brain activity and plasticity have also been used to enhance language learning.

Transcranial magnetic stimulation (TMS) and transcranial direct current stimulation (tDCS) are two forms of noninvasive neuromodulation that can target restricted regions of cortex to influence brain activity and plasticity. Studies have demonstrated that TMS and tDCS delivered to classical language brain regions like Wernicke’s and Broca’s areas can enhance plasticity and language recovery in patients suffering from post-stroke aphasias [12]. In non-aphasic, healthy volunteers tDCS of Wernicke’s area has been shown to improve verbal memory, enhance word retrieval, and reduce interference during word learning [13, 14]. Also in healthy adults tDCS has been shown to enhance language learning when applied to the posterior part of the left peri-sylvian area [15], enhance language performance when delivered to the left prefrontal cortex [16], improve word retrieval when applied to the primary motor cortex of older adults [17], and to facilitate the learning and maintenance of a novel vocabulary when administered acutely to the temporo-parietal junction on several consecutive days [18]. Despite the success of these methods they are restricted by their coarse targeting abilities, can only modulate one cortical network at a time, and are limited to modulating top-down cortical processes since they are incapable of reaching deeper brain structures.

With the major objective of developing Targeted Neuroplasticity Training (TNT) methods for accelerating foreign language learning, we began to explore the utility of cranial nerve neuromodulation modalities, such as vagal nerve stimulation (VNS). Stimulation of the cervical and auricular branches of the vagus nerve has been repeatedly demonstrated to induce short-term and long-term brain plasticity [19]. It has also been shown to modulate brain circuits and trigger signaling consequences supporting mechanisms of action for the observed plasticity [20–23]. Superficial branches of the vagus in particular the auricular branch of the vagus nerve (ABVN) are located in anatomical regions such that achieving functional electrical coupling to them can be achieved using a number of medical electrode designs including conventional transdermal electrical nerve stimulation (TENS) approaches.

### Vagus Nerve Stimulation for Enhancing Human Brain Plasticity

Electrical stimulation of vagal afferents in several animal species, including humans, has been shown to exert prominent bottom-up actions on key neuromodulatory brain circuits, such as the locus coeruleus (LC) and dorsal raphe (DR), as well as powerful endogenous neurotransmitters like norepinephrine (NE) and serotonin (5-HT) that are known to regulate arousal, attention, and neurophysiological/behavioral plasticity (Figure 1; [20–31]. Further, data from several studies has shown that electrical modulation of vagal afferents, including the ABVN, in healthy humans can safely produce biochemical, behavioral, and neurophysiological effects that are consistent with an enhancement of learning or skill training [32–34]. Based on this collective evidence we hypothesized that taVNS may represent a human factor optimized approach to achieving TNT enhancement of language learning.

**Figure 1.**
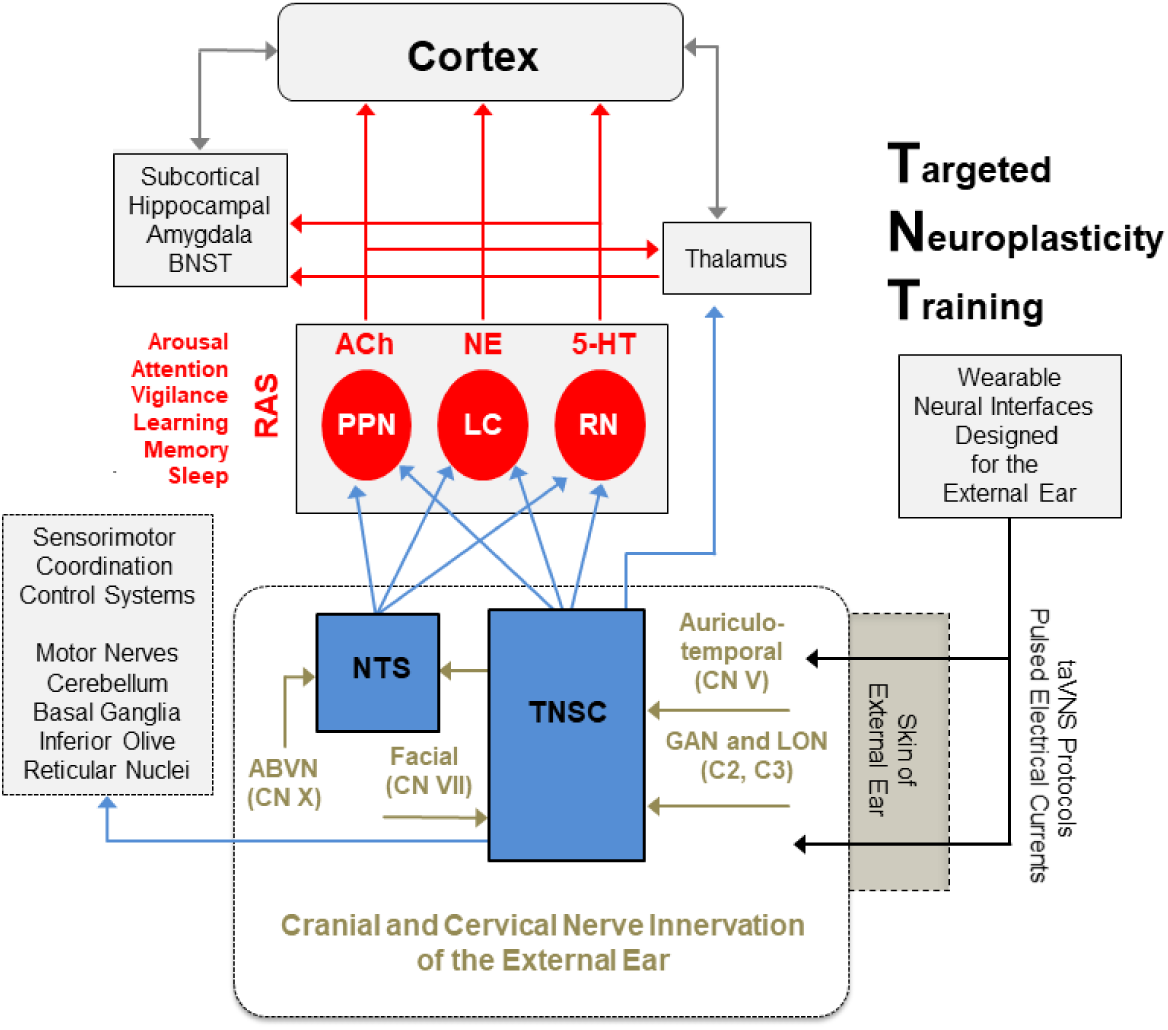
Modulation of peripheral nerves of the external ear for Targeted Neuroplasticity Training. Transcutaneous electrical stimulation of the auricular branch of the vagus and other nerves influence ascending brain activity and arousal systems to enhance skilled training and learning.

One of the major hurdles for developing VNS methods and systems intended to optimize skill training and learning will be to identify the optimal stimulation parameters required to drive acute neural plasticity. While the VNS parameters used across investigations are somewhat uniform, there are not standardized VNS protocols. Systematic studies have not been conducted to identify the best VNS parameters for optimizing neuromodulator signaling or plasticity in the healthy adult brain. Many of the VNS parameters that have been used to date were originally identified and adopted for their ability to desynchronize or interfere with aberrant neural activity like that, which occurs in diseased brain states, such as epilepsy, tinnitus, and depression. In fact, the VNS studies that have shown effects on plasticity have largely borrowed their stimulation devices and parameters from the clinical designs and practices [21,27,32–35]. This poses somewhat of an issue when developing a VNS system for enhancing normal plasticity related to training and learning. It is simply not known whether clinically identified and validated neurostimulation parameters are ideally suited for VNS applications where optimizing normal brain function is the desired outcome. Therefore, a significant amount of our performance effort has been devoted to investigating how different VNS strategies affect sensory or auditory plasticity and autonomic arousal.

### Targeting Auricular Branches of the Human Vagus Nerve

Anatomical and functional studies have shown the afferents of the AVBN innervate the superficial regions of the auricle specifically along the external meatus and concha (Figures 1 and 2A). For example, a study investigating the anatomical evidence for the involvement of the AVBN in Arnold’s cough-reflex showed branches of the AVBN project to the meatus in cadaver microdissection and high-resolution CT imaging studies [36]. Another eloquent histological study conducted on human tissues presented robust histological and immunocytological evidence for the presence of vagal fibers lining the skin outside of the cartilage of the meatus [37]. The presence of fewer fibers was observed in the skin of the concha and moreover those nerves were deeply embedded in the cartilage [37]. Based on this evidence we hypothesized the auditory meatus to be an idealized target for taVNS since there is a higher density of nerve fibers in this location and since cartilage will not interfere with the delivery of pulsed currents to the nerves as much as in other external ear regions. Therefore we engineered, optimized, designed and fabricated biocompatible, hydrogel-based earbud electrodes designed for wearing in the auditory meatus to gain preliminary insights into the safety and efficacy of this method for enhancing sensory processing and brain plasticity. We specifically investigated how varied taVNS stimulus protocols delivered through earbud electrodes influence the safety, tolerability, acute auditory plasticity, and autonomic physiology of healthy human volunteers.

### Human Dimensions & Real-world Considerations for Auricular Vagus Nerve Stimulation Methods

Transdermal auricular vagal nerve stimulation (taVNS) has been shown to be a largely safe and effective method of regulating ascending neuromodulatory brainstem circuits, as well as higher cortical regions in clinical populations. Evidence has suggested this safety extends into a healthy population since VNS has shown be safe and effective for modulating brain activity in healthy adults in many studies now. A potential hurdle to using previously methods described is that they use misguided targeting approaches and they have implemented poor industrial designs that properly account for real world human factors, such as comfort or ability to wear for extended durations.

Many taVNS reports to date have targeted the tragus, for example using crude methods with electrode clips [38]. It remains unclear why other than for convenience or lack of knowledge with respect to functional vagal nerve neuroanatomy. This method is problematic for extended wear due to electrode movement and potential user pain or discomfort. Electrode clips can become quite uncomfortable due to mechanical pinching. Further some of these clip electrodes are made from a high impedance rubber or carbon that is not ideal for noninvasive, transcutaneous electrical nerve modulation. Other fairly coarse approaches have implemented small (1-2 mm) stainless-steel ball electrodes positioned at two or more locations in the concha and external ear [39], which can result in discomfort due to high current densities at the electrode-skin interface. In our investigations, we aimed to ensure wear comfort and comfort during electrical stimulation treatments were optimized as not to distract users or induce pain that may cause significant confounds when trying to enhance brain plasticity or optimize human performance.

Many other reports exist in the field that have implemented poorly designed approaches to electrically interfacing with the skin of the external ear. For instance, Keute and colleagues (2019) used paper or cloth-backed backed electrodes with electrode coupling paste to position small (4 × 4 mm) electrodes in the concha. This again can produce locally high current densities due to the small electrode size used. Further it is a methodological approach subject to numerous confounds and inconsistencies due to unavoidable and significant fluctuations in electrode-skin impedances that will occur with motion or very small movements. Moreover, the electrode materials used were not ideally suited for taVNS. Perhaps most concerning is that the implementation of the above discussed methods can produce unreliable results, cloud the field with murky data, impede future progress towards the development of real world applications, and can influence public perception and funding initiatives.

Therefore, in our investigations, we aimed to utilize Best Practices from state-of-the-art peripheral nerve stimulation methods, as well as modern electrode design and engineering principles. We specifically ensured individualized and proper electrode fits to minimize mechanical movement and electrical impedance fluctuations while maximizing wear comfort and comfort during electrical stimulation treatments. These practices are critical because the methods should not to distract users or induce any pain/discomfort that may cause significant confounds when trying to enhance brain plasticity, modulate arousal and attention, or optimize human performance.

**Figure 2.**
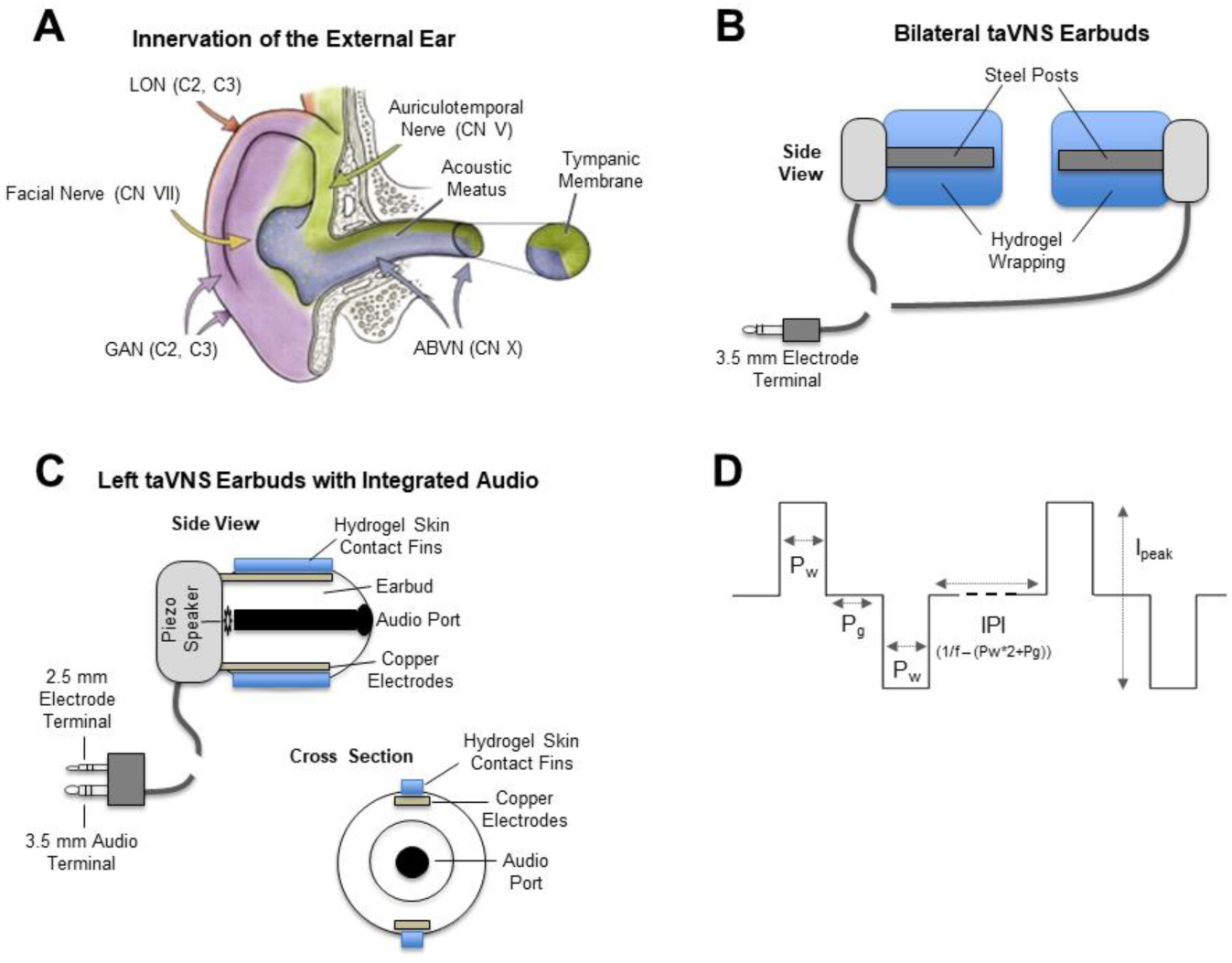
Overview of transdermal auricular vagal nerve stimulation approaches. A) Anatomical illustration showing several branches of cranial nerves (CN V auriculotemporal branch of the trigeminal nerve, CN VII facial nerve, and CN X auricular branch of the vagus nerve = AVBN) and cervical nerves (C2 and C3; lessor occipital nerve = LON and great auricular nerve = GAN) that innervate the external ear. We targeted the ABVN using custom (B) or semi-custom/modified (C) ear bud taVNS electrodes inserted into the acoustic meautus. D) Biphasic, charge-balanced, pulsed electrical currents having varied pulse frequencies and pulse widths (Pw) with an interphase gap (Pg) of 50 microseconds were delivered at different peak current intensities (Ipeak) to the left or both ears through taVNS electrodes after (baseline) and before passive auditory listening tasks while EEG and physiological data were recorded.

## Methods

Below we provide a technical description of the methods and procedures used during our investigations of transdermal auricular vagal nerve stimulation (taVNS).

### Participants

Human subject research participants were recruited for the study at Arizona State University (Tempe, Arizona USA). Following a screening survey to ensure eligibility based on exclusion criteria, subjects provided informed consent and were enrolled in the study. All procedures were approved by the local Institutional Review Board at Arizona State University, as well as by the Human Subjects Research Protection Office at the Naval Information Warfare Center (San Diego, CA USA).

### Inclusion/Exclusion criteria

Participants include adults 18-65 years old. Individuals acknowledging that they do <u>NOT</u> have the following exclusion conditions: undergoing treatment or medication for neurological or psychological disorder, including addiction; have a medical implant (such as a pacemaker, cochlear implant, brain stimulation device); have migraines or frequent headaches; have a history of panic attack or acute anxiety disorder; experience frequent fainting or experienced vaso-vagal syncope or neurocardiogenic syncope even once; have Raynaud’s disease; have Tempromandibular joint (TMJ) disorder or otherfacial neuropathy; have history of concussions or brain injury; have history of significant face/head injury or if you have cranial or facial metal plate or screw implants; have a history of hospitalization for neurological or psychological disorder; had a recent hospitalization for surgery/illness; have vision or hearing that is uncorrectable (corrected vision or hearing is okay); am pregnant; been in a recent drug or alcohol treatment (within past 3 months); have high blood pressure, heart disease, or diabetes.

### Recruitment

Study participants were recruited for Arizona State University student population. Interested participants were screened using an online survey for exclusion criteria and invited to participate in a 75-minute experiment. All participants were asked to arrive for the assessment with clean dry hair. Upon arrival, all participants were explained the study procedures, provided the opportunity to ask questions, and provided written informed consent to participate in the study in accordance with the approval of the Arizona State University Institutional Review Board. Basic gender and age demographic data for subjects participating in the study are provided in Table 1.

**Table 1.** Transdermal auricular vagal nerve modulation stimulus variables and treatment conditions by age and gender. We acquired EEG and psychophysiological data from 120 volunteer human research subjects.

| taVNS Variables |  |  |  |  | N | Gender |  | Age |  |  |
| --- | --- | --- | --- | --- | --- | --- | --- | --- | --- | --- |
| Frequency | Condition | Pulse Width (μsec) | Pulse Gap (μsec) | Intensity (mA; Mean ± SD) |  | Male | Female | Minimum | Maximum | Mean ± SD |
| - | All Active | - | - | 8.55 ± 7.72 | 73 | 49 | 24 | 18 | 50 | 25.11 ± 7.08 |
| 0 Hz | Inactive Sham | - | - | 0.00 ± 0.00 | 31 | 20 | 11 | 18 | 50 | 22.32 ± 5.90 |
| 300 Hz | Ramp Sham | 350 | 50 | 1.75 ± 0.41 | 16 | 11 | 5 | 19 | 48 | 23.81 ± 6.95 |
| 30 Hz | Bilateral | 50 | 50 | 14.42 ± 6.24 | 16 | 13 | 3 | 18 | 39 | 24.81 ± 5.55 |
| 300 Hz | Bilateral | 350 | 50 | 5.83 ± 7.02 | 28 | 19 | 9 | 18 | 50 | 27.79 ± 6.96 |
| 300 Hz | L. Unilateral | 350 | 50 | 1.60 ± 1.71 | 13 | 6 | 7 | 18 | 50 | 26.77 ± 8.99 |
| 3000 Hz | Bilateral | 50 | 50 | 14.14 ± 5.64 | 16 | 11 | 5 | 18 | 48 | 24.63 ± 7.43 |

### Pre-Screening assessment

A 15-question online survey hosted by REDCap assessing the exclusion criteria previously described, as well as age and gender demographics.

### Computer Administered Post-Treatment Survey

A 23-question online survey hosted by REDCap assessing stimulation protocol safety and tolerability with yes/no, 10-point Likert scale, and open-ended questions.

### 24-Hour Post Treatment Survey

A 12-question online survey hosted by REDCap assessing 24-hour stimulation protocol safety and tolerability with yes/no, 10-point Likert scale, and open-ended questions.

### Continuous EEG

For the EEG procedure, participants were seated at a workstation in a private, climate controlled, and sound attenuated recording suite. Participants were instructed to sit comfortably, remain relaxed, and keep their eyes opened and focused on a computer monitor approximately 30 inches in front of them. EEG data was recorded using a Nexus-32 amplifier with Biotrace+ software, version V2017A (mind Media B.V., Netherlands; 24 bit A/D conversion, 2048 Hz sampling rate). Data was acquired from 21 electrode sites (FP1, FP2, F7, F3, Fz, F4, F8, T3(T7), C3, Cz, C4, T4(T8), T5(P7), P3, Pz, P4, T6(P8), O1, O2, A1, and A2) according to the International 10-20 system, using an EEG cap with silver sintered electrodes, referenced in common average, with the ground electrode located at FCz. Four additional latex-free adhesive Ag/AgCl electrodes collected electro-ocular activity. Vertical electro-ocular (VEOG) activity was recorded electrodes attached 1 cm above and below the left eye. Horizontal electro-ocular (HEOG) was recorded with an electrode attached 1 cm beyond the external canthi of the right and left eye (data not shown). We parsed the EEG channel data into three regions and calculated the global averages for these three regions consisting of three EEG recording sites as shown in Figure 16B. Conductive gel was applied to all scalp electrodes and the DC offset was monitored and kept below ± 25,000 µV peak to peak. All data and event markers were exported in EDF+ format for signal processing in third-party software packages including Brain Vision Analyzer 2 (Brain Products GmbH, Germany) and MATLAB (The MathWorks, Inc., Natick, MA, USA).

### Physiological Data

Physiological sensors collected data throughout the recording and included respiration, skin conductance, heart rate, and skin temperature. A Velcro-elastic respiration sensor was used to monitor abdominal breathing (32 Hz sampling rate). Skin conductance (32 Hz sampling rate, µSiemens) was acquired using snap-on pre-gelled Ag/AgCl electrodes attached to the underside of the pointer and middle finger of the left hang. Heart rate and heart rate variability were acquired from electrocardiogram and blood volume pulse sensors. The electrocardiographic (512 Hz sampling rate) data was acquired snap-on pre-gelled Ag/AgCl electrodes attached to the right and left of the upper chest just below the collarbone. The blood volume pulse (128 Hz sampling rate) sensor was clamped on the pointer finger of the left hand. Peripheral hand temperature (32 Hz sampling rate, Fahrenheit) was acquired using a small tipped ceramic thermistor taped to the ring finger of the left hand.

### Pupillometry Data

Pupillometry data was collected in a subset of participants using the Gazepoint GP3 60 Hz machine-vision camera eye tracker (Gazepoint, Vancouver, BC). The distance and visual angle of the eye tracker was adjusted for each participant to meet height and visual detection requirements of the recording software. Prior to the start of the experiment, the recording room lights were dimmed and participants were led through a 5-9 point eye tracking calibration exercise.

### Experimental Tasks

This investigation included two experimental protocols.

Experiment 1 included the following tasks:

- 13 min eyes-open passive auditory oddball (pre-stimulation)
- 10 min eyes-open transcutaneous electrical nerve stimulation
- 13 min eyes-open passive auditory oddball (post-stimulation)
- EEG, EOG, ECG/BVP, respiration, skin conductance level, and hand temperature

Experiment 2 included the following tasks and measures:

- 23 min eyes-open passive auditory oddball (pre-stimulation)
- 10 min eyes-open transcutaneous electrical nerve stimulation
- 23 min eyes-open passive auditory oddball (post-stimulation)
- EEG, Pupillometry, ECG/BVP, respiration, skin conductance level, and hand temperature

### Passive Auditory Mismatch Negativity Stimuli

In Experiment 1, the 13 min passive auditory oddball task was designed to elicit the mismatch negativity (MMN) in an eyes-open condition. The task consisted of a series of randomized *Frequent* (N = 830, 750 Hz) and *Infrequent* (N = 150, 1500 Hz) tones with an inter-stimulus interval of 500 ms. Stimuli were presented over a pair of headphones and participants were instructed to passively attend to the stimuli while they watched a silent nature video. Following pre-stimulation baseline data collection on this task and subsequent taVNS sham or real treatment, this task was repeated after the stimulation task for the post-stimulus time measurements.

In experiment 2, the 23 min passive auditory oddball task was designed to elicit P1, N1, P2, N2, P3, and mismatch negativity (MMN) ERP components in an eyes-open condition. The task consisted of a series of randomized standard/frequent (N = 450, 750 Hz) and deviant/rare (N = 100, 1500 Hz) tones with an inter-stimulus interval of 2500 ms to allow for acquisition of pupillometry data. Stimuli were presented over headphones and participants were instructed to keep their eyes focused on a fixation cross while passively attending to the stimuli. This task was repeated after the stimulation task.

### Transcutaneous Electrical Nerve Stimulation

This study involves several Transcutaneous Electrical Nerve Stimulation (TENS) protocols designed to evaluate how stimulation of different peripheral nerves with different frequency, waveform, current, and timing variables influence auditory processing in the brain. TENS stimulation was generated using a custom-designed experimental generator of charge balanced current pulses (v 2.0, Remi Demers, 2017). Frequency, waveform, current, and timing settings used are shown in Table 1. Following the baseline passive auditory oddball task, the neurostimulation electrodes were placed into the ear(s) and all settings were entered using a MATLAB app. Participants were instructed to increase the current gain in 0.25 mA steps to the maximum level that was comfortable/tolerable or until 20 mA was achieved. Once the gain was set, the participant sat quietly and continued to watch the silent nature video during taVNS stimulation or sham treatment. In our studies we utilized two different control treatment groups. These groups included an inactive sham treatment (no current) and an active placebo (sham) treatment which included a brief (30-sec) current ramp to a low-intensity (sensory suprathreshold) level at the beginning and end of the 10-min stimulation protocol. The mean taVNS current intensity for each treatment group is shown in Table 1.

### Transdermal auricular vagal nerve stimulation electrodes for human use

Several methods have been described for targeting and modulating the ABVN. These approaches use ear clips clamped onto the tragus [40] or stainless-steel balls positioned in the cymbae concha [41]. Ear clip or clamp electrodes can be uncomfortable and tend to slip or change position when worn for extended periods of time, while steel ball electrodes result in locally high current densities (> 2 mA/cm^2^) causing itching, burning sensations, or discomfort for the user. Therefore, in our research and development we focused on engineering methods while considering several human dimensions and implemented design principles to make taVNS a comfortable experience for the end user. We took two approaches to fabricating custom taVNS earbud electrodes designed for insertion into the external acoustic meatus, which is innervated by the ABVN (aka Arnold’s nerve; Figure 2A).

One type of taVNS earbud electrodes were developed using a TENS lead wire (Discount TENS) having two connector cables, each with a steel post connector (2 mm diameter) bent at a 90° angle. The two cables connected to a single 3.5 mm terminal plug that was connected to a stimulus generator as described below. Each steel connector post was wrapped with hydrogel (AG735 or AG2550 hydrogel; Axelgaard Manufacturing Co., Ltd.) to produce earbud electrodes having diameters ranging from 8 to 16 mm and cut to length to ensure a snug fit in each participant’s acoustic meatus (Figure 2B). These earbud electrodes were used to deliver bilateral taVNS as described, while a set of over the ear headphones was used in conjunction with the the stimulating earbuds to deliver audio stimuli during experimental procedures.

Another type of electrode was fashioned to deliver left ear taVNS stimuli integrated into earbud headphones for delivering bilateral audio stimuli. The process involved the disassembly and modification of Nervana headphones (Nervana, LLC, Deerfield Beach, FL). The left earbud of these headphones included a pair of copper stimulating electrodes covered by a conductive silicone sleeve, which was designed to be wet with saline spray before inserting into the meatus. In our pilot studies, we found this wetting procedure to be highly distracting to users and dehydration occurred resulting in poor conductivity. Therefore, we dissembled the left earbuds and inserted layers of hydrogel (AG735 or AG2550 hydrogel) to contact the copper electrode strips while protruding through the silicon earbud sleeve in a manner, which contacted the skin of the acoustic meatus when inserted in the ear (Figure 2C). This procedure was repeated for every participant to ensure the use of fresh hydrogel and a snug fit for each user. The terminal end of the headphones consisted of a 2.5 mm electrode plug that was connected to a stimulus generator for delivering taVNS stimuli as described below. The terminal also included a 3.5 mm audio plug, which was connected to an audio generator for delivering sounds during experimental procedures.

### Signal Processing

Artifact rejection was based on visual and computer selection. During phase-one of the artifact rejection, each epoch was visually inspected and marked for technical artifacts. Computer based artifact rejection was completed using the independent component analysis (ICA) method. During ICA correction, the EEG was decomposed into 20 signal components. Only components that reflected VEOG, HEOG, EKG, and EMG artifacts, without reducing non-artifact signals, were removed. The file was visually inspected a second time to ensure all major artifacts were removed. The artifact free data and event markers were exported and analyzed using the Chronux and EEGLAB toolboxes as well as custom MATLAB software. EEG data was detrended using a local linear regression, and then an 8th order low-pass filter was applied at 55 Hz. The data was then segmented into 750 msec trials aligned on tone-onset (150 msec pre-tone, 600 msec post-tone) and averaged by trial type (pre-same/pre-diff/post-same/post-diff). Multi-taper time-frequency spectra were calculated using a time-bandwidth product of 3, 5 tapers, a moving window of 150 msec, and a step size of 10 msec. Mean pre-tone spectral activity (averaged over time-domain) was then subtracted from each time point in the trial.

### ERP Analysis

Artifact free segments and event markers were exported for analysis using Chronux, EEGLAB, and custom MATLAB software. The data was then segmented into 750 ms trials aligned on tone-onset (150 ms pre-tone, 600 ms post-tone) and averaged by trial type (pre-same/pre-diff/post-same/post-diff). Multi-taper time-frequency spectra were calculated using a time-bandwidth product of 3, 5 tapers, a moving window of 150 ms, and a step size of 10 ms. The pre-tone spectral activity (averaged over time-domain) was then subtracted from each time point in the trial. A grand average of the rare minus frequent tone difference waves was calculated independently for F3, Fz, F4, C3, Cz, C4, P3, Pz, and P4 during the pre-stimulation task. An overlay of the difference waves was used to define the windowing for P1, N1, P2, N2, P3, and MMN component peak amplitude and latency measures and applied to the pre- and post-stimulation oddball data for each participant. Latency (ms) and amplitude (mV) values were exported for statistical analysis.

### Continuous EEG Analysis

Artifact free segments were band-pass filtered at 1-40 Hz (2^nd^ Order IIR filter) and segmented into 10-minute pre- and post-stimulation blocks for spectral analysis. Fast Fourier Transformation (2 sec epochs, 10% Hanning window) was applied and the mean absolute and relative power activity for delta (1-4 Hz), theta (4-8 Hz), alpha (8-12 Hz), beta 1 (12-15 Hz), beta 2 (15-20 Hz), beta 3 (20-30 Hz), and gamma (30-40 Hz) was computed. The mean values for F3, Fz, F4, C3, Cz, C4, P3, Pz, and P4 were exported for statistical analysis.

### Physiological Analysis

ECG/BVP, respiration, skin conductance, and hand temperature data were segmented into 10-minute pre-, stim, and post-stimulation blocks. Due to signal dropout and contamination during the stimulation protocols, ECG data was excluded from the analysis and the BVP signal was used to calculate heart rate/heart-rate variability measures. The BVP signal was filtered (high-pass .5 Hz, low-pass 3 Hz, 2^nd^ order IIR filter) and exported for RR interval detection and analysis in the Kubios HRV Premium software. Time-domain (heart rate, SDNN, RMSSD, HRV (max HR – min HR) amplitude), frequency-domain (normalized low frequency (0.04-0.15 Hz) and high frequency (0.15-0.40 Hz) power, LF/HF ratio), and PNS, SNS, and Stress indexes (Appendix 5) were computed. Respiration activity was filtered (high-pass .05 Hz, low-pass 1 Hz, 2^nd^ order IIR filter) and the level trigger marker transform was applied to mark the inhalation peak of each breath cycle. Respiration intervals were used to calculate breathes per minute. Mean HR/HRV, respiration, skin conductance level, and hand temperature measure were exported for statistical analysis.

### Pupillometry Analysis

Pupil data was preprocessed using methods set forth by Geller, Landrigan, and Mirman (2019). First, samples with unrealistic pupil diameters (<2 mm or >8 mm) pupil diameters greater than or less than three standard deviations from the mean were removed. Samples occurring between 100 ms before and 100 ms after blinks were also marked as invalid. Samples were then linearly interpolated. Any artifacts due to rapid changes in pupil diameter, as determined by median absolute deviation, were then removed (Kret & Sjak-Shie, 2018). A 5-point moving average was applied to smooth the samples. The continuous data was then aligned on tone-onset, converted to z-scores, and baseline corrected by subtracting the median activity 0.5 s before tone-onset from each sample.

### Statistical Analysis of Safety and Tolerability Data

A Pearson’s chi-squared test was used to investigate the relationship between post-stimulation and 24-hour reports (Yes, No) of discomfort, dizziness, blurred vision, skin irritation and Vagus nerve TENS protocols (sham, ramp sham, 30 Hz, 300 Hz unilateral, 300 Hz bilateral, 3000 Hz). A one-way ANOVA was used to compare mean comfort, relaxation, discomfort, dizziness, blurred vision, and skin irritation ratings between each protocol group.

### Statistical Analysis of EEG Data

Separate mixed ANOVA were used to examine the main and simple effects of TENS Protocol, Time, Time*Protocol, and Time*Region*Protocol on peak amplitude and latency values of the P1, N1, P2, N2, and P3 components. The between-subjects “Protocol” factor included vagus TENS protocols (sham, ramp sham, 30 Hz, 300 Hz unilateral, 300 Hz bilateral, 3000 Hz). The within-subjects “Time” factor was defined as pre- and post-stimulation and the “Region” factor was defined as Frontal (F3, Fz, F4), Central (C3, Cz, C4), and Posterior (P3, Pz, P4) pooled electrode sites. Similarly, separate mixed ANOVA were used to examine the main and simple effects of Protocol, Time, Time*Protocol, and Time*Region*Protocol on absolute and relative delta, theta, alpha, beta 1, beta 2, beta 3, and gamma band activity. As multivariate analysis of variance (MANOVA) is not dependent upon assumptions of sphericity, the Wilk’s Lambda multivariate test statistics and p-values were reported. Bonferroni correction was applied to the alpha-level to control for multiple comparisons.

### Statistical Analysis of Autonomic Physiology Data

Separate mixed ANOVA were used to examine the main and simple effects of vagus TENS Protocols, Time (pre-stimulation, stim, post-stimulation), and Time*Protocol on heart rate/HRV measures (SDNN, RMSSD, HF normalized power, LF normalized power, LF/HF ratio, PNS tone, SNS tone, Stress index), respiration rate, hand temperature, and skin conductance. As multivariate analysis of variance (MANOVA) is not dependent upon assumptions of sphericity, the Wilk’s Lambda multivariate test statistics and p-values were reported. Bonferroni correction was applied to the alpha-level to control for multiple comparisons.

## Results

### Influence of taVNS on Autonomic Physiology

Since it has been shown that VNS and other electrical, peripheral (cranial) nerve stimulation methods can affect general autonomic arousal, we first investigated how taVNS affected several measures of autonomic arousal including heart rate (HR), heart rate variability (HRV), skin conductance, skin temperature, and respiration rate. Our analyses of HRV included specific sub-measures: standard deviation of normal-to-normal heart beat (SDNN), root mean squared of the standard deviation (RMSDD), and time-frequency analyses of high-frequency and low-frequency components of HRV. From the HR, HRV, and respiration rate we examined sympathetic and parasympathetic tones (index), as well as a global stress index. The mean +/- SEM for each one of the dependent variables measured to asses changes in autonomic activity produced by treatments are shown in Table 2.

**Table 2.** Measures of autonomic nervous system activity in response to transdermal auricular vagal nerve stimulation.

| Variable Mean ± SEM | Sham (N = 31) | | | Ramp Sham (N = 16) | | | 30 Hz bilateral (N = 16) | | | 300 Hz left bilateral (N = 28) | | | 300 Hz unilateral (N = 13) | | | 3000 Hz bilateral (N = 16) | | |
| --- | --- | --- | --- | --- | --- | --- | --- | --- | --- | --- | --- | --- | --- | --- | --- | --- | --- | --- |
|  | Baseline | Stim | Post | Baseline | Stim | Post-Stim | Baseline | Stim | Post | Baseline | Stim | Post | Baseline | Stim | Post | Baseline | Stim | Post |
| Heart Rate (bpm) | 77.26 ± 1.53 | 75.88 ± 1.51 | 75.71 ± 1.45 | 74.57 ± 2.31 | 72.66 ± 2.22 | 71.76 ± 2.34 | 72.75 ± 2.53 | 70.39 ± 2.49 | 70.39 ± 2.25 | 75.96 ± 2.02 | 74.66 ± 1.98 | 73.99 ± 1.89 | 72.92 ± 1.61 | 72.17 ± 2.11 | 72.94 ± 2.12 | 72.25 ± 2.79 | 70.74 ± 2.71 | 70.54 ± 2.62 |
| SDNN (msec) | 39.87 ± 2.30 | 43.61 ± 2.22 | 44.00 ± 2.57 | 47.40 ± 5.27 | 51.98 ± 4.10 | 51.36 ± 4.66 | 42.20 ± 4.84 | 46.33 ± 4.30 | 48.63 ± 4.53 | 44.51 ± 3.19 | 46.55 ± 2.90 | 49.17 ± 2.91 | 49.24 ± 3.90 | 45.96 ± 3.43 | 46.88 ± 3.66 | 46.37 ± 4.65 | 53.50 ± 5.00 | 56.15 ± 5.76 |
| RMSDD (msec) | 33.53 ± 2.31 | 38.14 ± 2.61 | 36.86 ± 2.87 | 41.81 ± 4.91 | 47.74 ± 4.43 | 48.26 ± 5.02 | 37.70 ± 4.21 | 42.69 ± 4.14 | 44.96 ± 4.44 | 40.74 ± 3.65 | 42.68 ± 3.05 | 45.16 ± 3.50 | 42.43 ± 4.32 | 41.41 ± 4.49 | 40.81 ± 4.15 | 46.77 ± 5.45 | 52.77 ± 6.11 | 52.96 ± 6.30 |
| HRV (HRmax - HRmin) | 26.52 ± 1.37 | 31.16 ± 1.39 | 30.51 ± 1.65 | 27.60 ± 2.02 | 30.93 ± 2.22 | 30.38 ± 2.47 | 25.02 ± 2.41 | 25.99 ± 1.30 | 28.83 ± 3.13 | 28.14 ± 1.37 | 30.30 ± 1.63 | 28.36 ± 1.28 | 29.28 ± 2.12 | 27.00 ± 1.57 | 28.49 ± 2.25 | 25.24 ± 2.01 | 37.55 ± 6.35 | 31.65 ± 2.71 |
| HF Power | 38.22 ± 2.78 | 40.28 ± 2.59 | 36.22 ± 2.52 | 36.68 ± 4.01 | 39.36 ± 3.53 | 39.33 ± 3.57 | 40.86 ± 4.19 | 40.36 ± 3.40 | 39.20 ± 3.26 | 42.38 ± 2.74 | 40.33 ± 2.38 | 37.49 ± 2.39 | 37.77 ± 4.11 | 38.93 ± 4.18 | 38.30 ± 4.97 | 42.62 ± 4.05 | 43.96 ± 3.94 | 38.68 ± 3.13 |
| LF Power | 61.73 ± 2.78 | 59.87 ± 2.59 | 63.73 ± 2.52 | 63.25 ± 4.01 | 60.59 ± 3.53 | 60.59 ± 3.57 | 59.09 ± 4.19 | 59.80 ± 3.40 | 60.78 ± 3.28 | 57.54 ± 2.74 | 59.59 ± 2.36 | 62.44 ± 2.38 | 62.18 ± 4.11 | 61.01 ± 4.18 | 61.65 ± 4.96 | 57.33 ± 4.05 | 55.92 ± 3.96 | 61.25 ± 3.14 |
| LF/HF Ratio | 2.12 ± 0.26 | 1.88 ± 0.22 | 2.52 ± 0.44 | 2.21 ± 0.32 | 1.88 ± 0.28 | 1.87 ± 0.27 | 1.84 ± 0.28 | 1.83 ± 0.31 | 1.85 ± 0.28 | 1.67 ± 0.20 | 1.77 ± 0.20 | 1.98 ± 0.19 | 2.02 ± 0.31 | 2.09 ± 0.44 | 2.31 ± 0.49 | 1.85 ± 0.42 | 1.71 ± 0.36 | 1.96 ± 0.35 |
| PNS Tone | -0.86 ± 0.13 | -0.66 ± 0.14 | -0.70 ± 0.13 | -0.48 ± 0.24 | -0.22 ± 0.22 | -0.14 ± 0.25 | -0.49 ± 0.23 | -0.21 ± 0.24 | -0.16 ± 0.23 | -0.56 ± 0.19 | -0.44 ± 0.20 | -0.34 ± 0.19 | -0.43 ± 0.18 | -0.38 ± 0.22 | -0.45 ± 0.20 | -0.18 ± 0.28 | 0.06 ± 0.28 | 0.06 ± 0.29 |
| SNS Tone | 1.00 ± 0.16 | 0.76 ± 0.16 | 0.75 ± 0.16 | 0.62 ± 0.25 | 0.33 ± 0.21 | 0.25 ± 0.22 | 0.71 ± 0.31 | 0.29 ± 0.26 | 0.24 ± 0.25 | 0.80 ± 0.23 | 0.60 ± 0.21 | 0.51 ± 0.21 | 0.47 ± 0.20 | 0.44 ± 0.21 | 0.48 ± 0.20 | 0.40 ± 0.28 | 0.20 ± 0.29 | 0.17 ± 0.30 |
| Stress Index | 10.97 ± 0.58 | 10.06 ± 0.51 | 10.01 ± 0.66 | 9.72 ± 0.86 | 8.72 ± 0.61 | 8.01 ± 0.60 | 11.05 ± 1.06 | 9.38 ± 0.79 | 9.08 ± 0.77 | 10.33 ± 0.78 | 9.57 ± 0.61 | 9.31 ± 0.71 | 9.36 ± 1.01 | 9.59 ± 0.69 | 9.47 ± 0.76 | 9.46 ± 1.05 | 8.80 ± 0.96 | 8.62 ± 1.01 |
| Resp Rate (bpm) | 16.45 ± 0.59 | 16.47 ± 0.56 | 16.43 ± 0.59 | 16.22 ± 1.02 | 16.11 ± 0.94 | 16.38 ± 0.87 | 15.87 ± 0.88 | 16.16 ± 0.86 | 16.29 ± 0.66 | 16.80 ± 0.59 | 16.93 ± 0.59 | 16.80 ± 0.59 | 16.01 ± 0.69 | 16.61 ± 0.74 | 16.33 ± 0.89 | 16.51 ± 0.86 | 16.36 ± 0.96 | 16.14 ± 0.90 |
| Hand Temp (°F) | 93.47 ± 0.84 | 91.76 ± 1.04 | 90.66 ± 1.20 | 92.40 ± 1.15 | 90.74 ± 1.40 | 89.14 ± 1.53 | 93.85 ± 0.56 | 92.68 ± 0.90 | 91.77 ± 1.12 | 95.22 ± 0.28 | 94.67 ± 0.31 | 94.43 ± 0.30 | 94.82 ± 0.53 | 94.04 ± 0.70 | 94.45 ± 0.54 | 93.03 ± 0.91 | 91.82 ± 1.15 | 91.18 ± 1.13 |
| Skin Conductance (μS) | 4.57 ± 1.04 | 7.19 ± 1.78 | 6.59 ± 1.77 | 6.38 ± 2.78 | 16.05 ± 5.96 | 15.71 ± 6.04 | 8.13 ± 3.26 | 10.33 ± 3.68 | 7.96 ± 3.55 | 9.45 ± 2.93 | 16.69 ± 5.96 | 20.16 ± 6.56 | 8.07 ± 1.83 | 7.61 ± 1.45 | 10.16 ± 2.42 | 13.32 ± 4.54 | 14.32 ± 5.91 | 14.97 ± 6.42 |

Statistical analyses by ANOVA revealed significant main effects of Time (baseline, stimulation, post-stimulation) on HR, SDNN, RMSDD, HRV, PNS Tone, SNS tone, stress index, and skin (hand) temperature. A significant main effect of taVNS Frequency on skin temperature and significant Time x taVNS Frequency interactions on HRV and skin temperature were also revealed by ANOVA (Table 3). The simple effects of Frequency revealed that hand temperature was significantly changed from baseline during the taVNS stimulation and post-stimulation Time conditions. The simple effects of Time showed that many treatment conditions including sham treatments significantly affected autonomic activity. We interpret these results due to a natural relaxation that occurs for subjects throughout the 60-75 minute experimental protocol. Subjects are asked to remain passive during these periods and significant psychophysiological relaxation occurs from the time baseline data begin being collected. For example, even the inactive sham treatment produced a significant decrease in heart rate and other autonomic measures across the Time factors (variables) of the experimental protocol. Data are summarized in Table 3. Data are illustrated as histograms showing mean values for 10 min baseline, taVNS stimulation, and post-stimulation treatment Time periods each in Figures 3 – 15 below. Also illustrated as histograms in those figures are percent changes from baseline for each treatment condition.

**Table 3.** Statistical results table of P-values from treatment Time*taVNS Frequency ANOVAs. Statistically significant results are indicated where p < 0.05 by bold text in a grey shaded cell.

| Variable Mean ± SEM | Main Effects (P-values) | | | Simple Effects of Frequency (P-values) | | | Simple Effects of Time (P-values) | | | | | |
| --- | --- | --- | --- | --- | --- | --- | --- | --- | --- | --- | --- | --- |
|  | Time | Frequency | Time*Frequency | Baseline | Stim | Post | Sham | Ramp Sham | 30 Hz bilateral | 300 Hz bilateral | 300 Hz unilateral | 3000 Hz bilateral |
| Heart Rate (bpm) | < 0.001 | 0.332 | 0.802 | 0.436 | 0.33 | 0.336 | 0.036 | 0.005 | 0.017 | 0.01 | 0.852 | 0.121 |
| SDNN (msec) | < 0.001 | 0.339 | 0.249 | 0.4 | 0.311 | 0.31 | 0.021 | 0.149 | 0.015 | 0.017 | 0.516 | 0.002 |
| RMSSD (msec) | < 0.001 | 0.125 | 0.272 | 0.21 | 0.162 | 0.086 | 0.037 | 0.009 | 0.004 | 0.026 | 0.794 | 0.012 |
| HRV (HRmax - HRmin) | < 0.001 | 0.633 | 0.029 | 0.605 | 0.096 | 0.861 | 0.008 | 0.275 | 0.177 | 0.578 | 0.748 | < 0.001 |
| HF Power | 0.106 | 0.957 | 0.613 | 0.765 | 0.94 | 0.974 | 0.152 | 0.504 | 0.827 | 0.062 | 0.925 | 0.167 |
| LF Power | 0.105 | 0.956 | 0.612 | 0.765 | 0.936 | 0.973 | 0.152 | 0.508 | 0.826 | 0.062 | 0.924 | 0.164 |
| LF/HF Ratio | 0.161 | 0.887 | 0.615 | 0.744 | 0.975 | 0.684 | 0.016 | 0.322 | 0.997 | 0.417 | 0.699 | 0.667 |
| PNS Tone | < 0.001 | 0.14 | 0.43 | 0.242 | 0.192 | 0.103 | 0.042 | 0.001 | 0.003 | 0.01 | 0.911 | 0.029 |
| SNS Tone | < 0.001 | 0.366 | 0.637 | 0.47 | 0.385 | 0.33 | 0.02 | 0.012 | 0.001 | 0.01 | 0.975 | 0.174 |
| Stress Index | < 0.001 | 0.676 | 0.779 | 0.622 | 0.708 | 0.707 | 0.059 | 0.146 | 0.004 | 0.078 | 0.095 | 0.358 |
| Resp Rate (bpm) | 0.729 | 0.941 | 0.961 | 0.961 | 0.969 | 0.99 | 0.983 | 0.781 | 0.563 | 0.882 | 0.443 | 0.618 |
| Hand Temp (°F) | < 0.001 | 0.016 | 0.025 | 0.094 | 0.036 | 0.003 | < 0.001 | < 0.001 | 0.004 | 0.2 | 0.246 | 0.009 |
| Skin Conductance (μS) | 0.06 | 0.323 | 0.052 | 0.318 | 0.477 | 0.245 | 0.654 | 0.009 | 0.596 | 0.014 | 0.675 | 0.946 |

**Figure 3.**
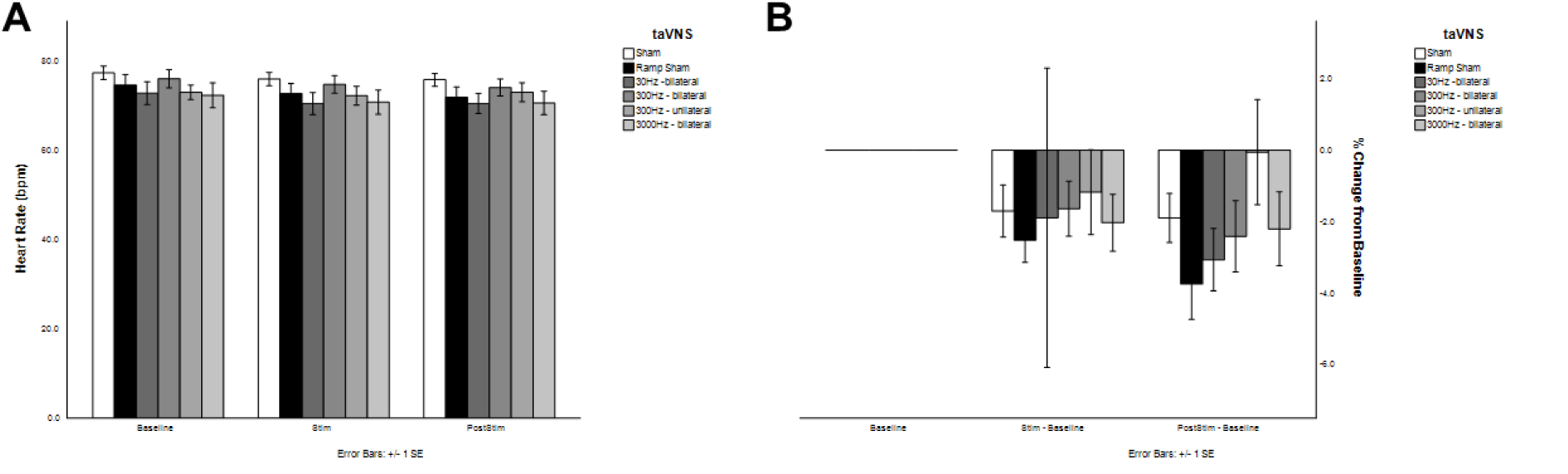
Influence of transdermal auricular vagal nerve stimulation on heart rate. A) Data illustrate mean HR in beats-per-minute (bpm) during 10-min baseline, taVNS stimulation, and post-stimulus periods for the inactive sham (sham), ramp sham, 30 Hz bilateral taVNS, 300 Hz bilateral taVNS, 300 Hz unilateral taVNS, and 3000 Hz taVNS. B) Data from panel A illustrated as a percent change from baseline for the taVNS and post-stimulus time periods across all taVNS treatment conditions.

**Figure 4.**
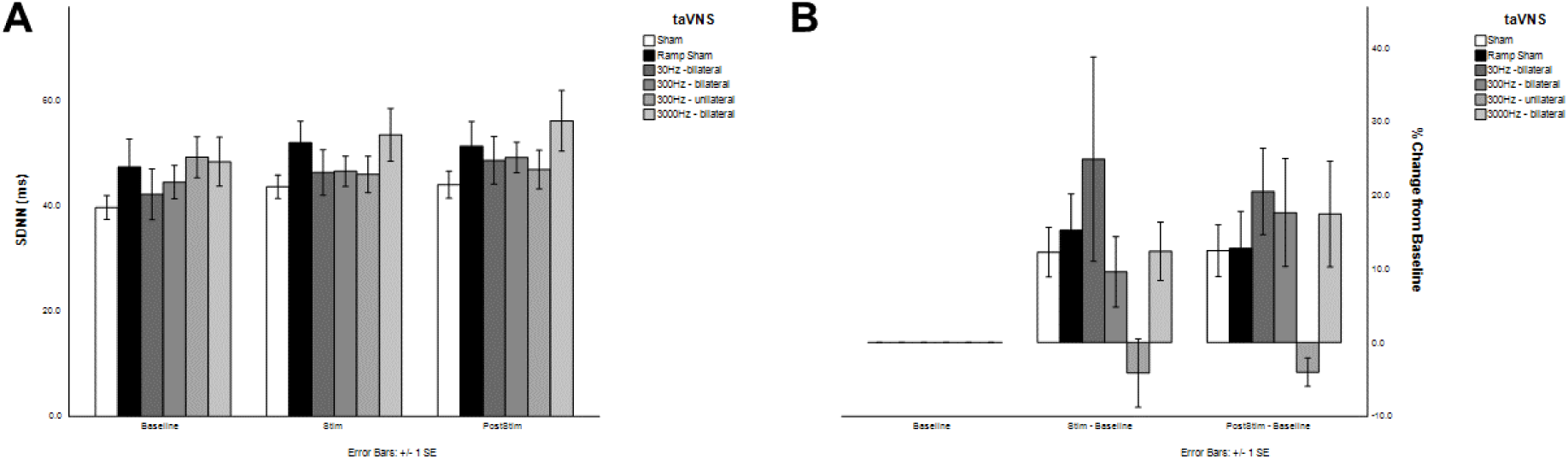
Influence of transdermal auricular vagal nerve stimulation on standard deviation of normal-to-normal (SDNN) heart beats. A) Data illustrate mean SDNN in msec during 10-min baseline, taVNS stimulation, and post-stimulus periods for the inactive sham (sham), ramp sham, 30 Hz bilateral taVNS, 300 Hz bilateral taVNS, 300 Hz unilateral taVNS, and 3000 Hz taVNS. B) Data from panel A illustrated as a percent change from baseline for the taVNS and post-stimulus time periods across all taVNS treatment conditions.

**Figure 5.**
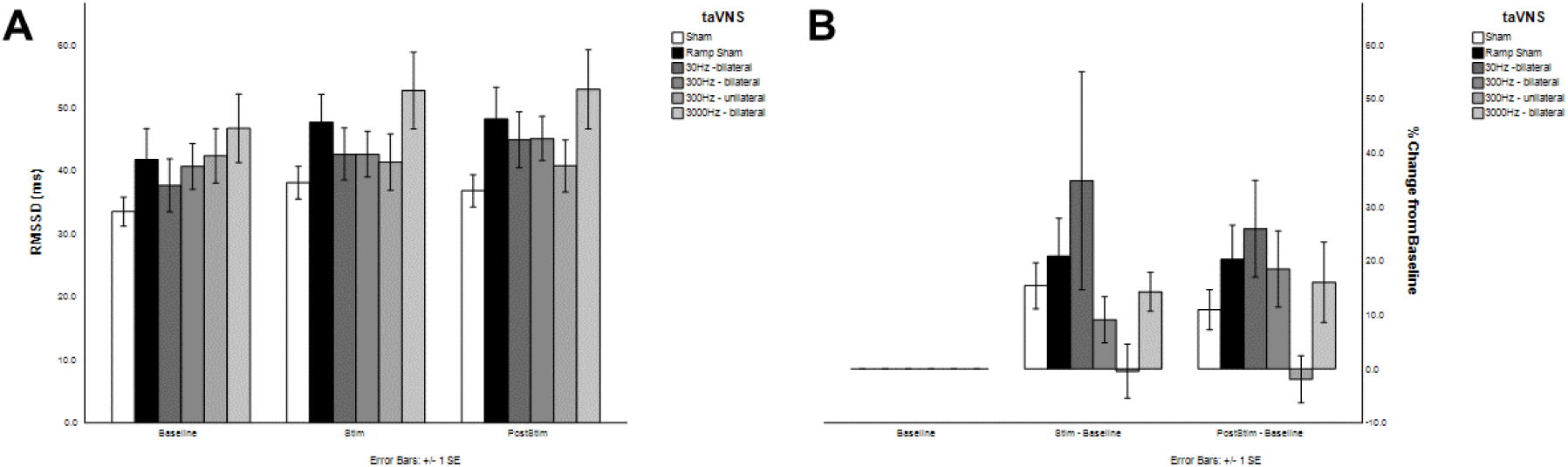
Influence of transdermal auricular vagal nerve stimulation on the root mean squared standard deviation (RMSDD) of heart rate. A) Data illustrate mean RMSDD in msec during 10- min baseline, taVNS stimulation, and post-stimulus periods for the inactive sham (sham), ramp sham, 30 Hz bilateral taVNS, 300 Hz bilateral taVNS, 300 Hz unilateral taVNS, and 3000 Hz taVNS. B) Data from panel A illustrated as a percent change from baseline for the taVNS and post-stimulus time periods across all taVNS treatment conditions.

**Figure 6.**
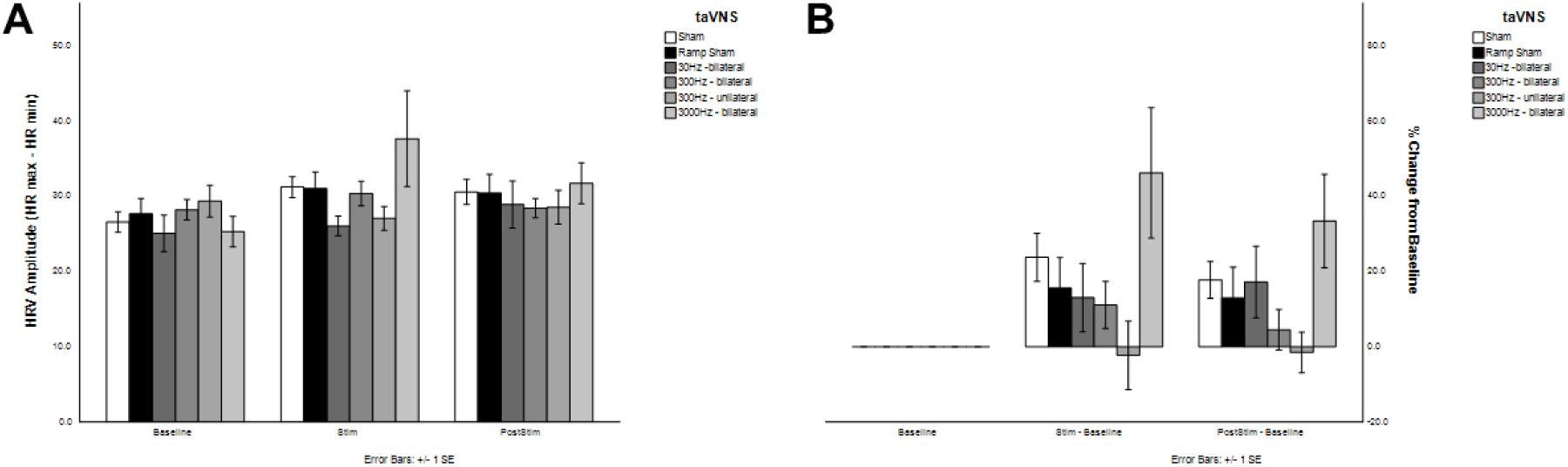
Influence of transdermal auricular vagal nerve stimulation on heart rate variability (HRV) amplitude. A) Data illustrate mean HRV amplitude (HRmax – HRmin) during 10-min baseline, taVNS stimulation, and post-stimulus periods for the inactive sham (sham), ramp sham, 30 Hz bilateral taVNS, 300 Hz bilateral taVNS, 300 Hz unilateral taVNS, and 3000 Hz taVNS. B) Data from panel A illustrated as a percent change from baseline for the taVNS and post-stimulus time periods across all taVNS treatment conditions.

**Figure 7.**
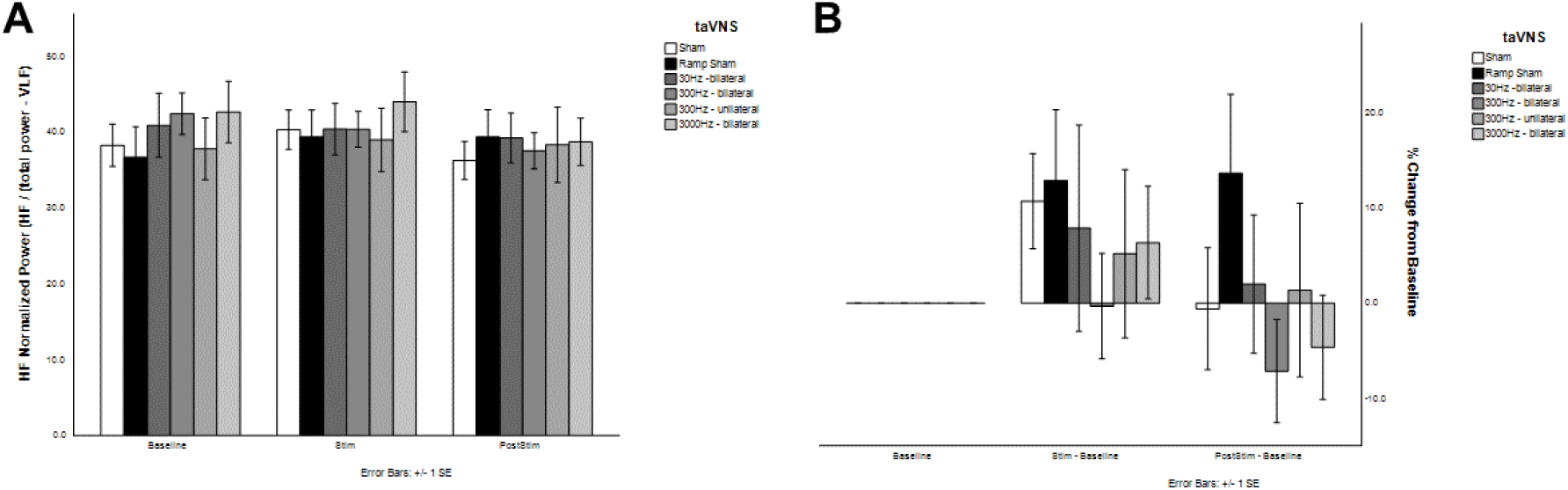
Influence of transdermal auricular vagal nerve stimulation on the high frequency (HF) normalized power of HRV. A) Data illustrate mean HRV normalized power (HRV HF power / total power – VLF power) during 10-min baseline, taVNS stimulation, and post-stimulus periods for the inactive sham (sham), ramp sham, 30 Hz bilateral taVNS, 300 Hz bilateral taVNS, 300 Hz unilateral taVNS, and 3000 Hz taVNS. B) Data from panel A illustrated as a percent change from baseline for the taVNS and post-stimulus time periods across all taVNS treatment conditions.

**Figure 8.**
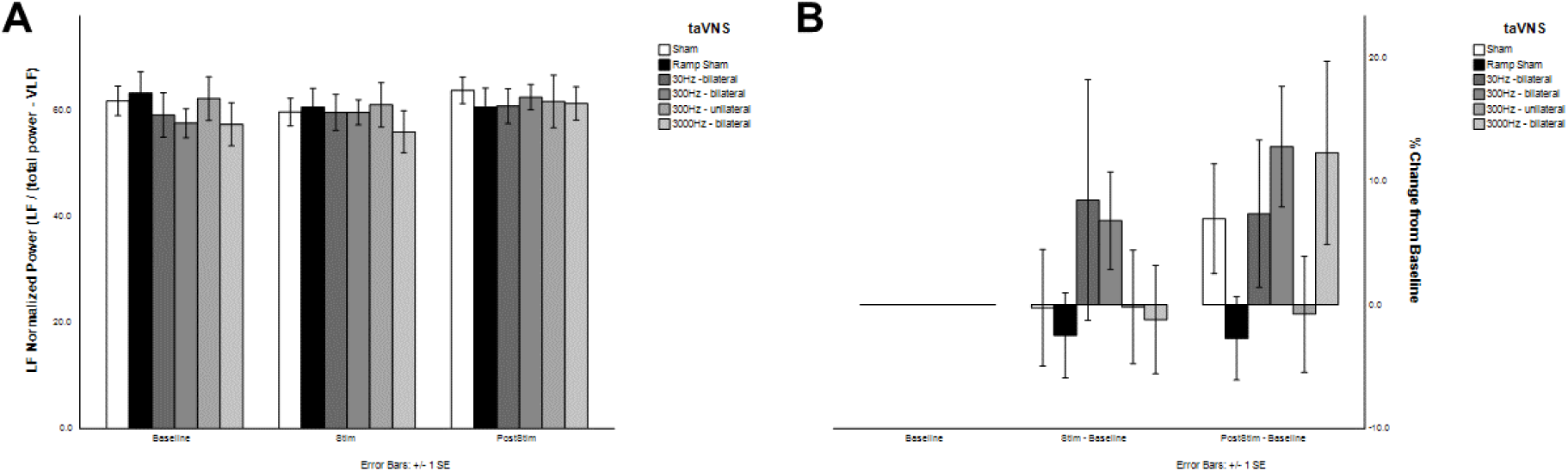
Influence of transdermal auricular vagal nerve stimulation on the low frequency (LF) normalized power of HRV. A) Data illustrate mean HRV normalized power (HRV LF power / total power – VLF power) during 10-min baseline, taVNS stimulation, and post-stimulus periods for the inactive sham (sham), ramp sham, 30 Hz bilateral taVNS, 300 Hz bilateral taVNS, 300 Hz unilateral taVNS, and 3000 Hz taVNS. B) Data from panel A illustrated as a percent change from baseline for the taVNS and post-stimulus time periods across all taVNS treatment conditions.

**Figure 9.**
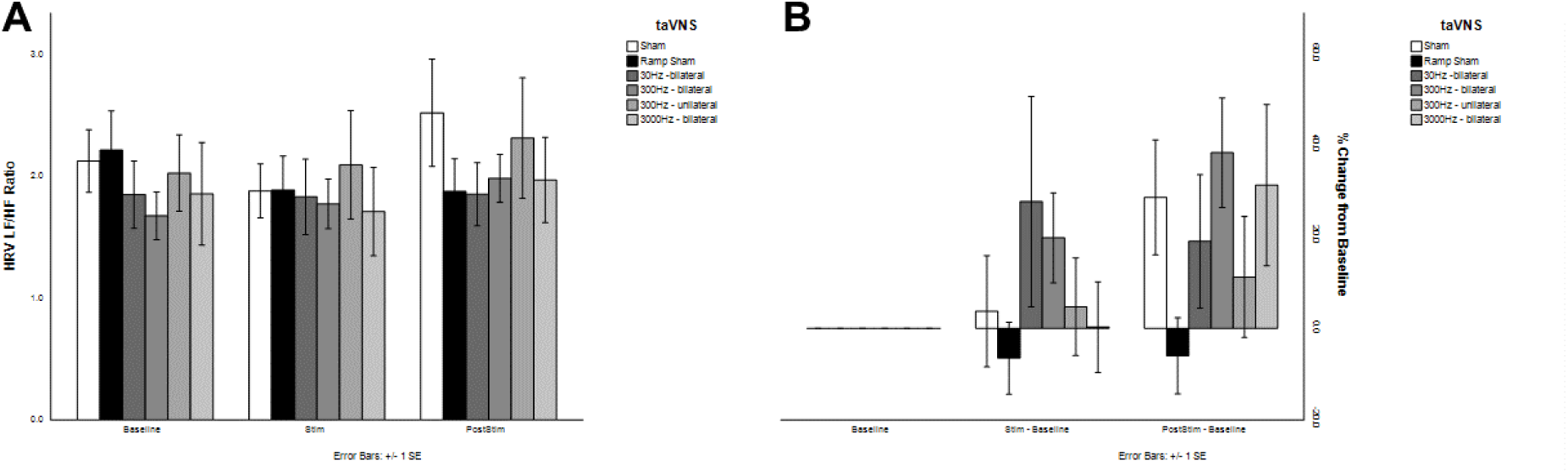
Influence of transdermal auricular vagal nerve stimulation on the HRV HF / LF ratio. A) Data illustrate mean HRV HF / LF power ratios during 10-min baseline, taVNS stimulation, and post-stimulus periods for the inactive sham (sham), ramp sham, 30 Hz bilateral taVNS, 300 Hz bilateral taVNS, 300 Hz unilateral taVNS, and 3000 Hz taVNS. B) Data from panel A illustrated as a percent change from baseline for the taVNS and post-stimulus time periods across all taVNS treatment conditions.

**Figure 10.**
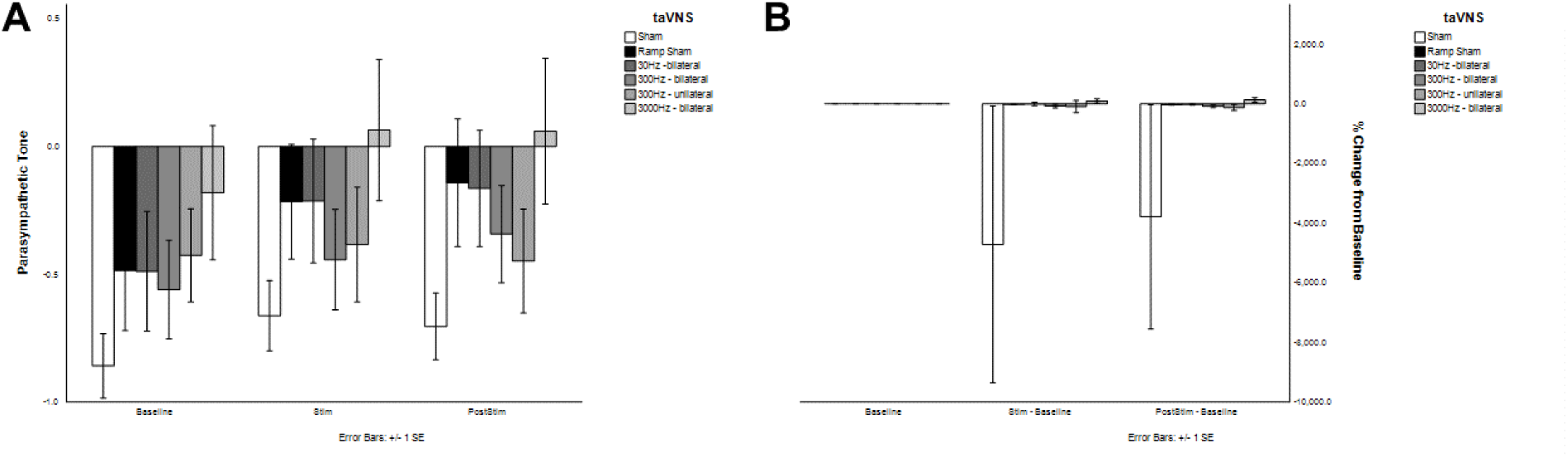
Influence of transdermal auricular vagal nerve stimulation on parasympathetic tone. A) Data illustrate mean parasympathetic tone (index derived using Kubios software from Mean RR, RMSSD and SD1 %) during 10-min baseline, taVNS stimulation, and post-stimulus periods for the inactive sham (sham), ramp sham, 30 Hz bilateral taVNS, 300 Hz bilateral taVNS, 300 Hz unilateral taVNS, and 3000 Hz taVNS. B) Data from panel A illustrated as a percent change from baseline for the taVNS and post-stimulus time periods across all taVNS treatment conditions.

**Figure 11.**
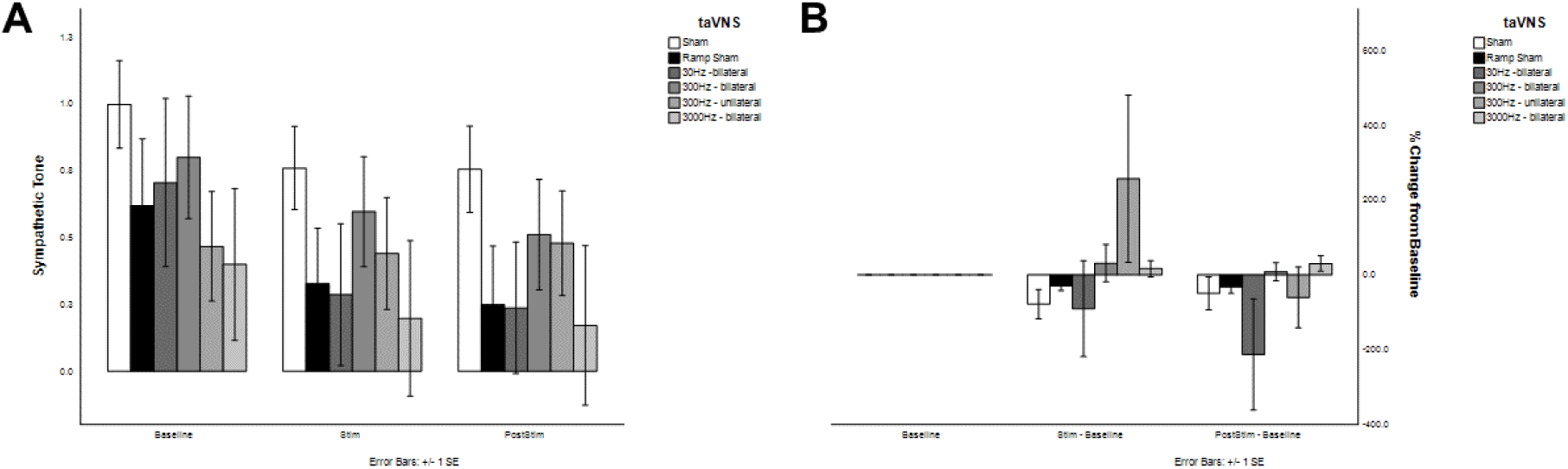
Influence of transdermal auricular vagal nerve stimulation on sympathetic tone. A) Data illustrate mean sympathetic tone (index derived using Kubios software from Mean HR, Stress Index (Figure 12) and SD2 %) during 10-min baseline, taVNS stimulation, and post-stimulus periods for the inactive sham (sham), ramp sham, 30 Hz bilateral taVNS, 300 Hz bilateral taVNS, 300 Hz unilateral taVNS, and 3000 Hz taVNS. B) Data from panel A illustrated as a percent change from baseline for the taVNS and post-stimulus time periods across all taVNS treatment conditions.

**Figure 12.**
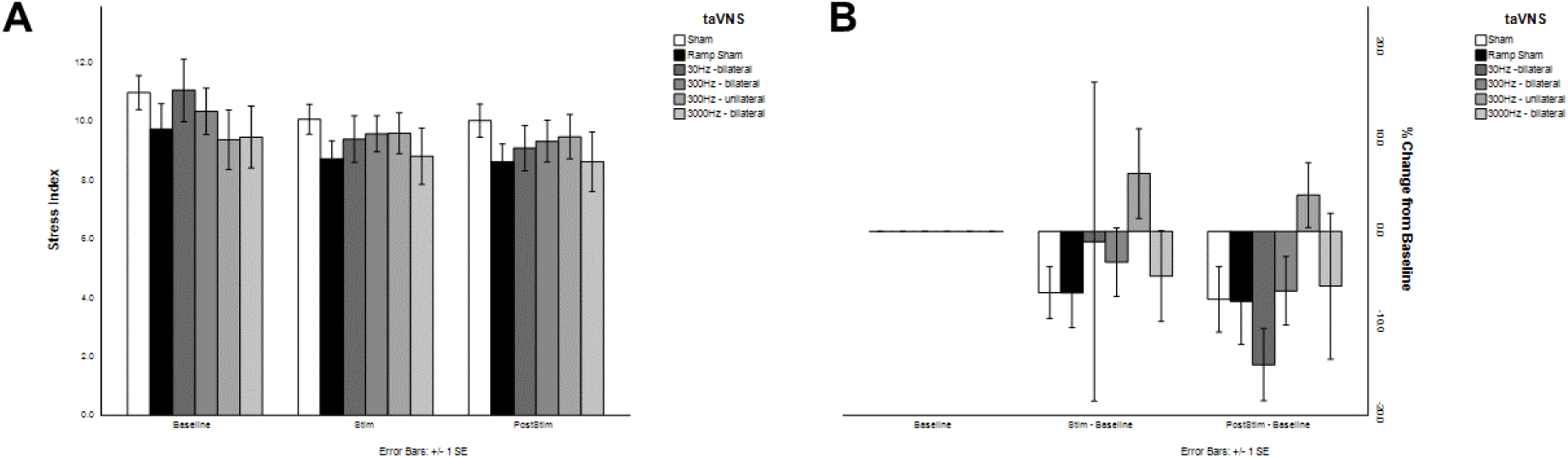
Influence of transdermal auricular vagal nerve stimulation on a Stress Index. A) Data illustrate mean Stress Index Score (square root of Baevsky’s stress index) of during 10-min baseline, taVNS stimulation, and post-stimulus periods for the inactive sham (sham), ramp sham, 30 Hz bilateral taVNS, 300 Hz bilateral taVNS, 300 Hz unilateral taVNS, and 3000 Hz taVNS. B) Data from panel A illustrated as a percent change from baseline for the taVNS and post-stimulus time periods across all taVNS treatment conditions.

**Figure 13.**
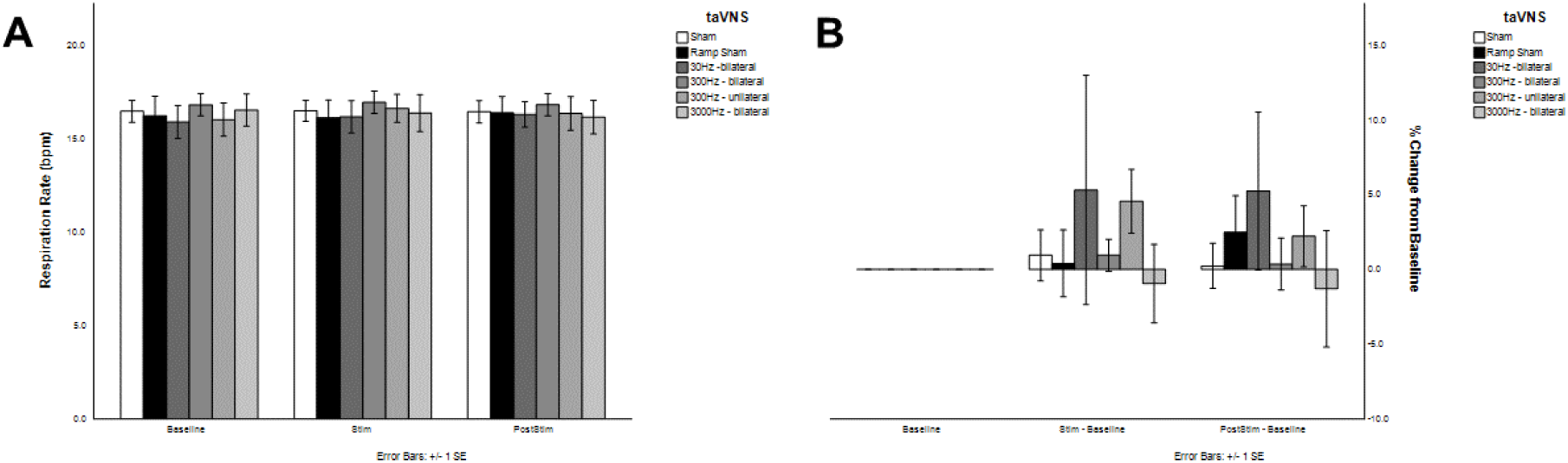
Influence of transdermal auricular vagal nerve stimulation on respiration rate. A) Data illustrate mean respiration rates in breathes per minute (bpm) during 10-min baseline, taVNS stimulation, and post-stimulus periods for the inactive sham (sham), ramp sham, 30 Hz bilateral taVNS, 300 Hz bilateral taVNS, 300 Hz unilateral taVNS, and 3000 Hz taVNS. B) Data from panel A illustrated as a percent change from baseline for the taVNS and post-stimulus time periods across all taVNS treatment conditions.

**Figure 14.**
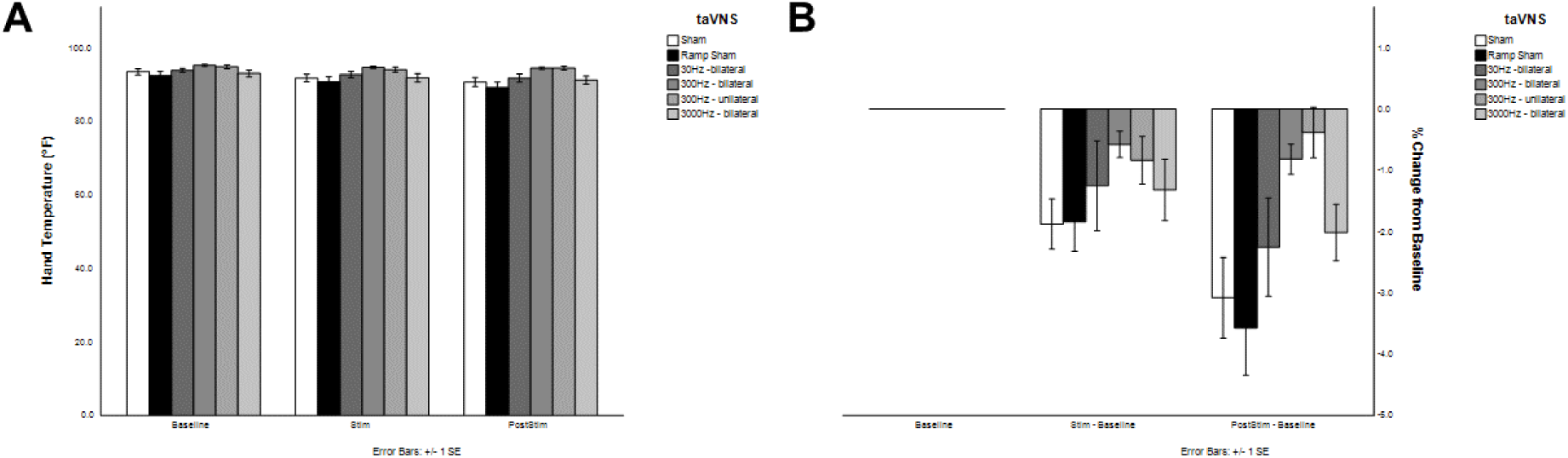
Influence of transdermal auricular vagal nerve stimulation on hand skin temperature. A) Data illustrate mean skin temperature of the hand in degrees Fahrenheit during 10-min baseline, taVNS stimulation, and post-stimulus periods for the inactive sham (sham), ramp sham, 30 Hz bilateral taVNS, 300 Hz bilateral taVNS, 300 Hz unilateral taVNS, and 3000 Hz taVNS. B) Data from panel A illustrated as a percent change from baseline for the taVNS and post-stimulus time periods across all taVNS treatment conditions.

**Figure 15.**
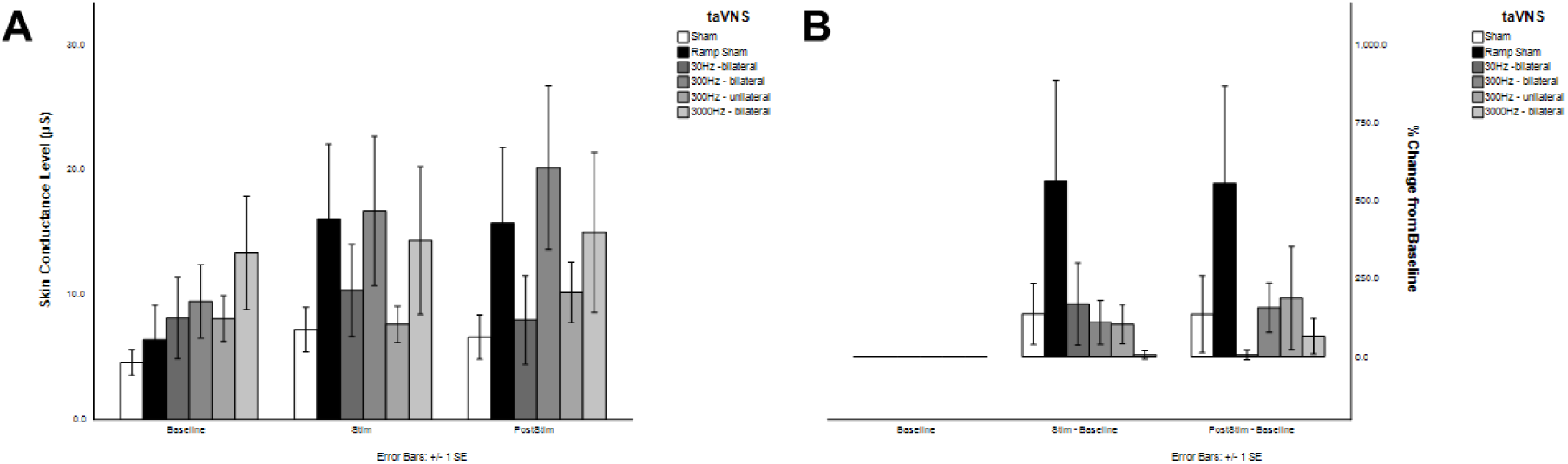
Influence of transdermal auricular vagal nerve stimulation on skin conductance. A) Data illustrate mean skin conductance in micro-Siemens during 10-min baseline, taVNS stimulation, and post-stimulus periods for the inactive sham (sham), ramp sham, 30 Hz bilateral taVNS, 300 Hz bilateral taVNS, 300 Hz unilateral taVNS, and 3000 Hz taVNS. B) Data from panel A illustrated as a percent change from baseline for the taVNS and post-stimulus time periods across all taVNS treatment conditions.

### Influence of taVNS on EEG Auditory Evoked Potentials

We investigated several measures of brain activity using EEG. We recorded EEG using a standard 10-20 montage. We recorded EEG activity in response to auditory tone presentation of frequent and infrequent pure tones. Here we analyzed the responses to frequent tones to determine the influence of taVNS on auditory evoked potentials (AEP) assayed by EEG. We examined five distinct evoked related potentials (ERP) by calculating their amplitudes and latencies across different taVNS treatment conditions. Figure 16 illustrates the basic features of AEPs we studied including the P1, N1, P2, N2, and P3 ERPs. In order to better resolve changes in brain activity we calculated the global averages across three EEG channels for each of three different scalp regions (region 1 = frontal, 2 = central, 3 = posterior) as illustrated in Figure 16. Mean peak amplitudes and latencies are shown for the EEG ERP components analyzed in Table 4.

**Figure 16.**
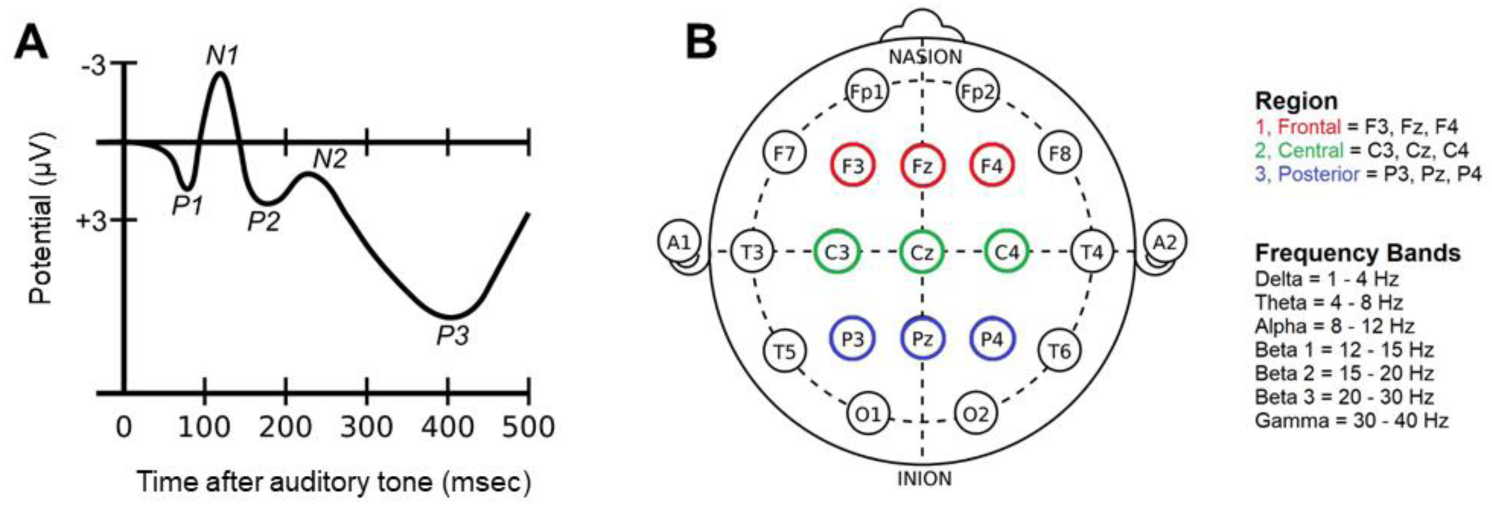
Basic methodological approach for measuring auditory evoked related potential changes produced by transdermal auricular nerve stimulation. A) Schematic of an auditory ERP is illustrated to show the P1, N1, P2, N2, and P3 components of the potentials analyzed. B) Topological map showing the 10-20 EEG montage implemented in our studies with the regions and electrodes used for calculating global averages across the scalp.

**Table 4.**
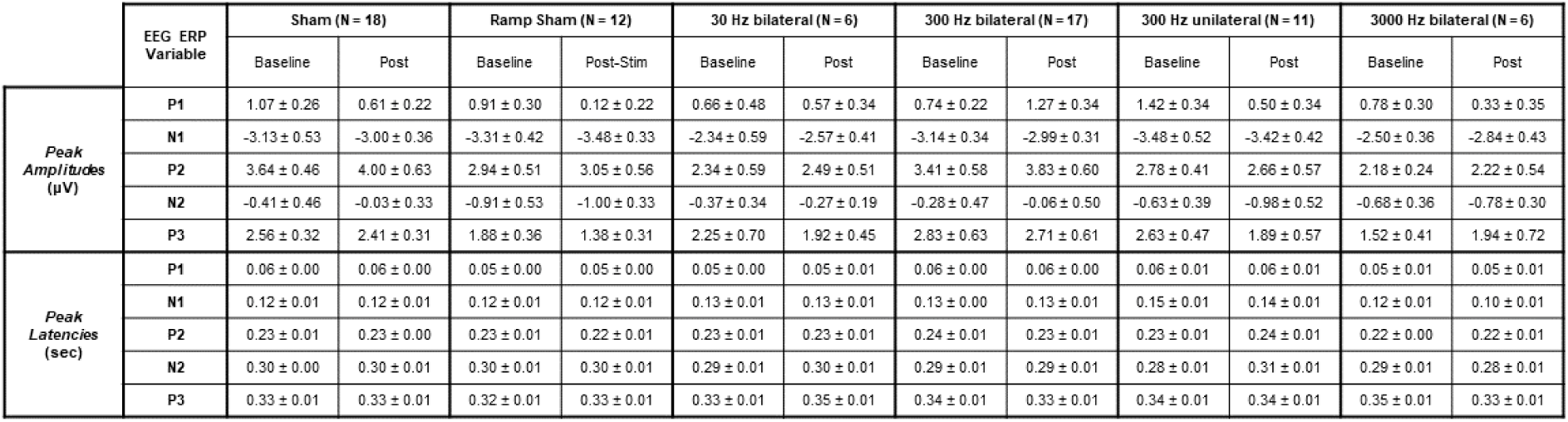
Summary of EEG AEP component amplitudes and latencies across taVNS treatment conditions.

Statistical analyses performed using Time*taVNS Frequency*Region ANOVAs revealed significant (p < 0.05) Time main effects for P1 amplitudes and N1 latencies (Table 5). Significant main effects of Frequency were also found on N1 latencies. A significant Time*taVNS Frequency interaction was found on N2 latencies and a significant Time*Frequency*Region interaction was observed on P2 latencies. No significant simple effects of taVNS Frequency were observed on ERP peak amplitudes of latencies. There were however significant simple effects of Time found for the 300 Hz unilateral (left ear) treatment condition on P1 amplitudes and N2 latencies, as well as for 3000 Hz bilateral taVNS on N1 latencies.

**Table 5.**
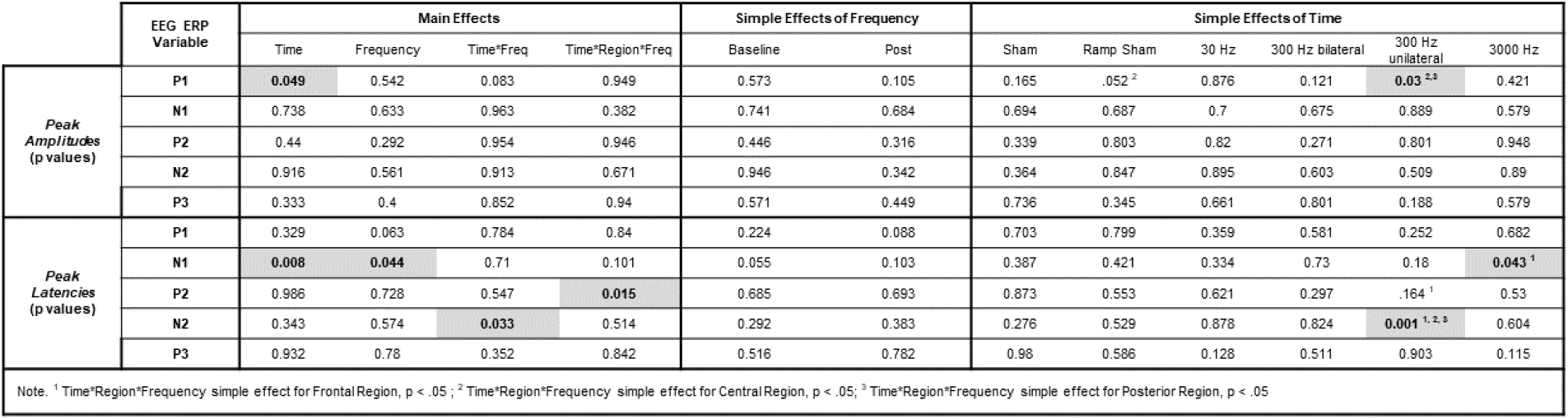
Statistical results table of P-values for influence of transdermal auricular vagal nerve stimulation on EEG auditory evoked related potential components. Statistically significant results are indicated where p < 0.05 by bold text in a grey shaded cell for the amplitudes and latencies of the P1, N1, P2, N2, and P3 auditory ERP components.

|  | EEG ERP Variable | Main Effects |  |  |  | Simple Effects of Frequency |  | Simple Effects of Time |  |  |  |  |  |
| --- | --- | --- | --- | --- | --- | --- | --- | --- | --- | --- | --- | --- | --- |
|  |  | Time | Frequency | Time*Freq | Time*Region*Freq | Baseline | Post | Sham | Ramp Sham | 30 Hz | 300 Hz bilateral | 300 Hz unilateral | 3000 Hz |
| Peak Amplitudes (p values) | P1 | **0.049** | 0.542 | 0.083 | 0.949 | 0.573 | 0.105 | 0.165 | .052<sup>2</sup> | 0.876 | 0.121 | **0.03<sup>2,3</sup>** | 0.421 |
|  | N1 | 0.738 | 0.633 | 0.983 | 0.382 | 0.741 | 0.684 | 0.694 | 0.687 | 0.7 | 0.675 | 0.889 | 0.579 |
|  | P2 | 0.44 | 0.292 | 0.954 | 0.946 | 0.446 | 0.316 | 0.339 | 0.803 | 0.82 | 0.271 | 0.801 | 0.948 |
|  | N2 | 0.916 | 0.561 | 0.913 | 0.671 | 0.946 | 0.342 | 0.364 | 0.847 | 0.895 | 0.603 | 0.509 | 0.89 |
|  | P3 | 0.333 | 0.4 | 0.852 | 0.94 | 0.571 | 0.449 | 0.736 | 0.345 | 0.661 | 0.801 | 0.188 | 0.579 |
| Peak Latencies (p values) | P1 | 0.329 | 0.063 | 0.784 | 0.84 | 0.224 | 0.088 | 0.703 | 0.799 | 0.359 | 0.581 | 0.252 | 0.682 |
|  | N1 | **0.008** | **0.044** | 0.71 | 0.101 | 0.055 | 0.103 | 0.387 | 0.421 | 0.334 | 0.73 | 0.18 | **0.043<sup>1</sup>** |
|  | P2 | 0.986 | 0.728 | 0.547 | **0.015** | 0.685 | 0.693 | 0.873 | 0.553 | 0.621 | 0.297 | .164<sup>1</sup> | 0.53 |
|  | N2 | 0.343 | 0.574 | **0.033** | 0.514 | 0.292 | 0.383 | 0.276 | 0.529 | 0.878 | 0.824 | **0.001<sup>1,2,3</sup>** | 0.604 |
|  | P3 | 0.932 | 0.78 | 0.352 | 0.842 | 0.516 | 0.782 | 0.98 | 0.586 | 0.128 | 0.511 | 0.903 | 0.115 |

Note. <sup>1</sup> Time\*Region\*Frequency simple effect for Frontal Region, $p < .05$ ; <sup>2</sup> Time\*Region\*Frequency simple effect for Central Region, $p < .05$ ; <sup>3</sup> Time\*Region\*Frequency simple effect for Posterior Region, $p < .05$

**Figure 17.**
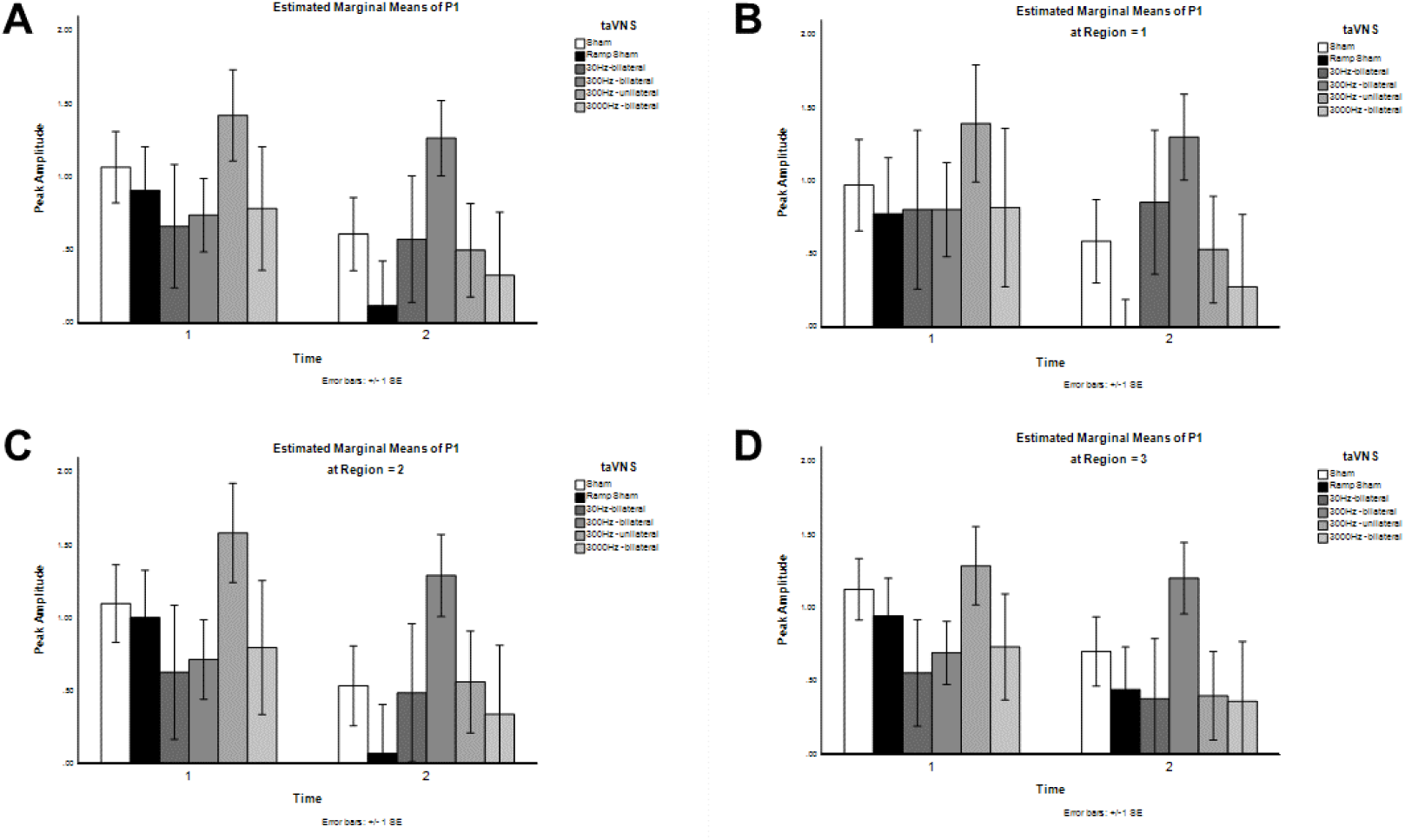
Influence of transdermal auricular vagal nerve stimulation on the amplitude of the P1 component of auditory evoked related potentials. A) The global averages (estimated marginal means) calculated from EEG channels F3, Fz, F4, C3, Cz, C4, P3, Pz, and P4 are shown for the amplitudes of the P1 ERP component as calculated for inactive sham (sham), ramp sham, 30 Hz bilateral taVNS, 300 Hz bilateral taVNS, 300 Hz unilateral (left) taVNS), and 3000 Hz taVNS treatment conditions in responses to frequent tones before (Time 1) and after taVNS (Time 2). B) The global averages calculated for region 1 from EEG channels F3, Fz, and F4 are shown for the amplitudes of the P1 ERP component. C) The global averages calculated for region 2 from EEG channels C3, Cz, and C4 are shown for the amplitudes of the P1 ERP component. D) The global averages calculated for region 3 from EEG channels P3, Pz, and P4 are shown for the amplitudes of the P1 ERP component.

**Figure 18.**
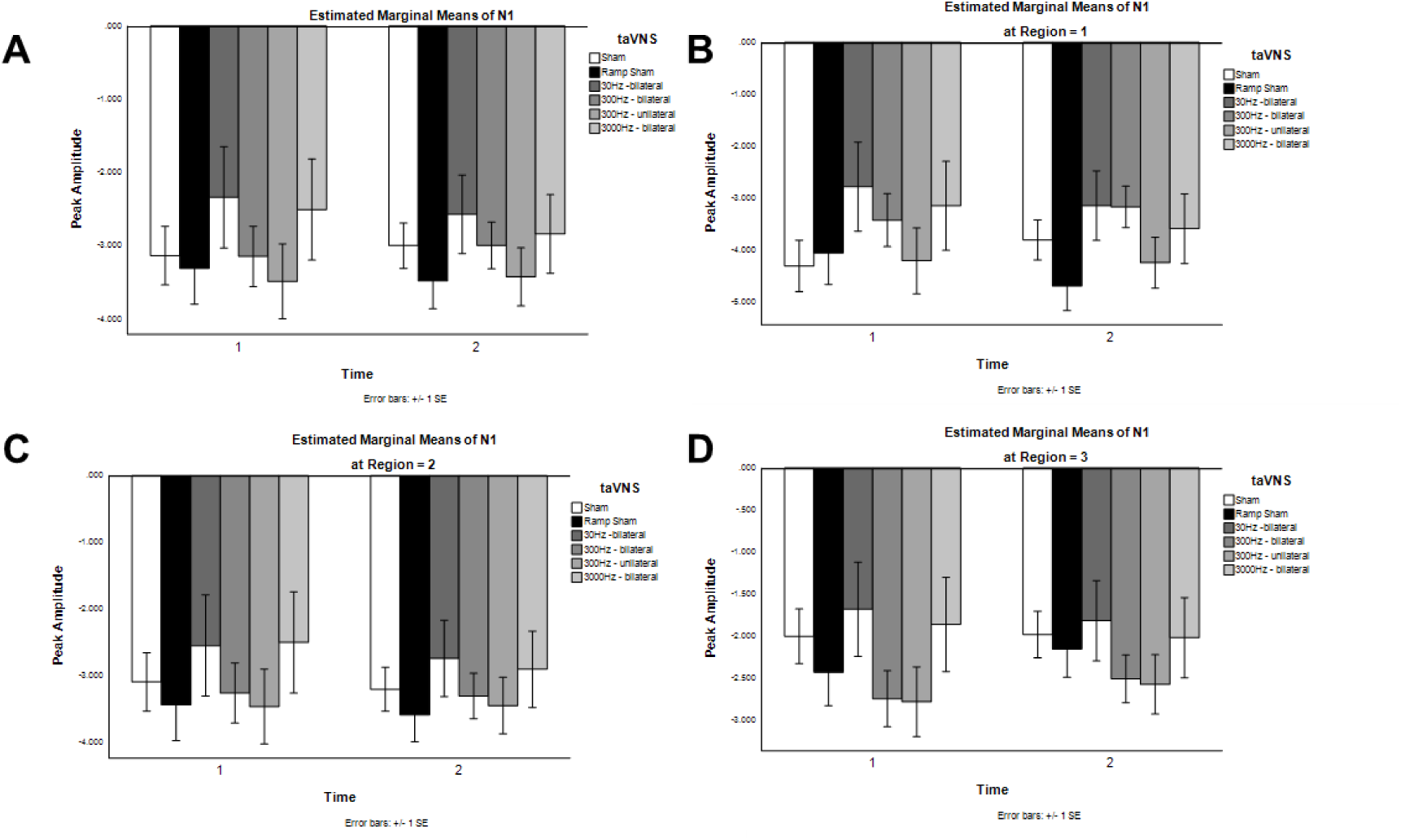
Influence of transdermal auricular vagal nerve stimulation on the amplitude of the N1 component of auditory evoked related potentials. A) The global averages calculated from EEG channels F3, Fz, F4, C3, Cz, C4, P3, Pz, and P4 are shown for the amplitudes of the N1 ERP component as calculated for inactive sham (sham), ramp sham, 30 Hz bilateral taVNS, 300 Hz bilateral taVNS, 300 Hz unilateral (left) taVNS), and 3000 Hz taVNS treatment conditions in responses to frequent tones before (Time 1) and after taVNS (Time 2). B) The global averages calculated for region 1 from EEG channels F3, Fz, and F4 are shown for the amplitudes of the N1 ERP component. C) The global averages calculated for region 2 from EEG channels C3, Cz, and C4 are shown for the amplitudes of the N1 ERP component. D) The global averages calculated for region 3 from EEG channels P3, Pz, and P4 are shown for the amplitudes of the N1 ERP component.

**Figure 19.**
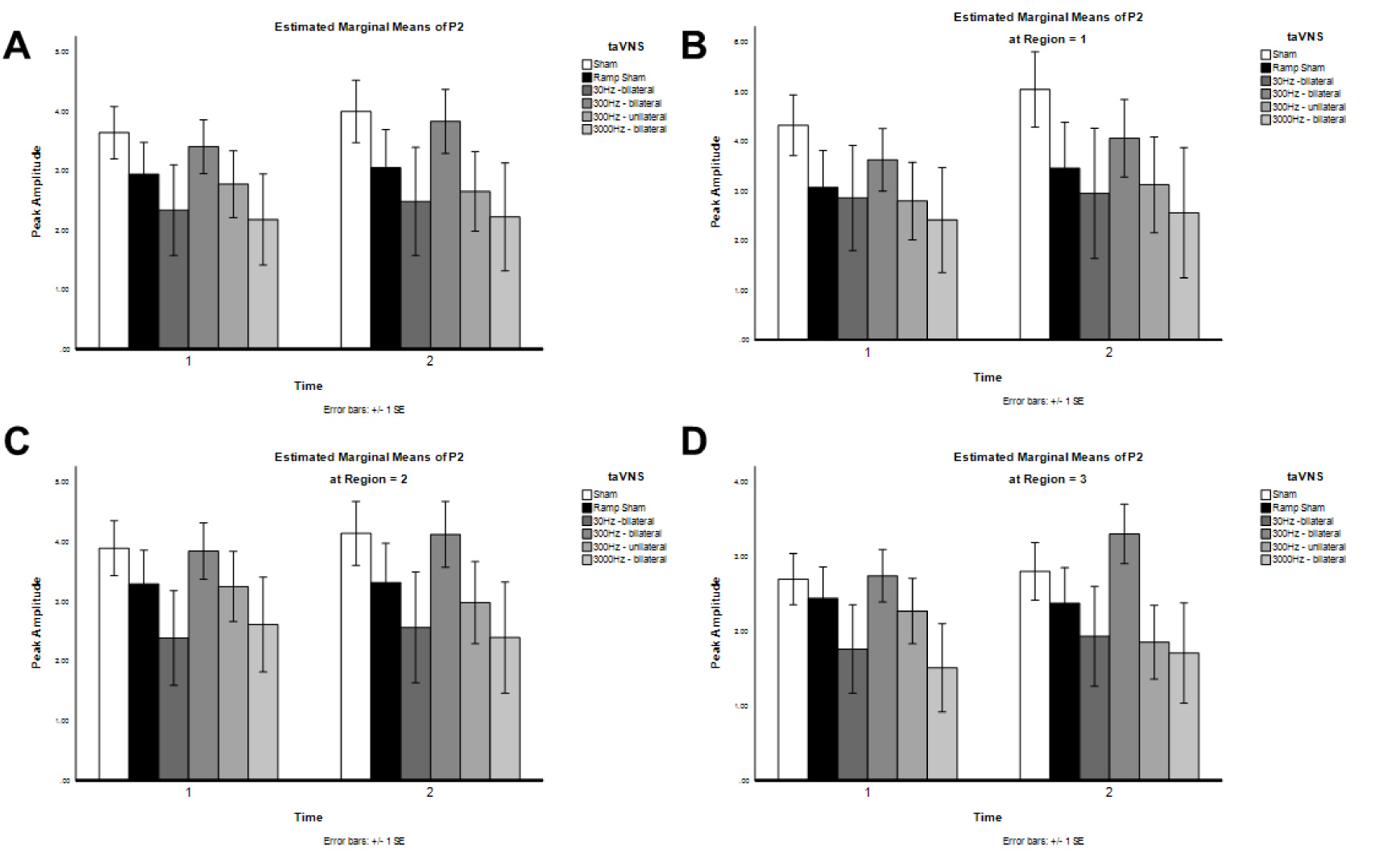
Influence of transdermal auricular vagal nerve stimulation on the amplitude of the P2 component of auditory evoked related potentials. A) The global averages calculated from EEG channels F3, Fz, F4, C3, Cz, C4, P3, Pz, and P4 are shown for the amplitudes of the P2 ERP component as calculated for inactive sham (sham), ramp sham, 30 Hz bilateral taVNS, 300 Hz bilateral taVNS, 300 Hz unilateral (left) taVNS), and 3000 Hz taVNS treatment conditions in responses to frequent tones before (Time 1) and after taVNS (Time 2). B) The global averages calculated for region 1 from EEG channels F3, Fz, and F4 are shown for the amplitudes of the P2 ERP component. C) The global averages calculated for region 2 from EEG channels C3, Cz, and C4 are shown for the amplitudes of the P2 ERP component. D) The global averages calculated for region 3 from EEG channels P3, Pz, and P4 are shown for the amplitudes of the P2 ERP component.

**Figure 20.**
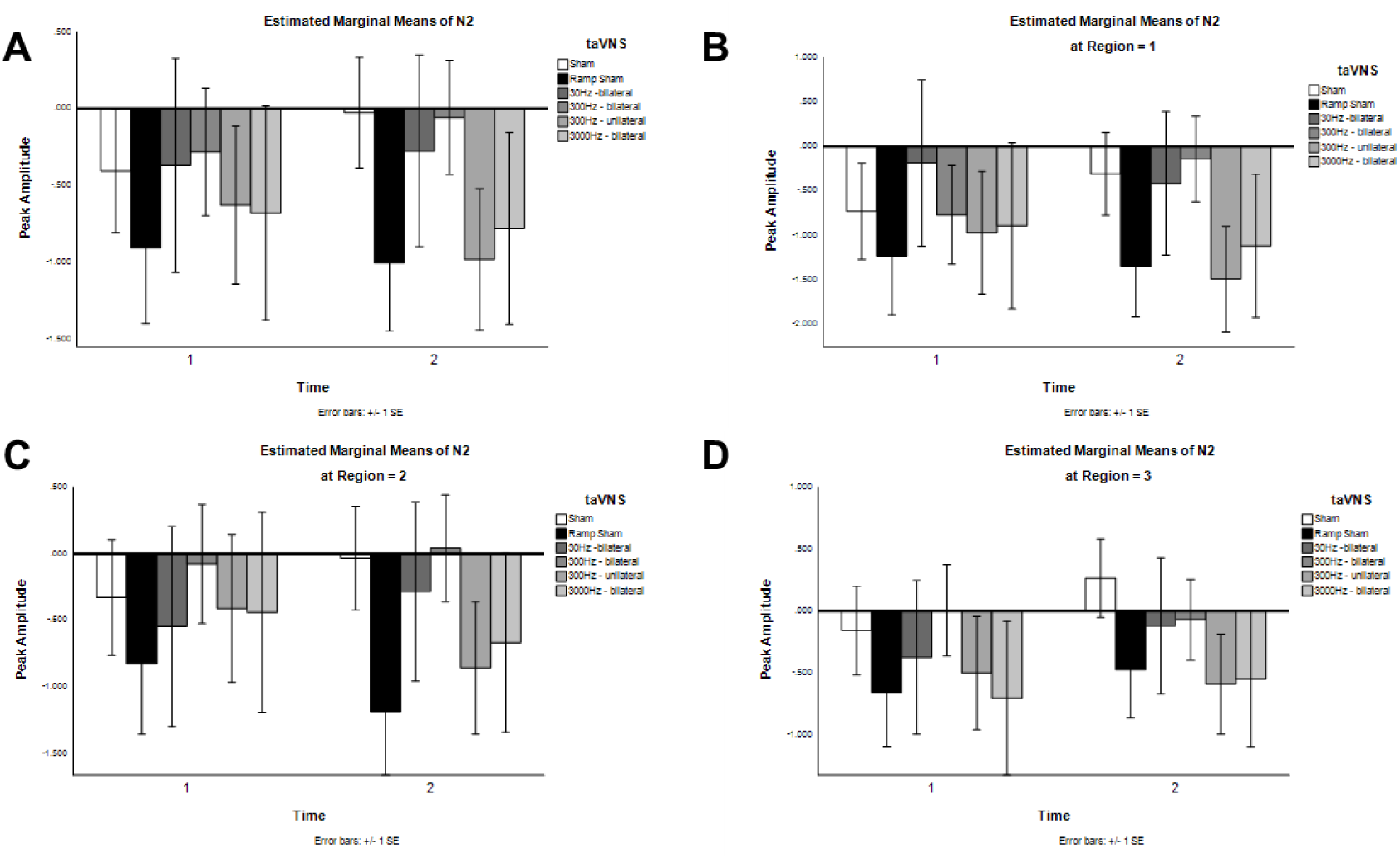
Influence of transdermal auricular vagal nerve stimulation on the amplitude of the N2 component of auditory evoked related potentials. A) The global averages calculated from EEG channels F3, Fz, F4, C3, Cz, C4, P3, Pz, and P4 are shown for the amplitudes of the N2 ERP component as calculated for inactive sham (sham), ramp sham, 30 Hz bilateral taVNS, 300 Hz bilateral taVNS, 300 Hz unilateral (left) taVNS), and 3000 Hz taVNS treatment conditions in responses to frequent tones before (Time 1) and after taVNS (Time 2). B) The global averages calculated for region 1 from EEG channels F3, Fz, and F4 are shown for the amplitudes of the N2 ERP component. C) The global averages calculated for region 2 from EEG channels C3, Cz, and C4 are shown for the amplitudes of the N2 ERP component. D) The global averages calculated for region 3 from EEG channels P3, Pz, and P4 are shown for the amplitudes of the N2 ERP component.

**Figure 21.**
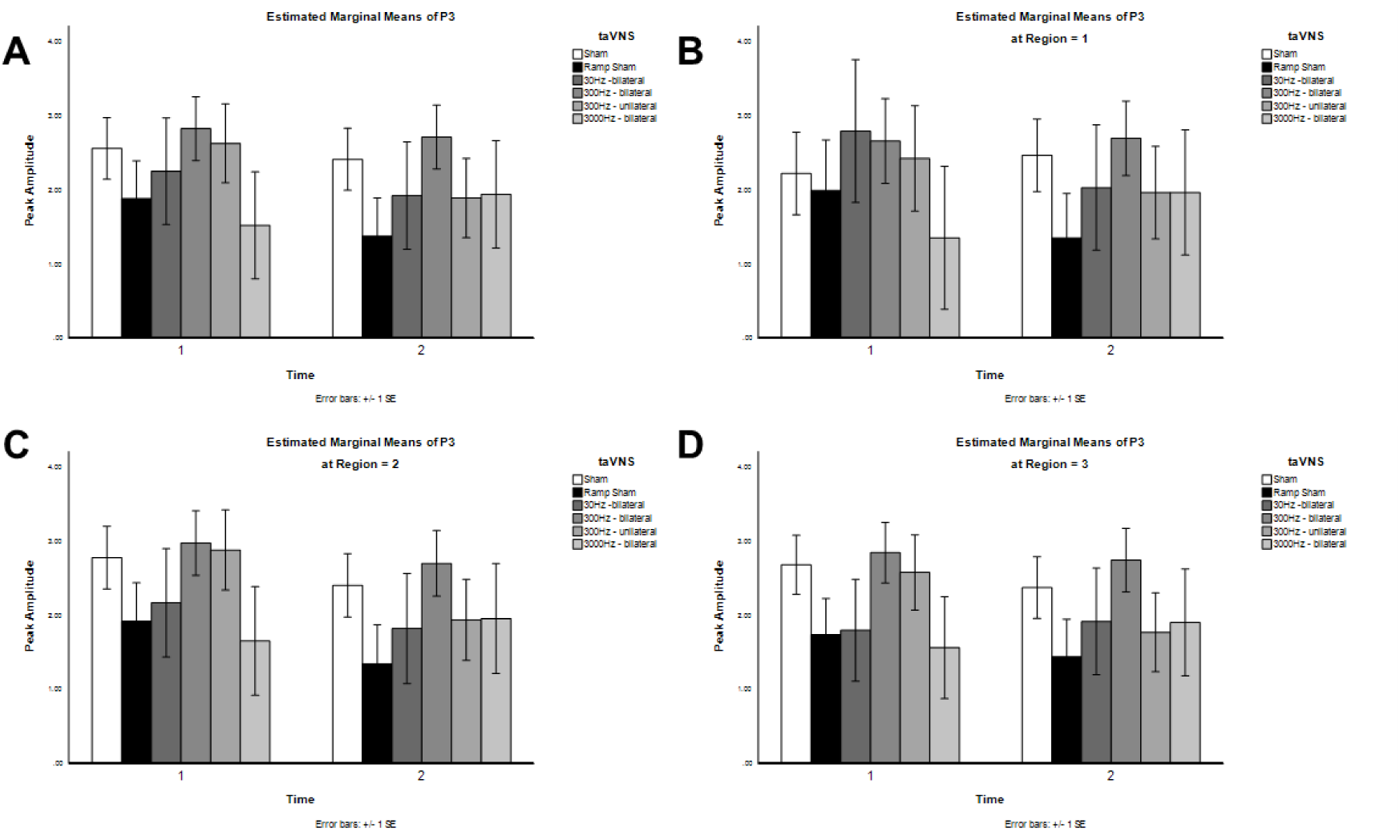
Influence of transdermal auricular vagal nerve stimulation on the amplitude of the P3 component of auditory evoked related potentials. A) The global averages calculated from EEG channels F3, Fz, F4, C3, Cz, C4, P3, Pz, and P4 are shown for the amplitudes of the P3 ERP component as calculated for inactive sham (sham), ramp sham, 30 Hz bilateral taVNS, 300 Hz bilateral taVNS, 300 Hz unilateral (left) taVNS), and 3000 Hz taVNS treatment conditions in responses to frequent tones before (Time 1) and after taVNS (Time 2). B) The global averages calculated for region 1 from EEG channels F3, Fz, and F4 are shown for the amplitudes of the P3 ERP component. C) The global averages calculated for region 2 from EEG channels C3, Cz, and C4 are shown for the amplitudes of the P3 ERP component. D) The global averages calculated for region 3 from EEG channels P3, Pz, and P4 are shown for the amplitudes of the P3 ERP component.

**Figure 22.**
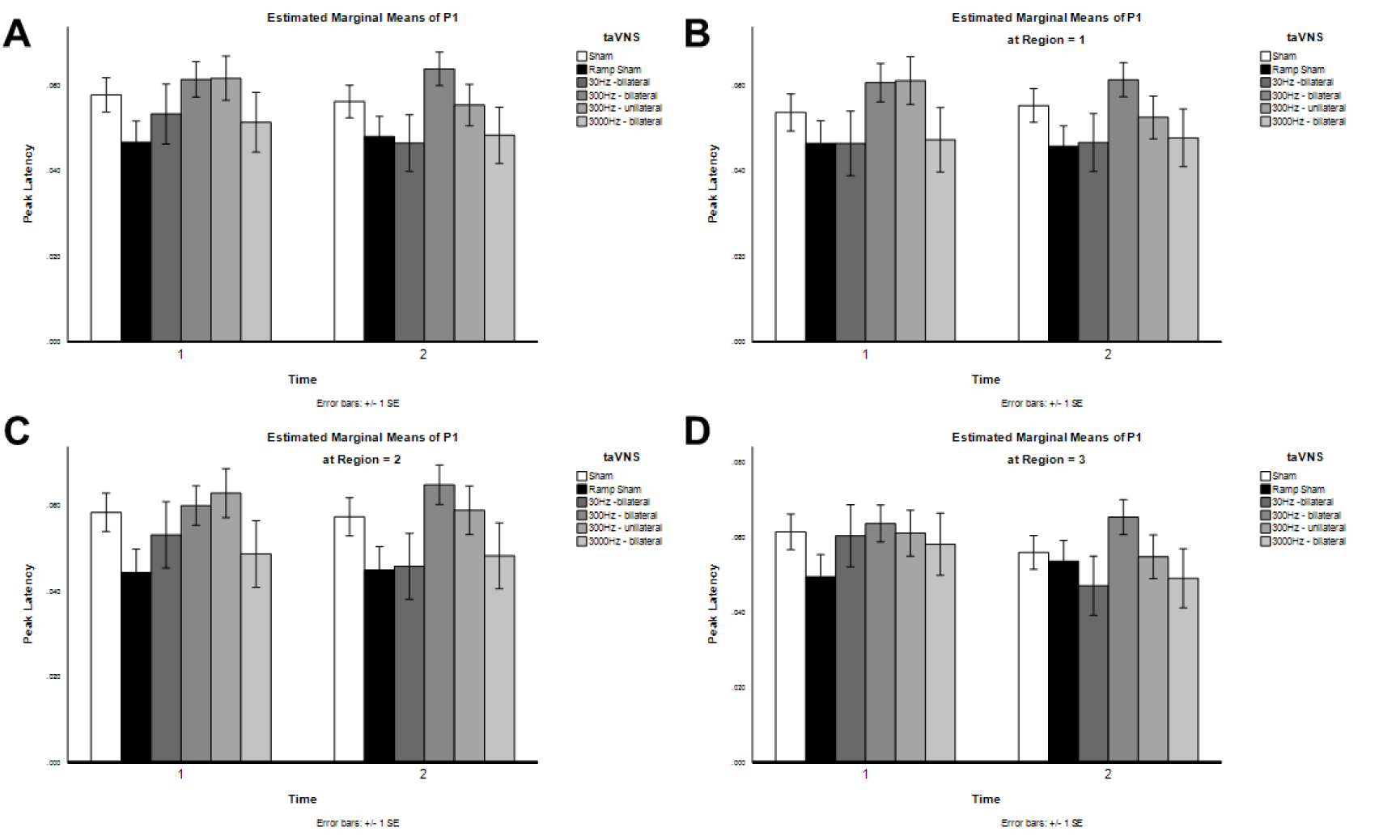
Influence of transdermal auricular vagal nerve stimulation on the latency of the P1 component of auditory evoked related potentials. A) The global averages calculated from EEG channels F3, Fz, F4, C3, Cz, C4, P3, Pz, and P4 are shown for the latencies of the P1 ERP component as calculated for inactive sham (sham), ramp sham, 30 Hz bilateral taVNS, 300 Hz bilateral taVNS, 300 Hz unilateral (left) taVNS), and 3000 Hz taVNS treatment conditions in responses to frequent tones before (Time 1) and after taVNS (Time 2). B) The global peak latency averages calculated for region 1 from EEG channels F3, Fz, and F4 are shown for the latencies of the P1 ERP component. C) The global averages calculated for region 2 from EEG channels C3, Cz, and C4 are shown for the latencies of the P1 ERP component. D) The global averages calculated for region 3 from EEG channels P3, Pz, and P4 are shown for the latencies of the P1 ERP component.

**Figure 23.**
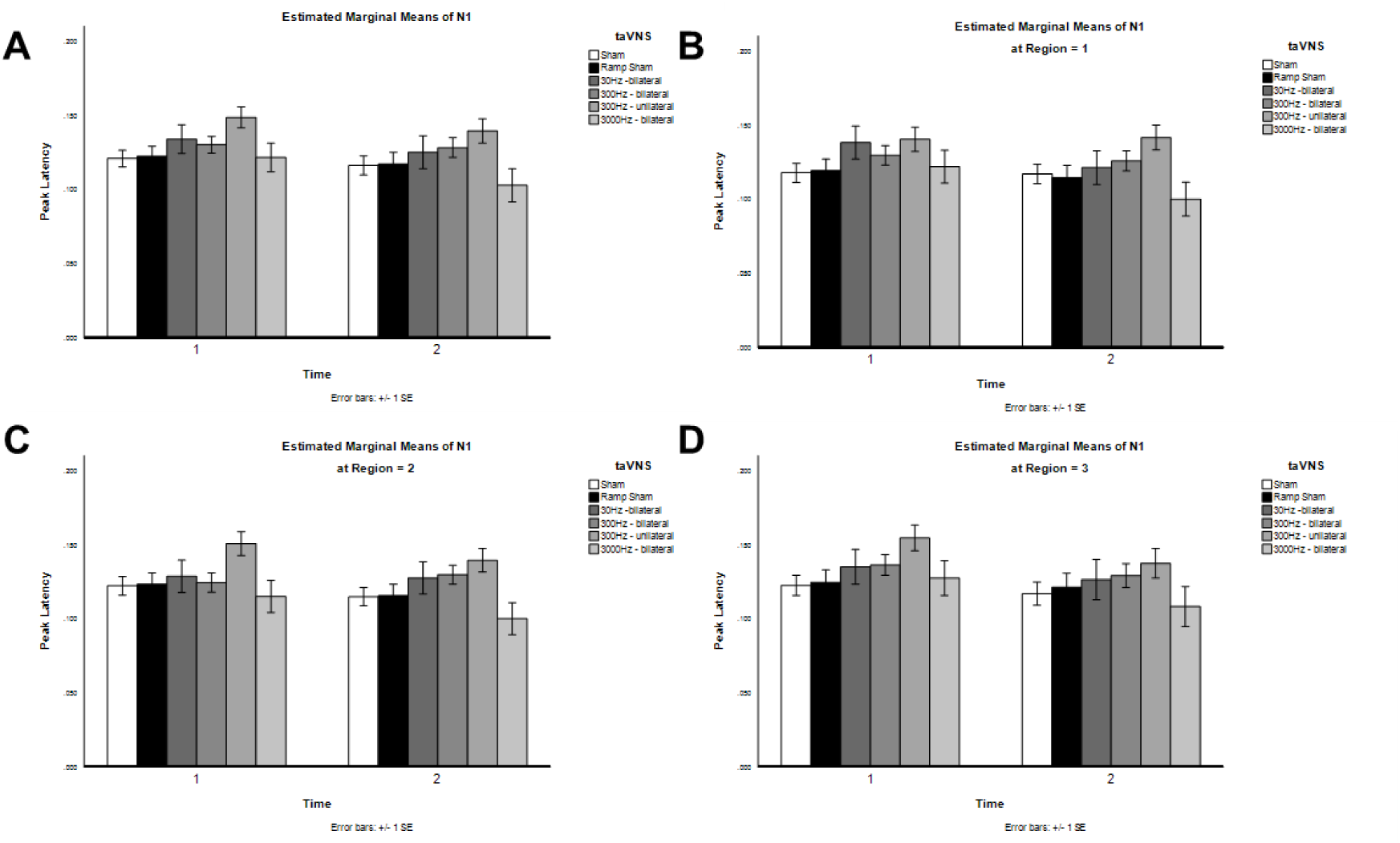
Influence of transdermal auricular vagal nerve stimulation on the latency of the N1 component of auditory evoked related potentials. A) The global averages calculated from EEG channels F3, Fz, F4, C3, Cz, C4, P3, Pz, and P4 are shown for the latencies of the N1 ERP component as calculated for inactive sham (sham), ramp sham, 30 Hz bilateral taVNS, 300 Hz bilateral taVNS, 300 Hz unilateral (left) taVNS), and 3000 Hz taVNS treatment conditions in responses to frequent tones before (Time 1) and after taVNS (Time 2). B) The global peak latency averages calculated for region 1 from EEG channels F3, Fz, and F4 are shown for the latencies of the N1 ERP component. C) The global averages calculated for region 2 from EEG channels C3, Cz, and C4 are shown for the latencies of the N1 ERP component. D) The global averages calculated for region 3 from EEG channels P3, Pz, and P4 are shown for the latencies of the N1 ERP component.

**Figure 24.**
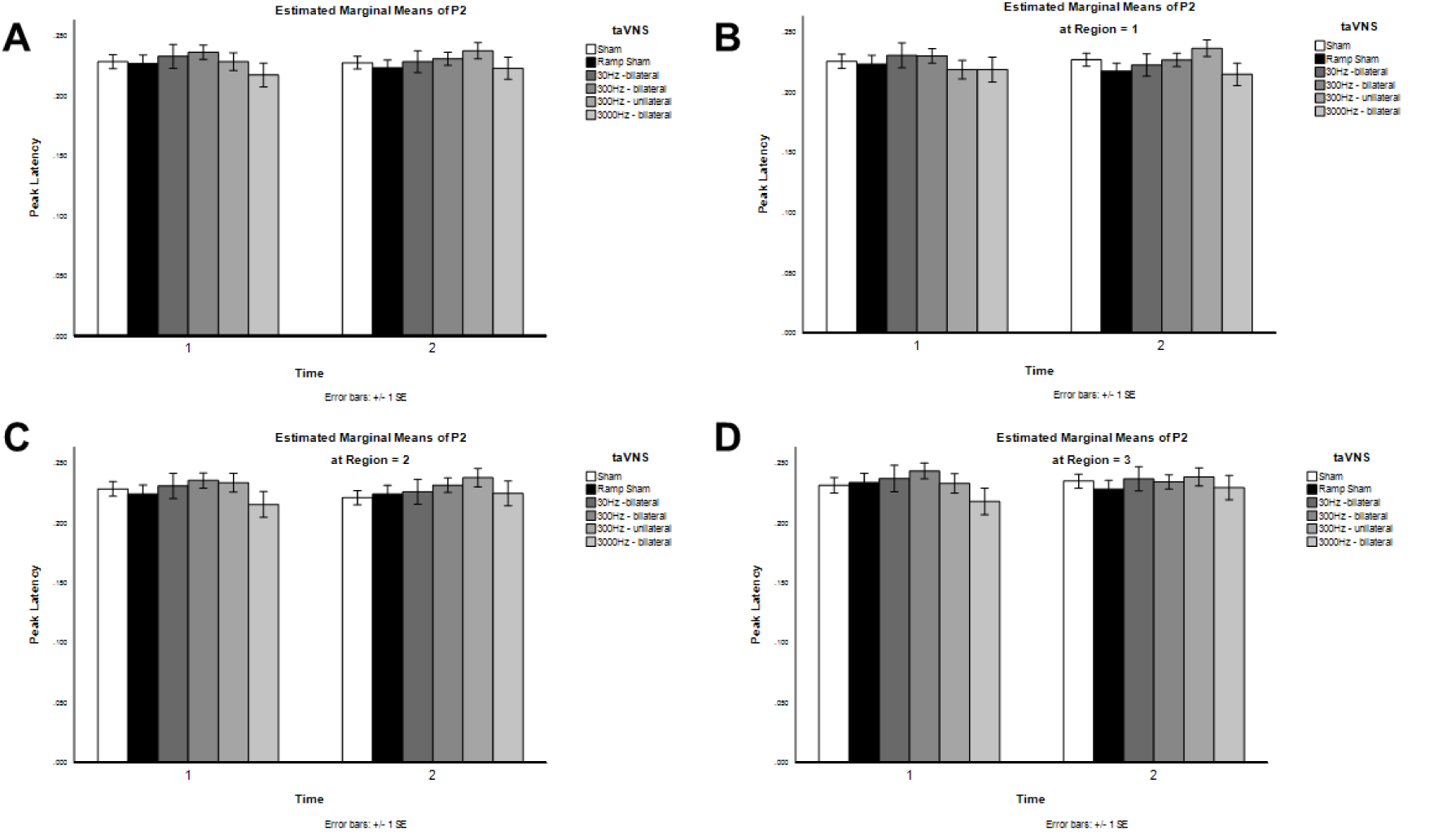
Influence of transdermal auricular vagal nerve stimulation on the latency of the P2 component of auditory evoked related potentials. A) The global averages calculated from EEG channels F3, Fz, F4, C3, Cz, C4, P3, Pz, and P4 are shown for the latencies of the P2 ERP component as calculated for inactive sham (sham), ramp sham, 30 Hz bilateral taVNS, 300 Hz bilateral taVNS, 300 Hz unilateral (left) taVNS), and 3000 Hz taVNS treatment conditions in responses to frequent tones before (Time 1) and after taVNS (Time 2). B) The global peak latency averages calculated for region 1 from EEG channels F3, Fz, and F4 are shown for the latencies of the P2 ERP component. C) The global averages calculated for region 2 from EEG channels C3, Cz, and C4 are shown for the latencies of the P2 ERP component. D) The global averages calculated for region 3 from EEG channels P3, Pz, and P4 are shown for the latencies of the P2 ERP component.

**Figure 25.**
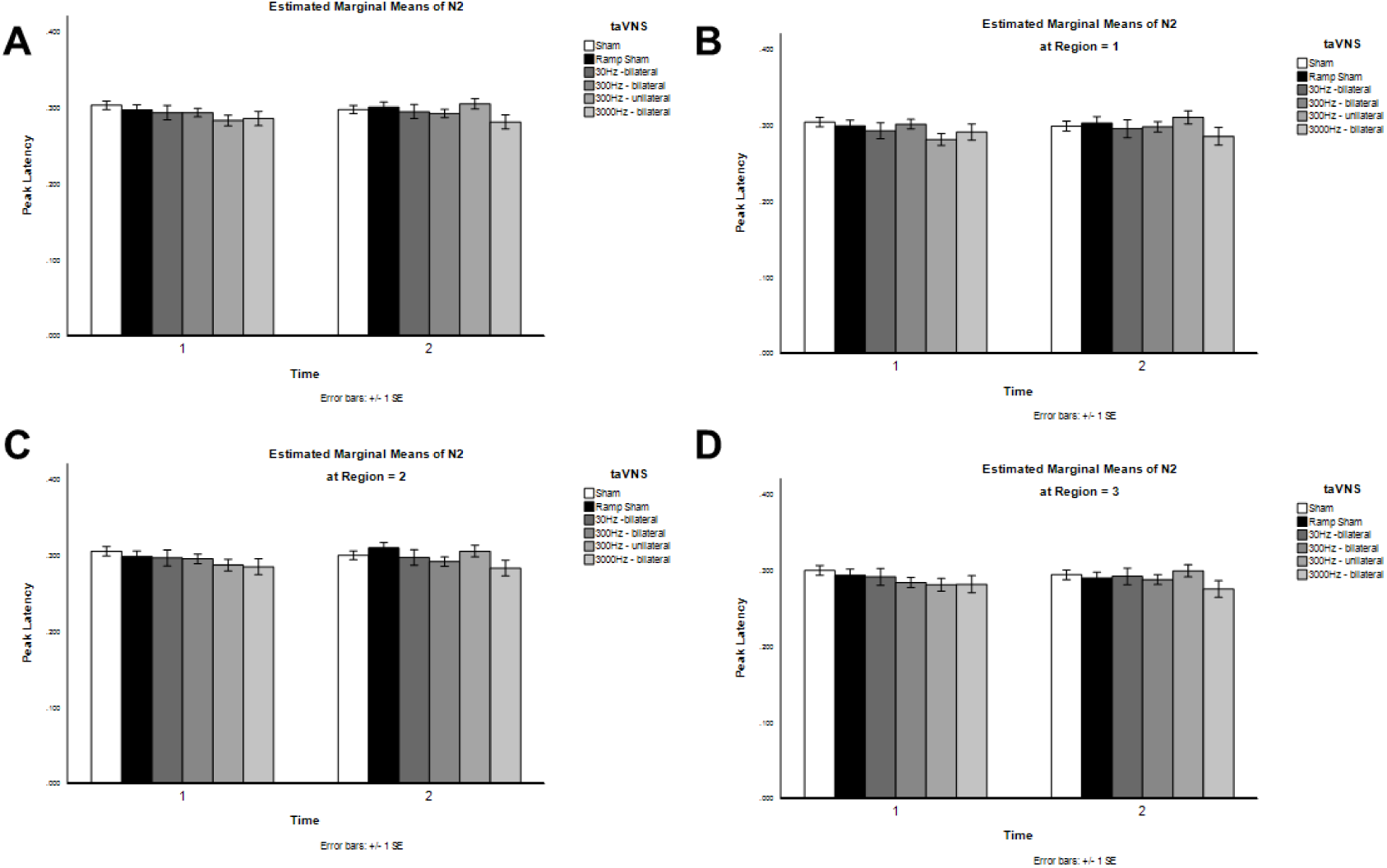
Influence of transdermal auricular vagal nerve stimulation on the latency of the N2 component of auditory evoked related potentials. A) The global averages calculated from EEG channels F3, Fz, F4, C3, Cz, C4, P3, Pz, and P4 are shown for the latencies of the N2 ERP component as calculated for inactive sham (sham), ramp sham, 30 Hz bilateral taVNS, 300 Hz bilateral taVNS, 300 Hz unilateral (left) taVNS), and 3000 Hz taVNS treatment conditions in responses to frequent tones before (Time 1) and after taVNS (Time 2). B) The global peak latency averages calculated for region 1 from EEG channels F3, Fz, and F4 are shown for the latencies of the N2 ERP component. C) The global averages calculated for region 2 from EEG channels C3, Cz, and C4 are shown for the latencies of the N2 ERP component. D) The global averages calculated for region 3 from EEG channels P3, Pz, and P4 are shown for the latencies of the N2 ERP component.

**Figure 26.**
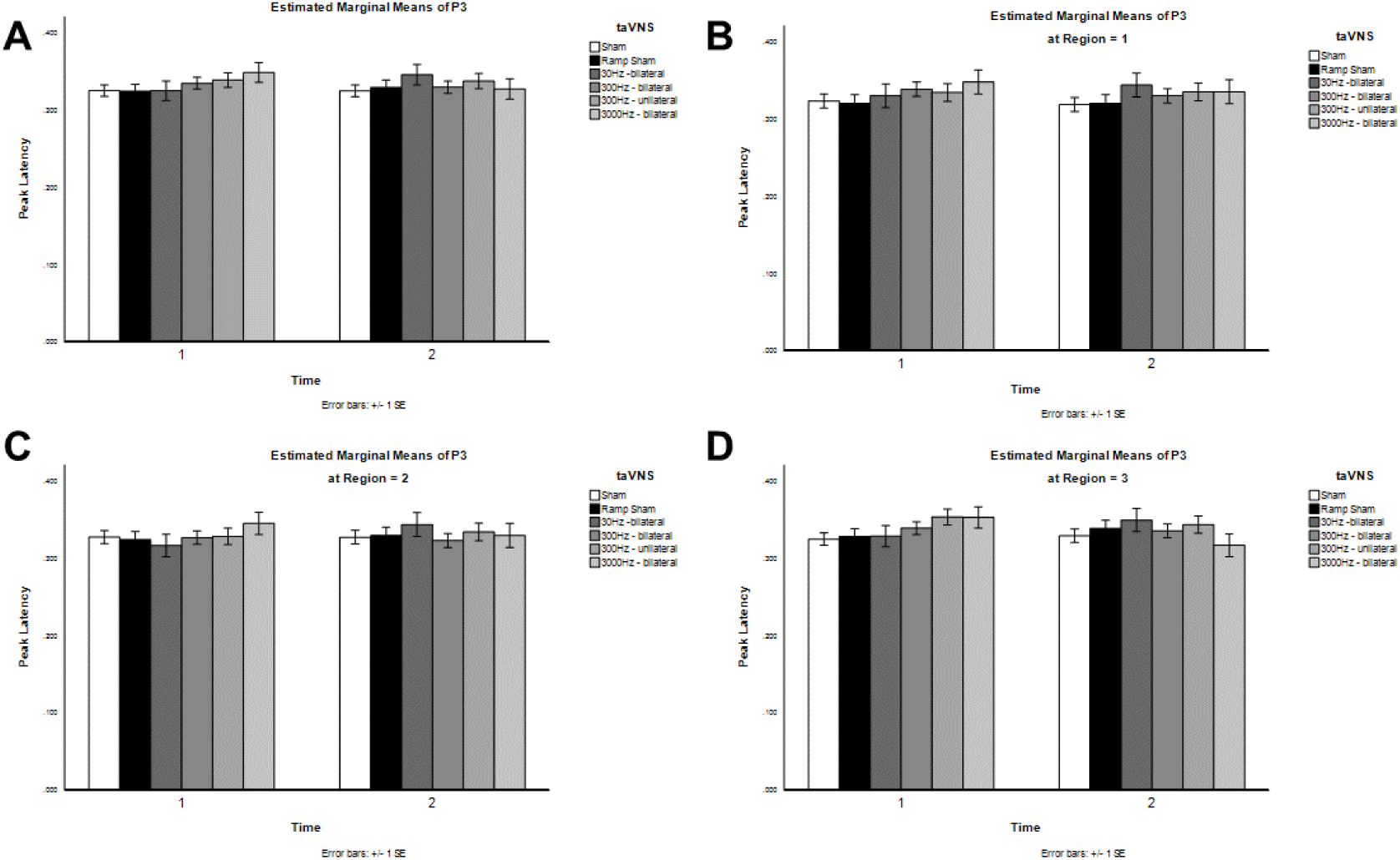
Influence of transdermal auricular vagal nerve stimulation on the latency of the P3 component of auditory evoked related potentials. A) The global averages calculated from EEG channels F3, Fz, F4, C3, Cz, C4, P3, Pz, and P4 are shown for the latencies of the P3 ERP component as calculated for inactive sham (sham), ramp sham, 30 Hz bilateral taVNS, 300 Hz bilateral taVNS, 300 Hz unilateral (left) taVNS), and 3000 Hz taVNS treatment conditions in responses to frequent tones before (Time 1) and after taVNS (Time 2). B) The global peak latency averages calculated for region 1 from EEG channels F3, Fz, and F4 are shown for the latencies of the P3 ERP component. C) The global averages calculated for region 2 from EEG channels C3, Cz, and C4 are shown for the latencies of the P3 ERP component. D) The global averages calculated for region 3 from EEG channels P3, Pz, and P4 are shown for the latencies of the P3 ERP component.

### Influence of taVNS on Continuous EEG Spectral Power

In addition to analyzing the conventional peak amplitude and latency properties of auditory ERP components, we also probed for power changes in the continuous EEG frequency spectrum spanning delta to gamma band activity. We analyzed the continuous EEG spectral data collected globally from channels F3, Fz, F4, C3, Cz, C4, P3, Pz, and P4 during passive auditory tasks before and following taVNS treatment. For each of the treatment conditions evaluated, we investigated changes in the absolute and relative power of the following EEG spectral bands: delta (1 – 4 Hz), theta (4 – 8 Hz), alpha (8 – 12 Hz), beta 1 (12 – 15 Hz), beta 2 (15 – 20 Hz), beta 3 (20 – 30 Hz) and gamma (30 – 40 Hz). Data from these investigations are summarized in Tables 6 and 7 and illustrated in Figures 27-32.

**Table 6.**
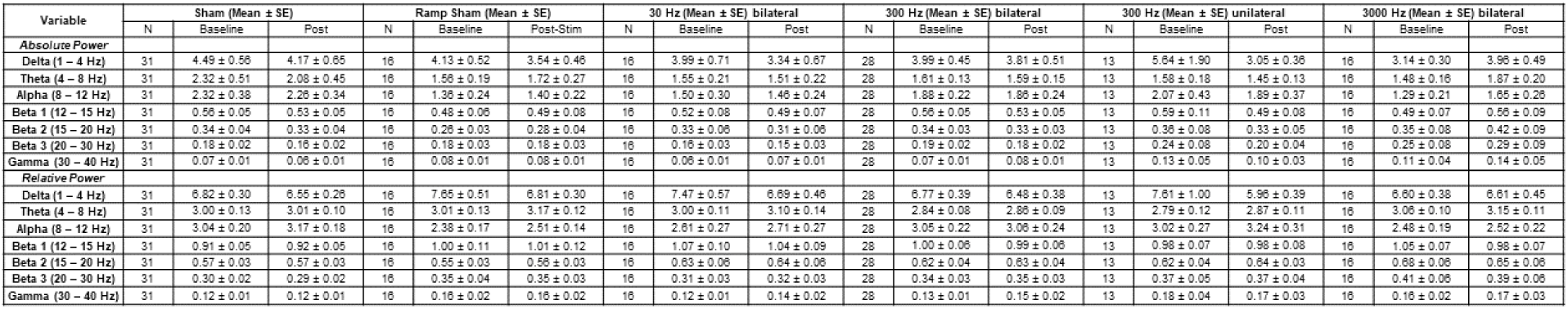
Summary of the changes in the absolute and relative powers for EEG spectral bands in response to transdermal auricular vagal nerve stimulation.

**Table 7.**
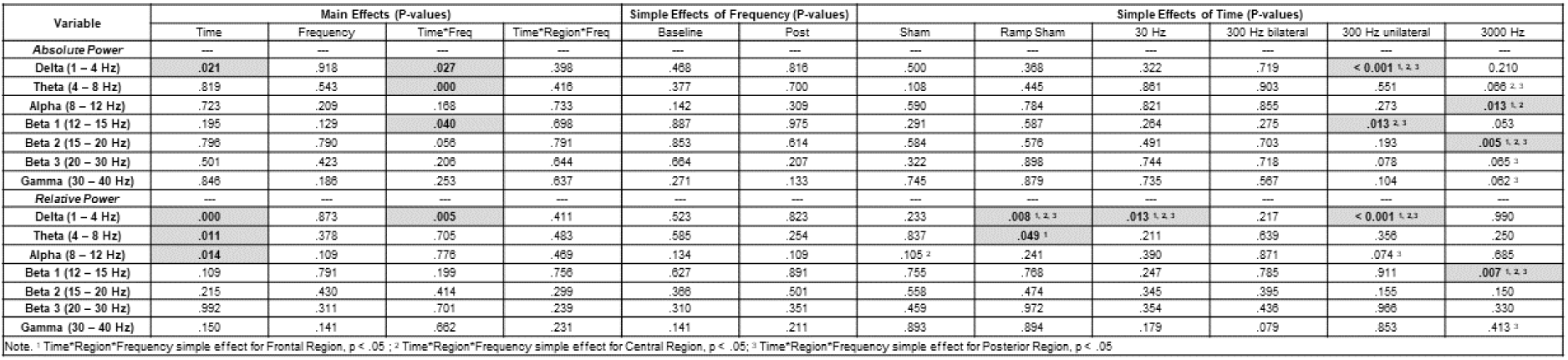
Statistical results table of P-values for influence of transdermal auricular vagal nerve stimulation on continuous EEG spectral activity. Statistically significant results are indicated where p < 0.05 by bold text in a grey shaded cell for the absolute and relative powers of delta, theta, alpha, beta 1-3, and gamma EEG frequency bands.

**Figure 27.**
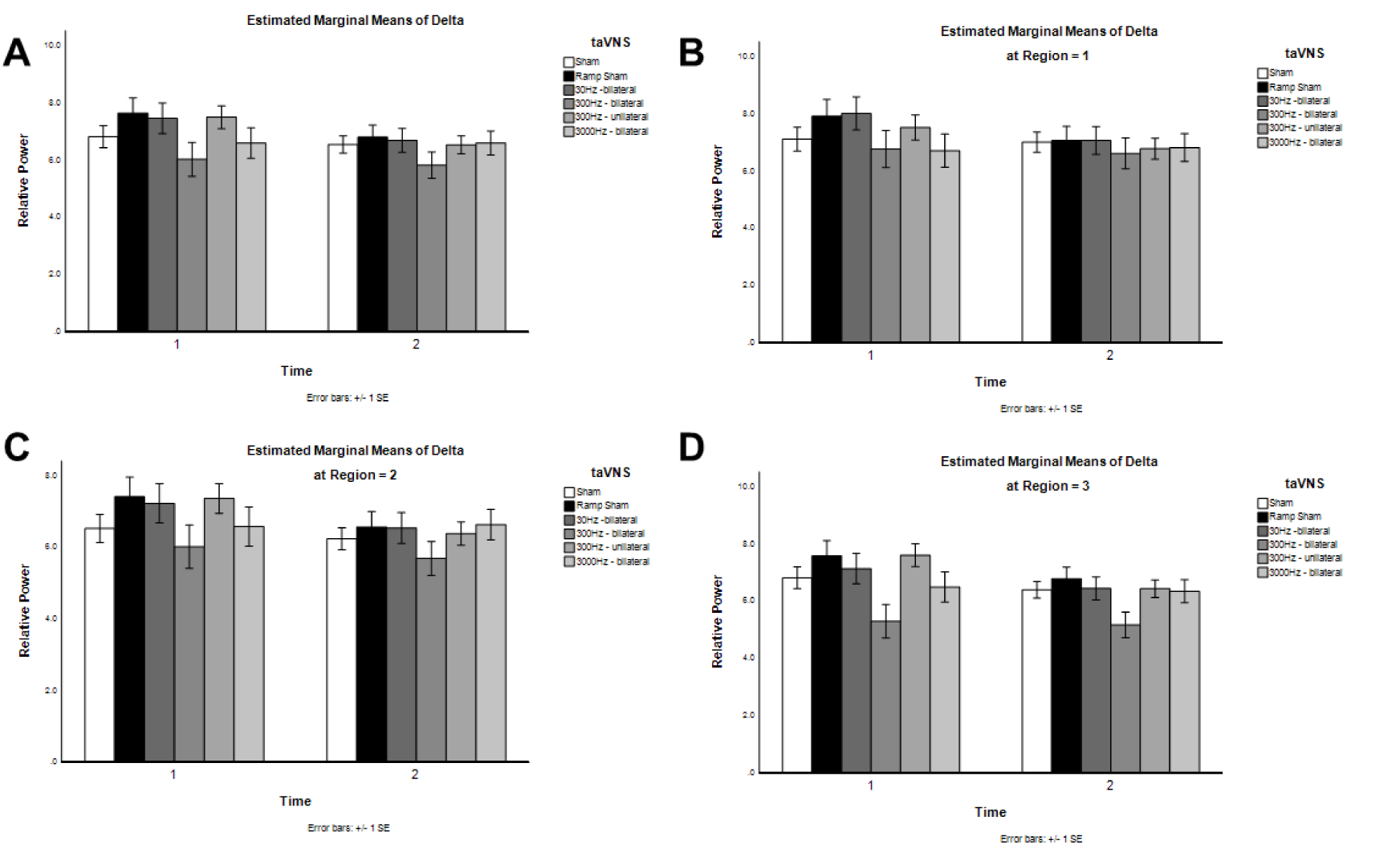
Influence of transdermal auricular vagal nerve stimulation on delta EEG frequency band activity. **A**) The mean relative powers of delta (1 – 4 Hz) frequency band activity calculated from EEG channels F3, Fz, F4, C3, Cz, C4, P3, Pz, and P4 are shown as calculated for inactive sham (sham), ramp sham, 30 Hz bilateral taVNS, 300 Hz bilateral taVNS, 300 Hz unilateral (left) taVNS), and 3000 Hz taVNS treatment conditions in responses to frequent tones before (Time 1) and after taVNS (Time 2). **B**) Mean relative powers of delta frequency band activity calculated for region 1 (frontal) from EEG channels F3, Fz, and F4 are shown. **C**) Mean relative powers of delta frequency band activity calculated for region 2 (central) from EEG channels C3, Cz, and C4 are shown. **D**) Mean relative powers of delta frequency band activity calculated for region 3 (parietal) from EEG channels P3, Pz, and P4 are shown.

**Figure 28.**
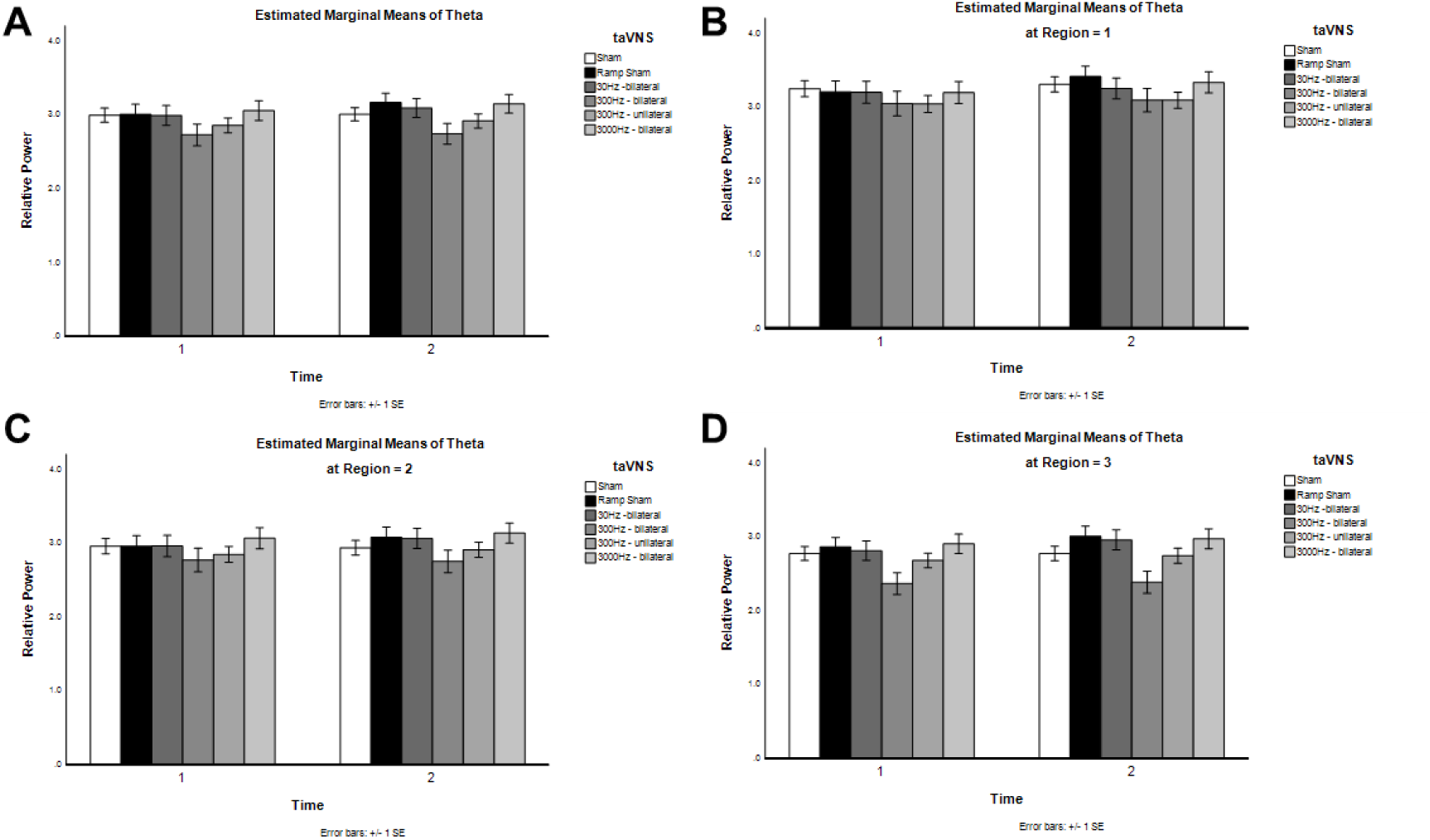
Influence of transdermal auricular vagal nerve stimulation on theta EEG frequency band activity. **A**) The mean relative powers of theta (4 – 8 Hz) frequency band activity calculated from EEG channels F3, Fz, F4, C3, Cz, C4, P3, Pz, and P4 are shown as calculated for inactive sham (sham), ramp sham, 30 Hz bilateral taVNS, 300 Hz bilateral taVNS, 300 Hz unilateral (left) taVNS), and 3000 Hz taVNS treatment conditions in responses to frequent tones before (Time 1) and after taVNS (Time 2). **B**) Mean relative powers of theta frequency band activity calculated for region 1 (frontal) from EEG channels F3, Fz, and F4 are shown. **C**) Mean relative powers of theta frequency band activity calculated for region 2 (central) from EEG channels C3, Cz, and C4 are shown. **D**) Mean relative powers of theta frequency band activity calculated for region 3 (parietal) from EEG channels P3, Pz, and P4 are shown.

**Figure 29.**
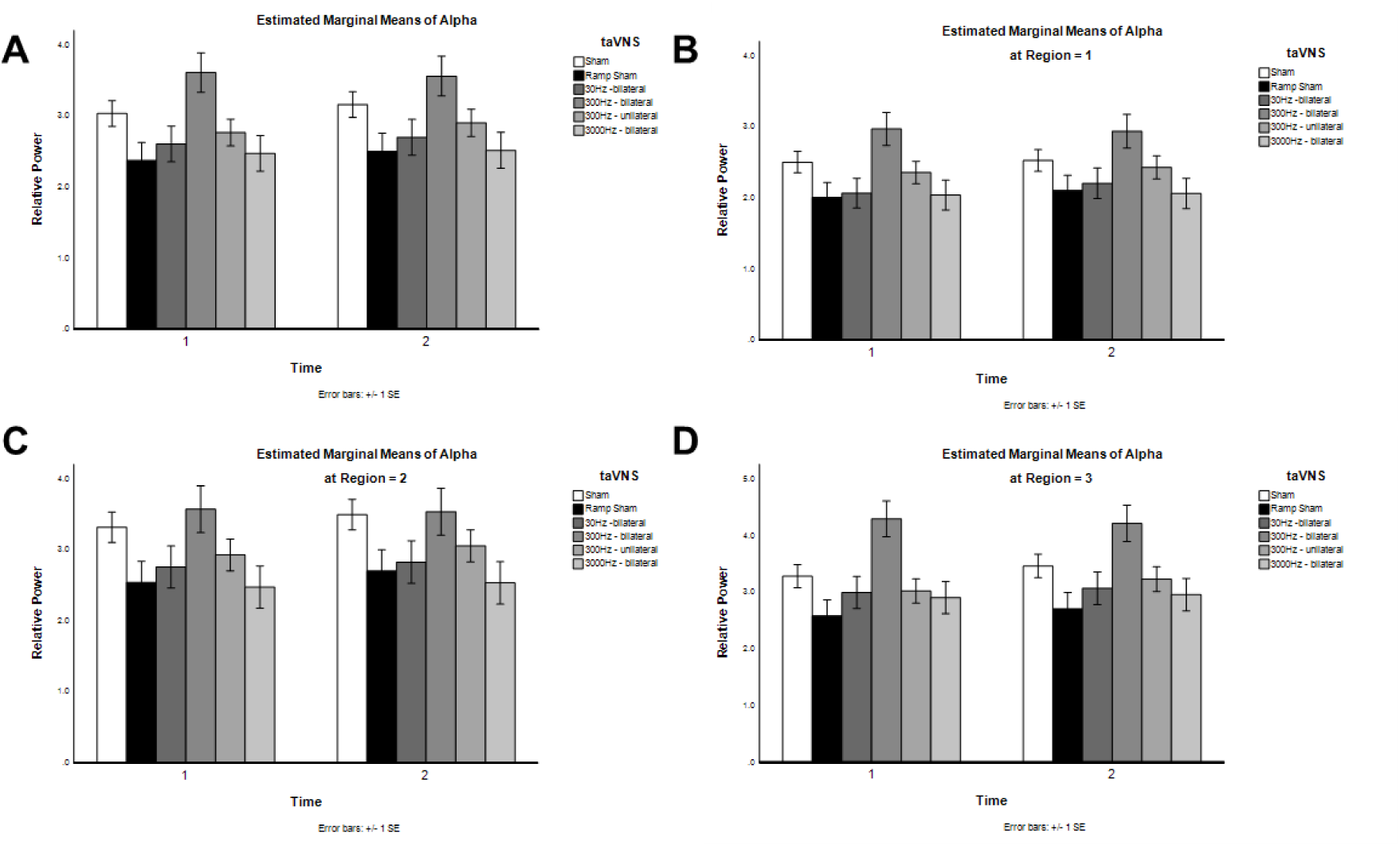
Influence of transdermal auricular vagal nerve stimulation on alpha EEG frequency band activity. **A**) The mean relative powers of alpha (8 – 12 Hz) frequency band activity calculated from EEG channels F3, Fz, F4, C3, Cz, C4, P3, Pz, and P4 are shown as calculated for inactive sham (sham), ramp sham, 30 Hz bilateral taVNS, 300 Hz bilateral taVNS, 300 Hz unilateral (left) taVNS), and 3000 Hz taVNS treatment conditions in responses to frequent tones before (Time 1) and after taVNS (Time 2). **B**) Mean relative powers of alpha frequency band activity calculated for region 1 (frontal) from EEG channels F3, Fz, and F4 are shown. **C**) Mean relative powers of alpha frequency band activity calculated for region 2 (central) from EEG channels C3, Cz, and C4 are shown. **D**) Mean relative powers of alpha frequency band activity calculated for region 3 (parietal) from EEG channels P3, Pz, and P4 are shown.

**Figure 30.**
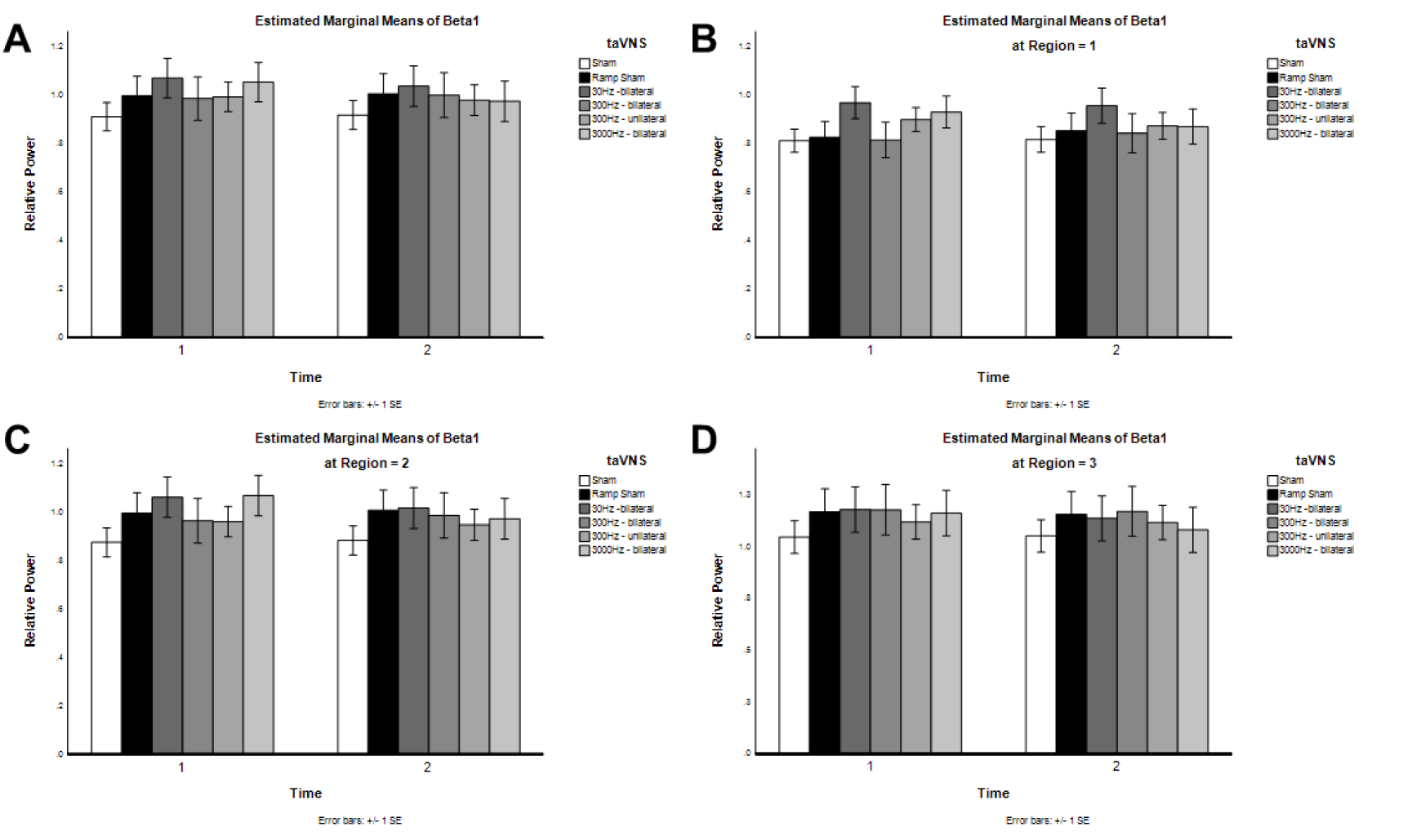
Influence of transdermal auricular vagal nerve stimulation on beta 1 EEG frequency band activity. **A**) The mean relative powers of beta 1 (12 – 15 Hz) frequency band activity calculated from EEG channels F3, Fz, F4, C3, Cz, C4, P3, Pz, and P4 are shown as calculated for inactive sham (sham), ramp sham, 30 Hz bilateral taVNS, 300 Hz bilateral taVNS, 300 Hz unilateral (left) taVNS), and 3000 Hz taVNS treatment conditions in responses to frequent tones before (Time 1) and after taVNS (Time 2). **B**) Mean relative powers of beta 1 frequency band activity calculated for region 1 (frontal) from EEG channels F3, Fz, and F4 are shown. **C**) Mean relative powers of beta 1 frequency band activity calculated for region 2 (central) from EEG channels C3, Cz, and C4 are shown. **D**) Mean relative powers of beta 1 frequency band activity calculated for region 3 (parietal) from EEG channels P3, Pz, and P4 are shown.

**Figure 31.**
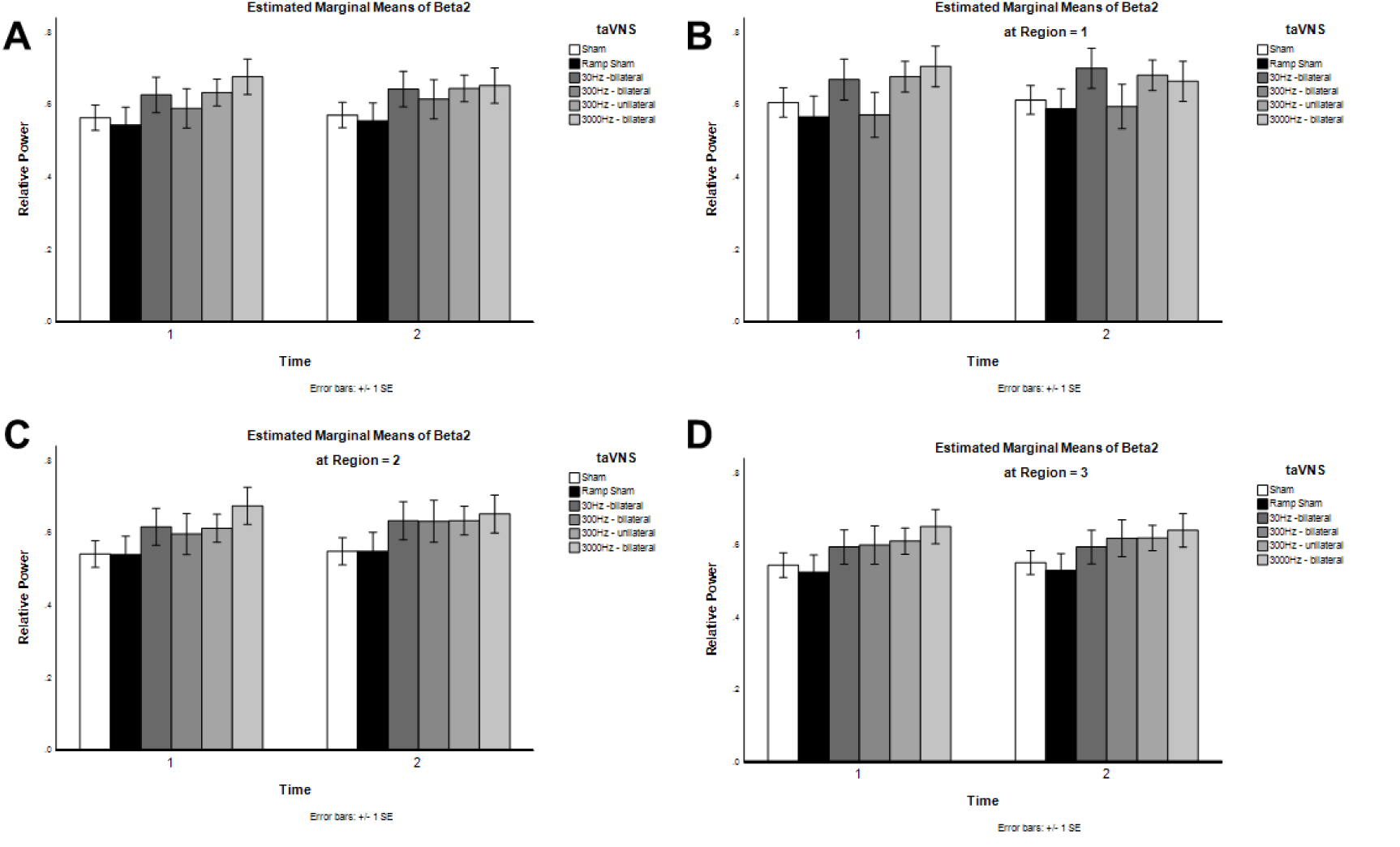
Influence of transdermal auricular vagal nerve stimulation on beta 2 EEG frequency band activity. **A**) The mean relative powers of beta 2 (15 – 20 Hz) frequency band activity calculated from EEG channels F3, Fz, F4, C3, Cz, C4, P3, Pz, and P4 are shown as calculated for inactive sham (sham), ramp sham, 30 Hz bilateral taVNS, 300 Hz bilateral taVNS, 300 Hz unilateral (left) taVNS), and 3000 Hz taVNS treatment conditions in responses to frequent tones before (Time 1) and after taVNS (Time 2). **B**) Mean relative powers of beta 2 frequency band activity calculated for region 1 (frontal) from EEG channels F3, Fz, and F4 are shown. **C**) Mean relative powers of beta 2 frequency band activity calculated for region 2 (central) from EEG channels C3, Cz, and C4 are shown. **D**) Mean relative powers of beta 2 frequency band activity calculated for region 3 (parietal) from EEG channels P3, Pz, and P4 are shown.

**Figure 32.**
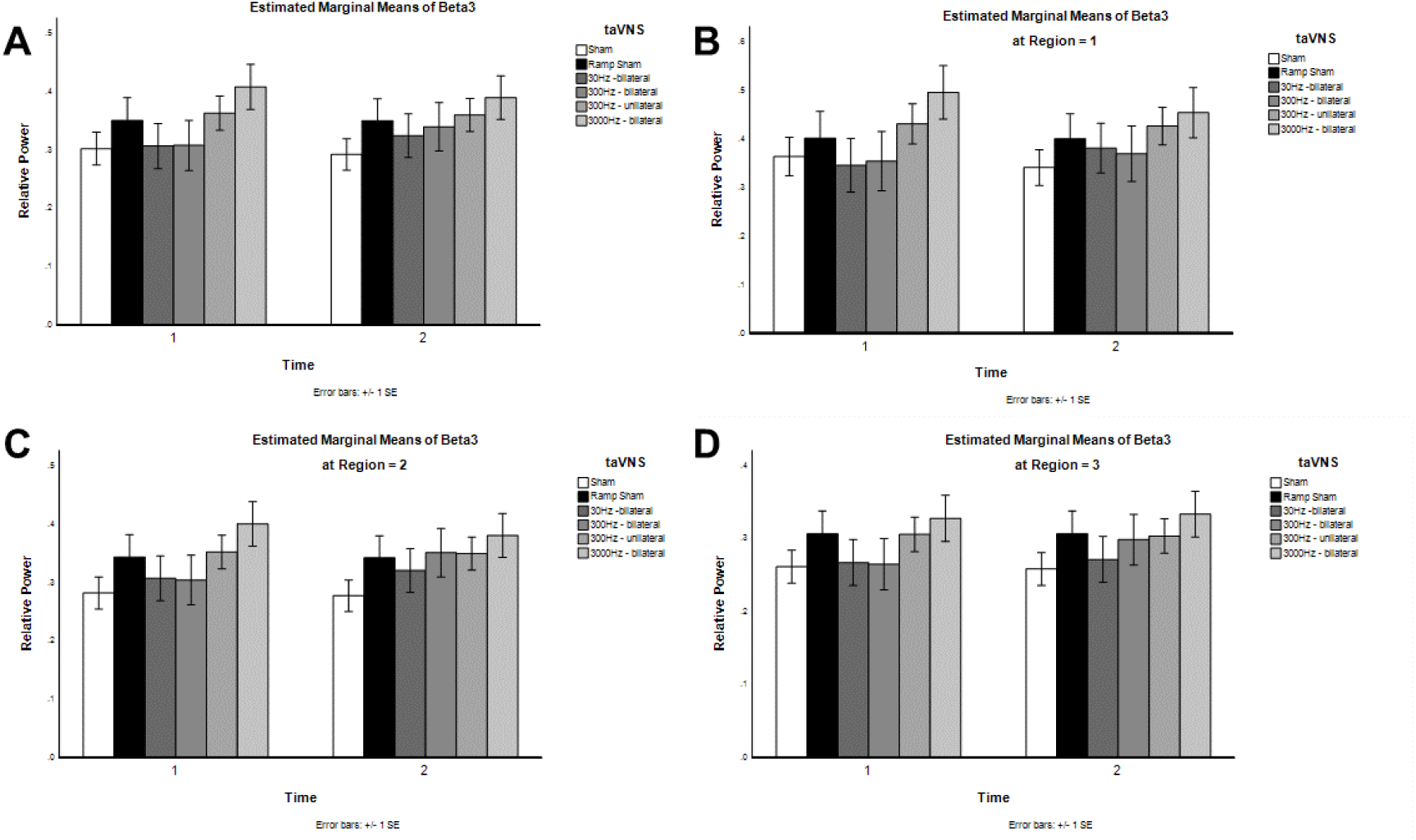
Influence of transdermal auricular vagal nerve stimulation on beta 3 EEG frequency band activity. **A**) The mean relative powers of beta 3 (20 – 30 Hz) frequency band activity calculated from EEG channels F3, Fz, F4, C3, Cz, C4, P3, Pz, and P4 are shown as calculated for inactive sham (sham), ramp sham, 30 Hz bilateral taVNS, 300 Hz bilateral taVNS, 300 Hz unilateral (left) taVNS), and 3000 Hz taVNS treatment conditions in responses to frequent tones before (Time 1) and after taVNS (Time 2). **B**) Mean relative powers of beta 3 frequency band activity calculated for region 1 (frontal) from EEG channels F3, Fz, and F4 are shown. **C**) Mean relative powers of beta 3 frequency band activity calculated for region 2 (central) from EEG channels C3, Cz, and C4 are shown. **D**) Mean relative powers of beta 3 frequency band activity calculated for region 3 (parietal) from EEG channels P3, Pz, and P4 are shown.

**Figure 33.**
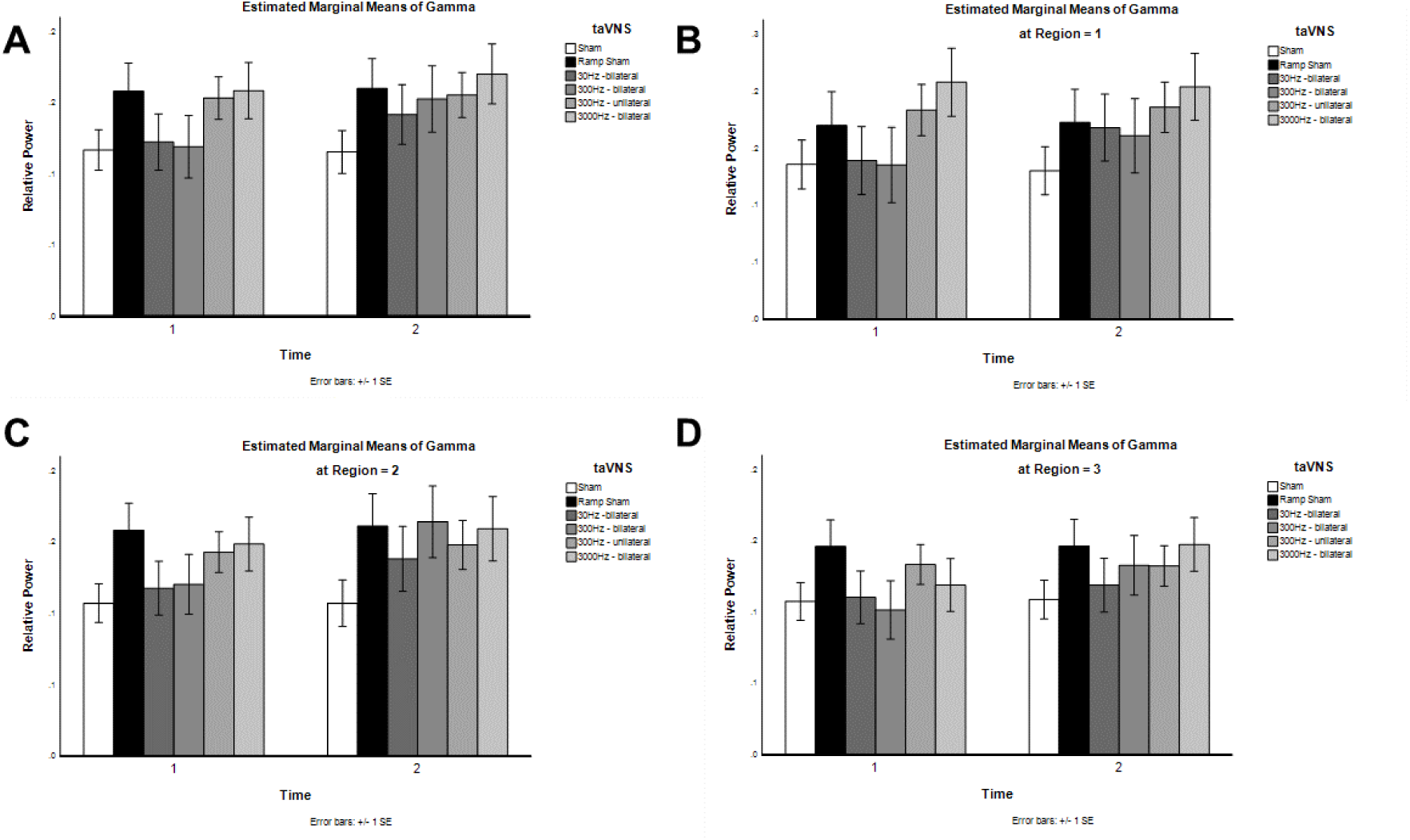
Influence of transdermal auricular vagal nerve stimulation on gamma EEG frequency band activity. **A**) The mean relative powers of gamma (30 – 40 Hz) frequency band activity calculated from EEG channels F3, Fz, F4, C3, Cz, C4, P3, Pz, and P4 are shown as calculated for inactive sham (sham), ramp sham, 30 Hz bilateral taVNS, 300 Hz bilateral taVNS, 300 Hz unilateral (left) taVNS), and 3000 Hz taVNS treatment conditions in responses to frequent tones before (Time 1) and after taVNS (Time 2). **B**) Mean relative powers of gamma frequency band activity calculated for region 1 (frontal) from EEG channels F3, Fz, and F4 are shown. **C**) Mean relative powers of gamma frequency band activity calculated for region 2 (central) from EEG channels C3, Cz, and C4 are shown. **D**) Mean relative powers of gamma frequency band activity calculated for region 3 (parietal) from EEG channels P3, Pz, and P4 are shown.

Statistical analyses conducted using ANOVAs revealed significant (p < 0.05) main effects of Time on the absolute power of the delta EEG band and significant main Time effects on the relative powers of delta, theta, and alpha EEG bands (Table 7). There were significant Time*taVNS Frequency interactions on the absolute powers of delta, theta, and beta 1 EEG bands, as well as on the relative power of the delta EEG band. Significant simple effects of Time were found for the 300 Hz unilateral condition on absolute and relative powers of the delta band and absolute power of the beta 1 EEG frequency band. Significant simple effects of Time were found for the 3000 Hz bilateral condition on the absolute powers of the alpha and beta 2 frequency bands, as well as the relative power of the beta 1 band. The 30 Hz condition also yielded a significant simple Time effect on the delta frequency band while the ramp sham condition also had significant simple Time effects on the relative powers of the delta and theta frequency bands. There were no simple significant Time effects for the inactive sham or 300 Hz bilateral conditions.

### Influence of taVNS on EEG Auditory Mismatch Negativity

One of our primary aims has been to determine if taVNS can produce acute changes in auditory plasticity by modulating auditory responses to “oddball” stimuli. To this end, we implemented a standard auditory mismatch negativity task in which volunteer human research subjects passively listened to *frequent* and *infrequent* pure tones. We recorded the EEG responses to the frequent and infrequent tones before and after taVNS treatment. From the auditory evoked potentials obtained in response to the tones we calculated the mismatch negativity (MMN) potential or EEG auditory oddball response. Data from these investigations are illustrated in Figures 34-39 below.

**Figure 34.**
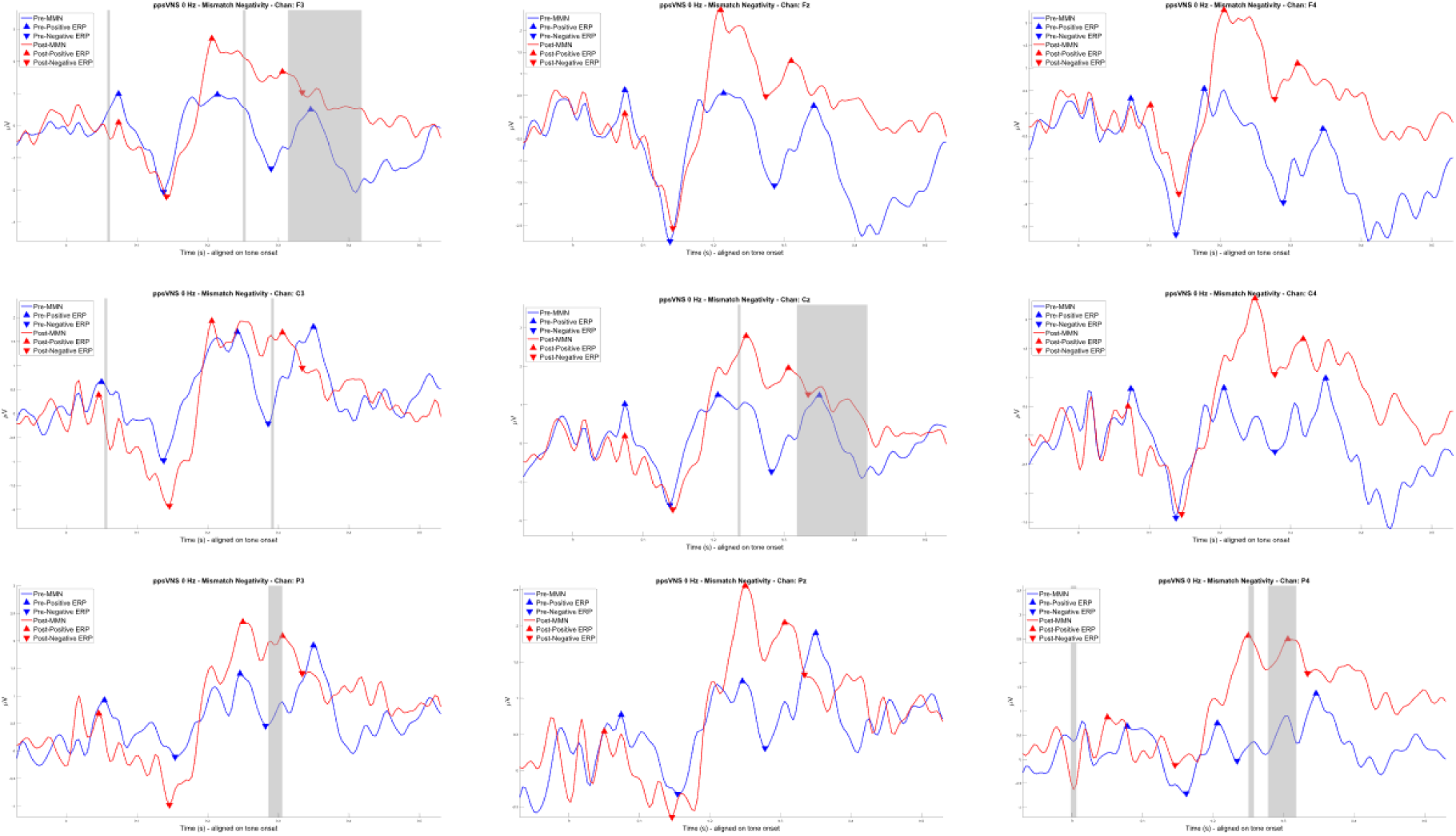
Influence of inactive sham taVNS on auditory MMN potentials. Starting from the upper left and moving across to the right are shown MMN potentials before (blue) and after (red) sham treatment for EEG channels F3, Fz, and F4. Similarly, taVNS inactive sham effects produced on MMN potentials for EEG sites C3, Cz, and C4 are shown in the middle row from left to right respectively. MMN potentials are shown for EEG sites P3, Pz, and P4 on the bottom row. Time regions highlighted by grey windows indicate regions of significant (p < 0.05) difference between the pre- and post-treatment MMN potentials.

**Figure 35.**
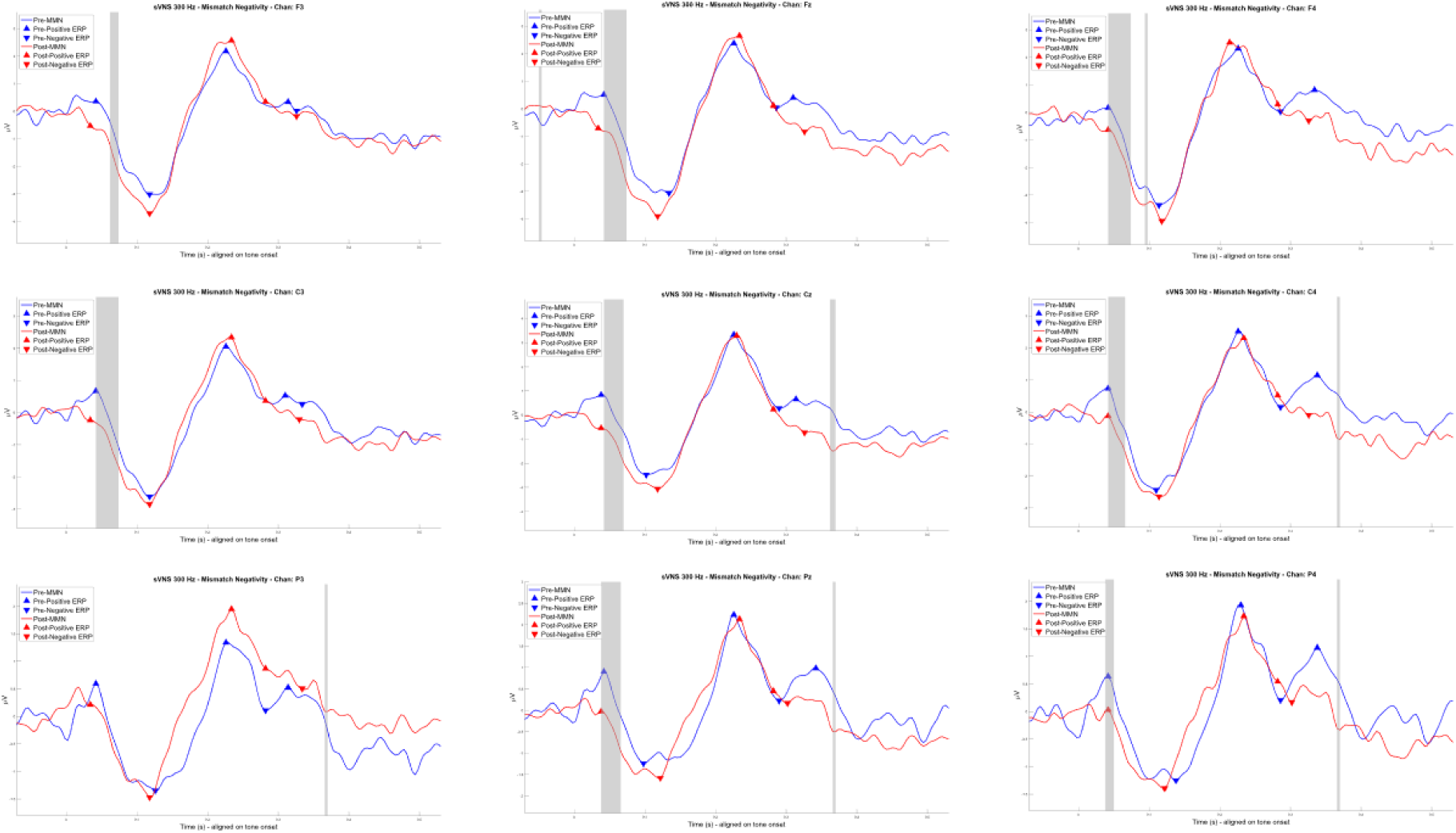
Influence of ramping sham taVNS treatment on auditory MMN potentials. Starting from the upper left and moving across to the right are shown MMN potentials before (blue) and after (red) ramping sham treatment for EEG channels F3, Fz, and F4. Similarly, taVNS ramped sham effects produced on MMN potentials for EEG sites C3, Cz, and C4 are shown in the middle row from left to right respectively. MMN potentials are shown for EEG sites P3, Pz, and P4 on the bottom row. Time regions highlighted by grey windows indicate regions of significant (p < 0.05) difference between the pre- and post-treatment MMN potentials.

**Figure 36.**
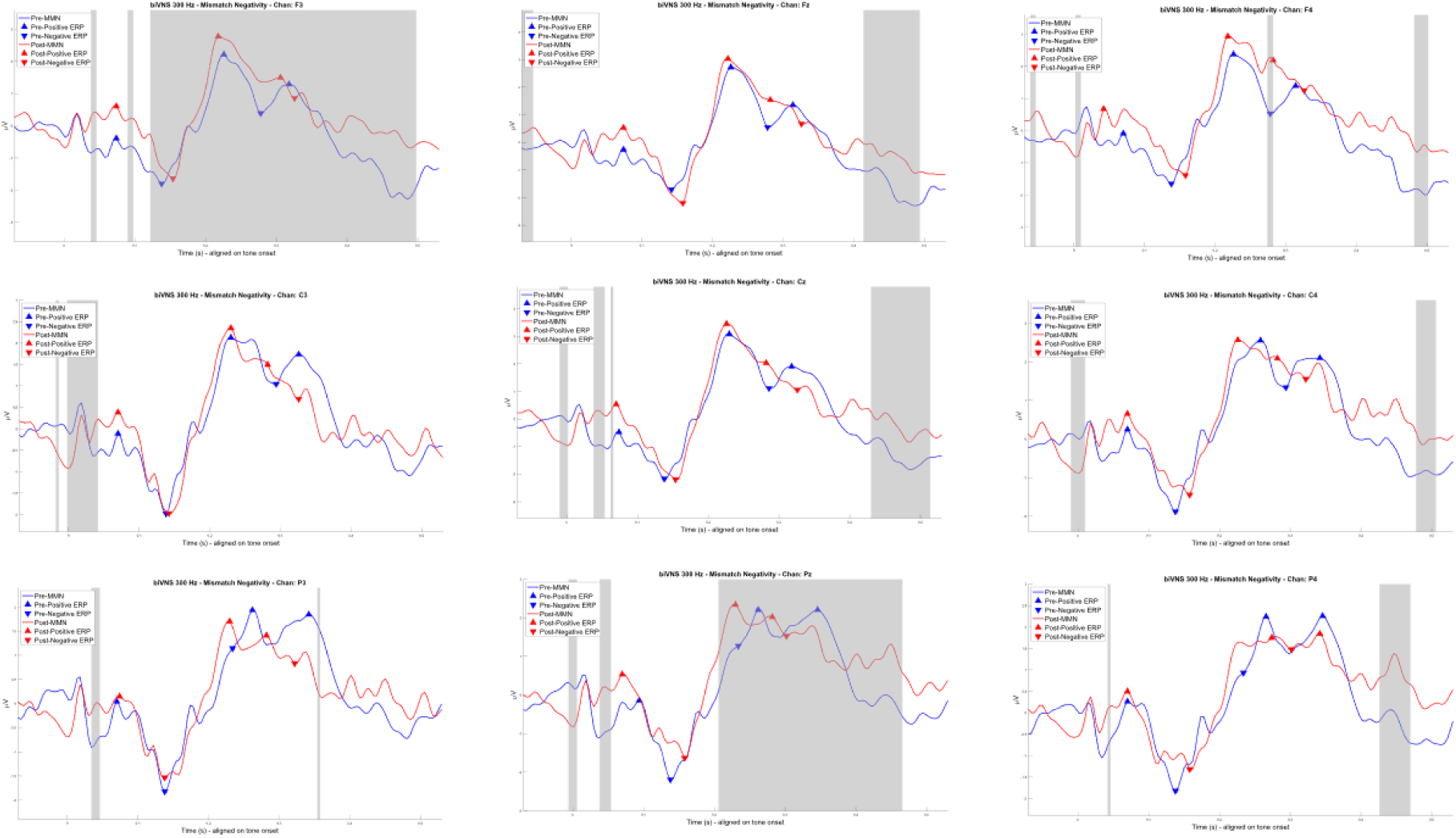
Influence of 300 Hz bilateral taVNS on auditory MMN potentials. Starting from the upper left and moving across to the right are shown MMN potentials before (blue) and after (red) 300 Hz bilateral taVNS for EEG channels F3, Fz, and F4. Similarly, 300 Hz bilateral taVNS effects produced on MMN potentials for EEG sites C3, Cz, and C4 are shown in the middle row from left to right respectively. MMN potentials are shown for EEG sites P3, Pz, and P4 on the bottom row. Time regions highlighted by grey windows indicate regions of significant (p < 0.05) difference between the pre- and post-treatment MMN potentials.

**Figure 37.**
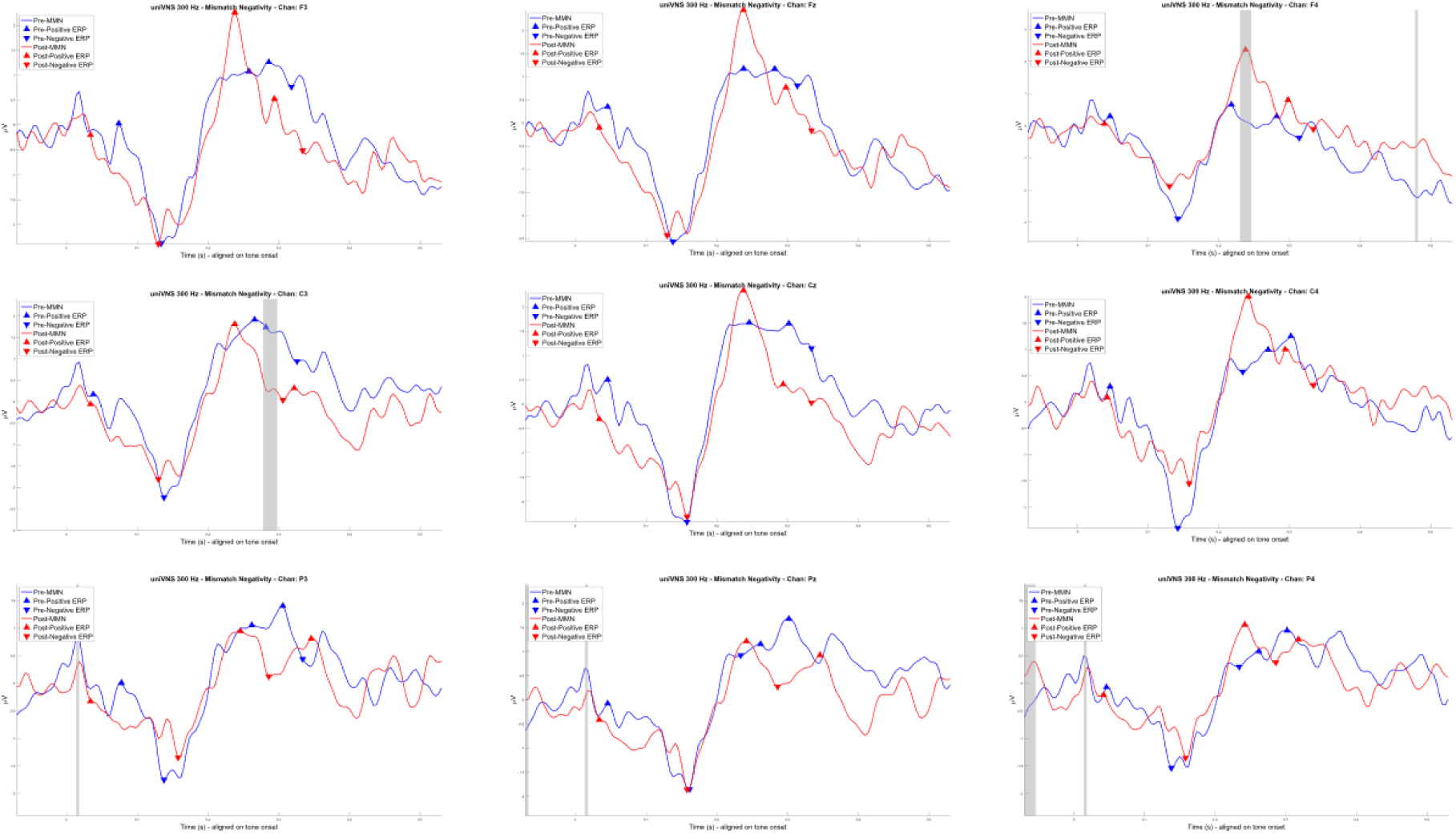
Influence of 300 Hz left unilateral taVNS on auditory MMN potentials. Starting from the upper left and moving across to the right are shown MMN potentials before (blue) and after (red) 300 Hz left unilateral taVNS for EEG channels F3, Fz, and F4. Similarly, 300 Hz left unilateral taVNS effects produced on MMN potentials for EEG sites C3, Cz, and C4 are shown in the middle row from left to right respectively. MMN potentials are shown for EEG sites P3, Pz, and P4 on the bottom row. Time regions highlighted by grey windows indicate regions of significant (p < 0.05) difference between the pre- and post-treatment MMN potentials.

**Figure 38.**
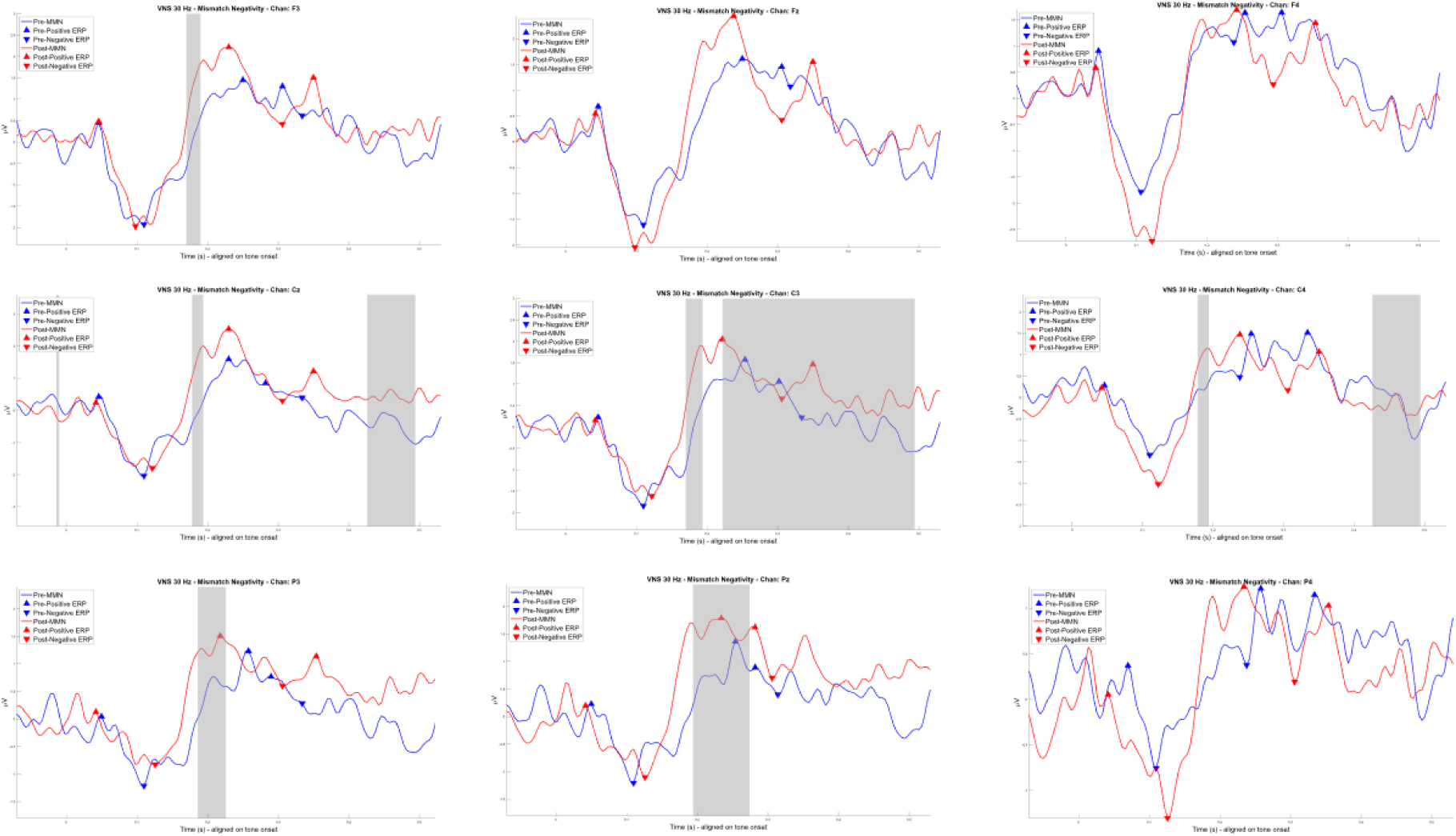
Influence of 30 Hz bilateral taVNS on auditory MMN potentials. Starting from the upper left and moving across to the right are shown MMN potentials before (blue) and after (red) 30 Hz bilateral taVNS for EEG channels F3, Fz, and F4. Similarly, 30 Hz bilateral taVNS effects produced on MMN potentials for EEG sites C3, Cz, and C4 are shown in the middle row from left to right respectively. MMN potentials are shown for EEG sites P3, Pz, and P4 on the bottom row. Time regions highlighted by grey windows indicate regions of significant (p < 0.05) difference between the pre- and post-treatment MMN potentials.

**Figure 39.**
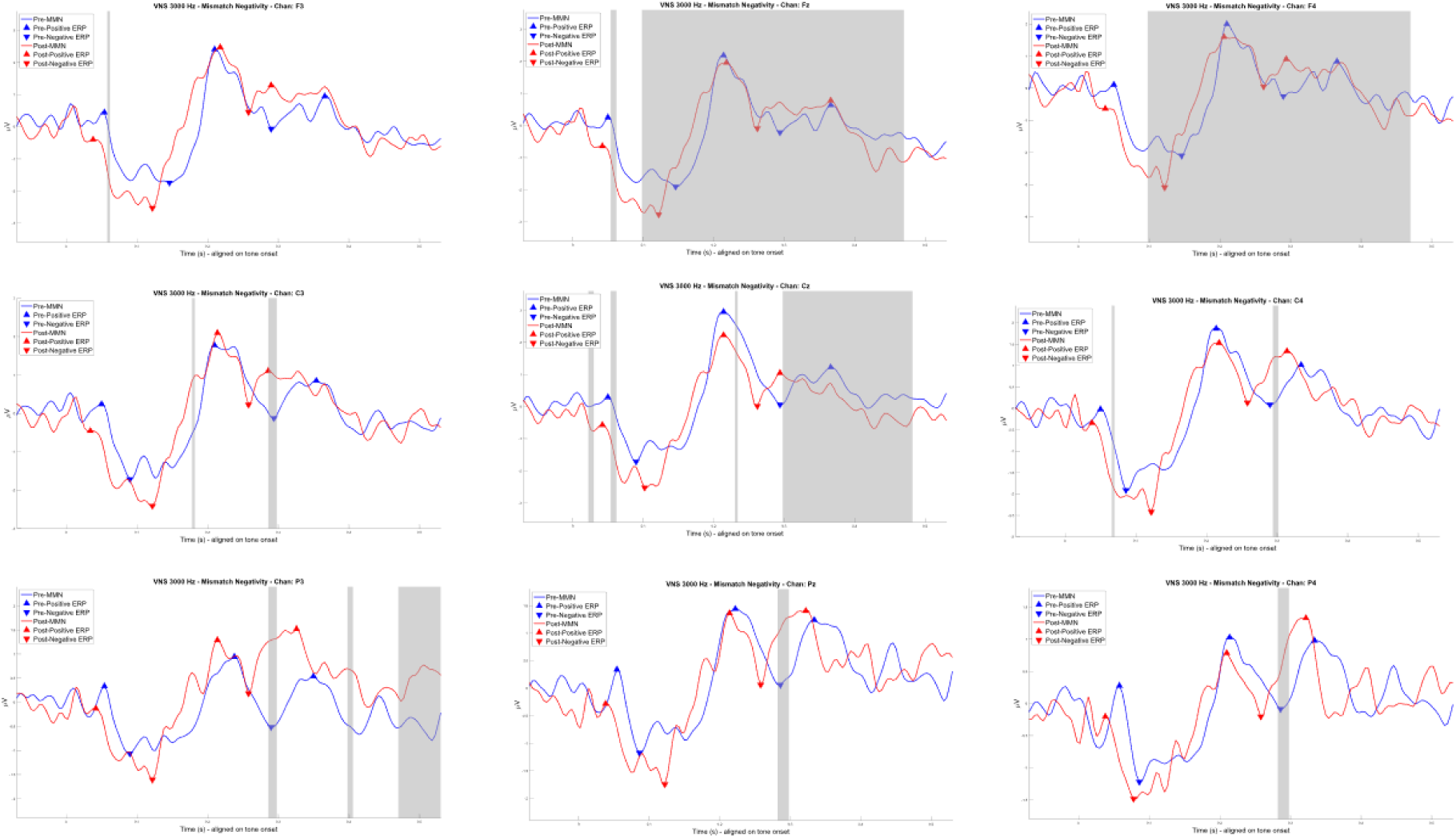
Influence of 3000 Hz bilateral taVNS on auditory MMN potentials. Starting from the upper left and moving across to the right are shown MMN potentials before (blue) and after (red) 3000 Hz bilateral taVNS for EEG channels F3, Fz, and F4. Similarly, 3000 Hz bilateral taVNS effects produced on MMN potentials for EEG sites C3, Cz, and C4 are shown in the middle row from left to right respectively. MMN potentials are shown for EEG sites P3, Pz, and P4 on the bottom row. Time regions highlighted by grey windows indicate regions of significant (p < 0.05) difference between the pre- and post-treatment MMN potentials.

### Influence of taVNS on Pupil Dynamics

All prior data shown have been collected using protocols described in the methods for Experiment 1. In order to investigate the influence of taVNS on pupil dynamics we changed the timing of auditory stimulus presentations to accommodate for the slower, time-lagging pupil responses compared to EEG responses. Thus in Experiment 2, we used a 23 min passive auditory oddball task designed to elicit P1, N1, P2, N2, P3, and mismatch negativity (MMN) ERP components in an eyes-open condition. The task consisted of a series of randomized standard/frequent (N = 450, 750 Hz) and deviant/infrequent (N = 100, 1500 Hz) tones with an inter-stimulus interval of 2500 ms to allow for acquisition of pupillometry data. Stimuli were presented over headphones and participants were instructed to keep their eyes focused on a fixation cross while passively attending to the stimuli. This task was repeated after the stimulation task.

Pupil data was preprocessed using methods set forth by Geller and colleagues [42]. First, samples with unrealistic pupil diameters (< 2 mm or > 8 mm) pupil diameters greater than or less than three standard deviations from the mean were removed. Samples occurring between 100 ms before and 100 ms after blinks were also marked as invalid. Samples were then linearly interpolated. Any artifacts due to rapid changes in pupil diameter, as determined by median absolute deviation, were then removed [43]. A 5-point moving average was applied to smooth the samples. The continuous data was then aligned on tone-onset, converted to z-scores, and baseline corrected by subtracting the median activity 0.5 s before tone-onset from each sample. Data from these investigations are illustrated in Figures 40-42.

**Figure 40.**
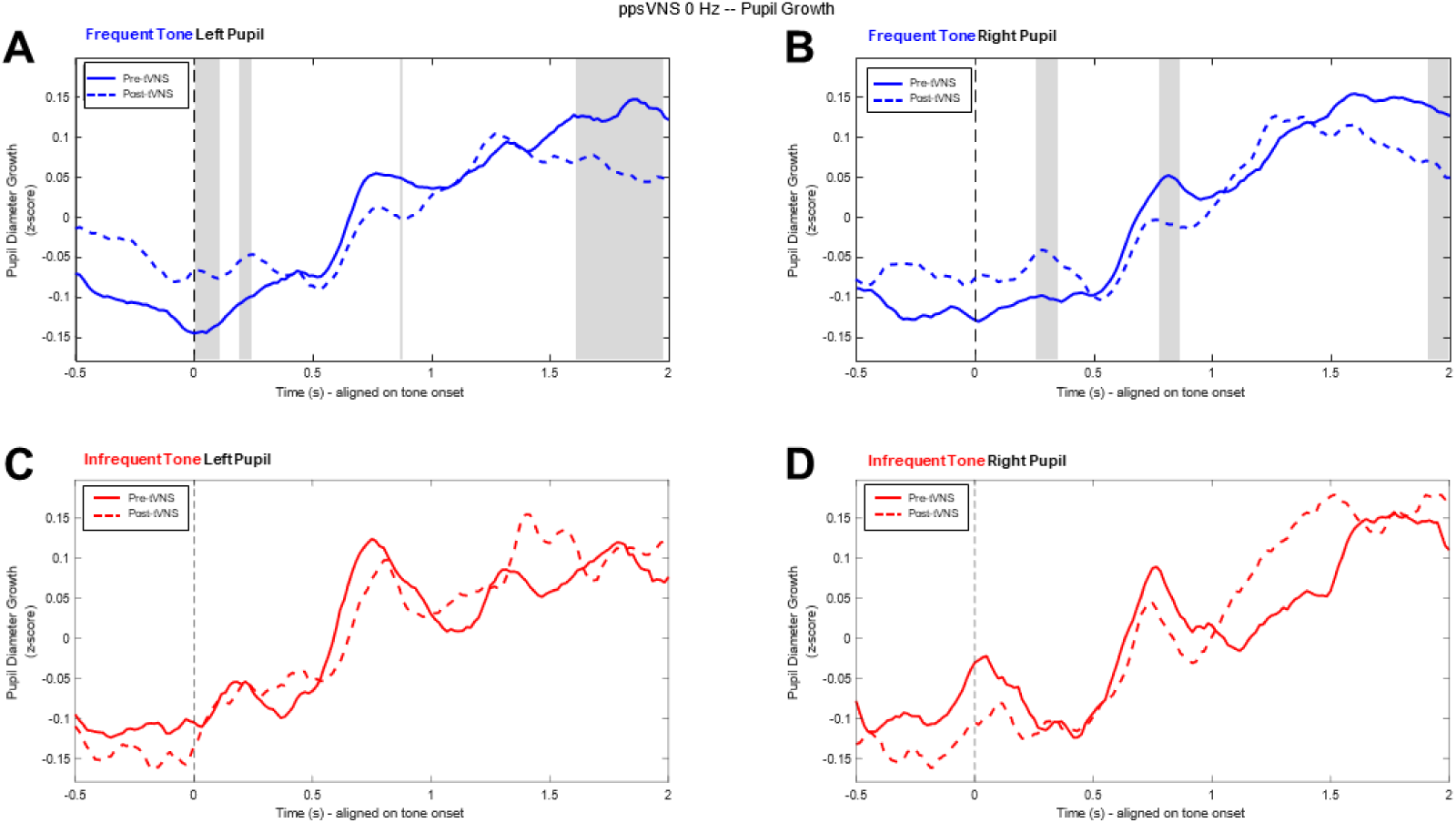
Influence of inactive sham taVNS on pupil growth curves obtained during passive auditory oddball tasks. A) Left pupil growth curves shown as Z-scores from data obtained during passive auditory oddball listening tasks in response to frequent tones before (solid line) and after (dashed line) inactive sham taVNS treatment. B) Right pupil growth curves are similarly shown in response to frequent tones. C) Left and D) right pupil growth curves in response to infrequent tones are shown for data obtained before (solid line) and after (dashed line) sham taVNS treatment. Time regions highlighted by grey windows indicate regions of significant (p < 0.05) difference between the pre- and post-treatment conditions.

**Figure 41.**
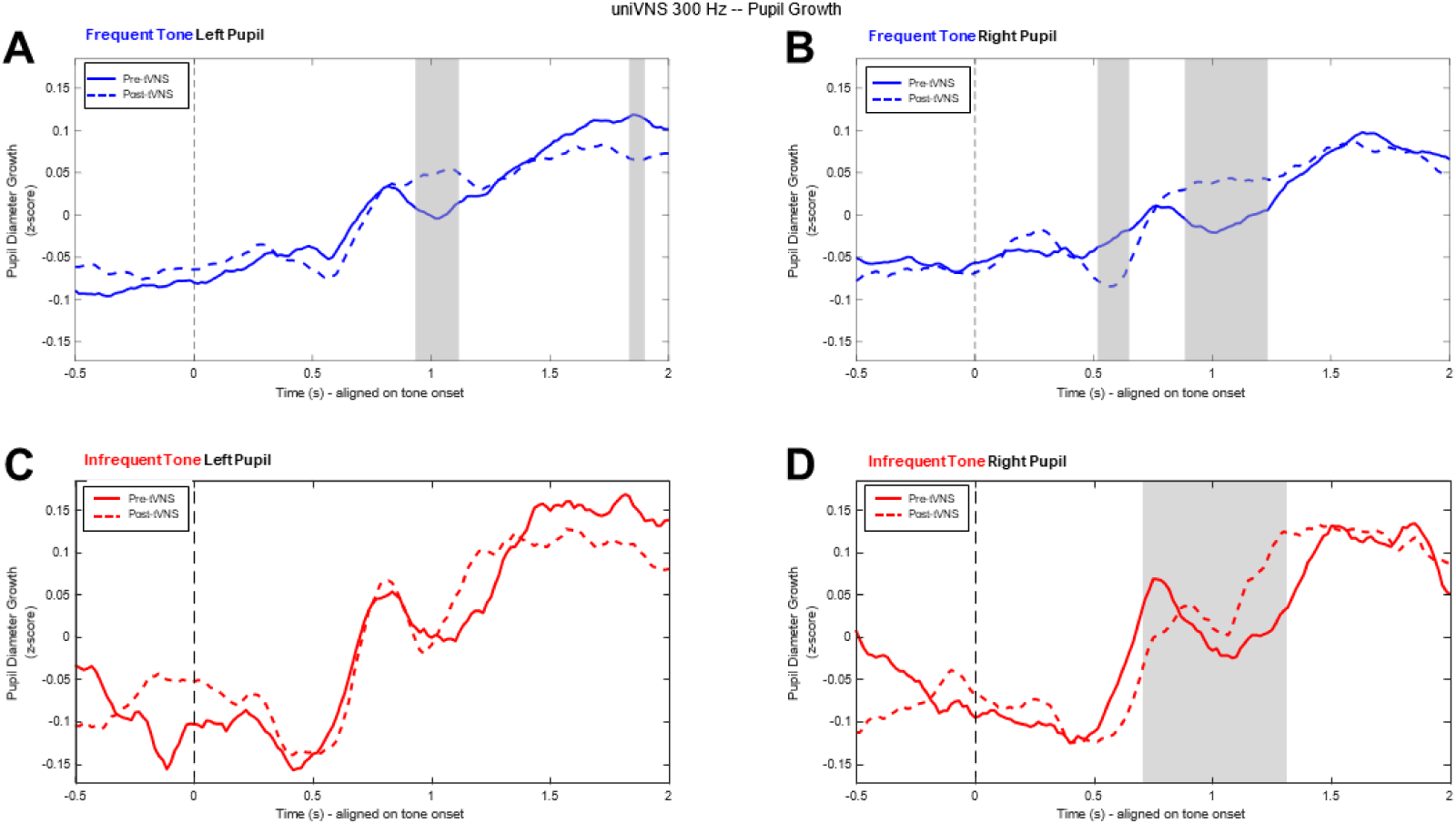
Influence of left unilateral 300 Hz taVNS on pupil growth curves obtained during passive auditory oddball tasks. A) Left pupil growth curves shown as Z-scores from data obtained during passive auditory oddball listening tasks in response to frequent tones before (solid line) and after (dashed line) left unilateral 300 Hz taVNS. B) Right pupil growth curves are similarly shown in response to frequent tones. C) Left and D) right pupil growth curves in response to infrequent tones are shown for data obtained before (solid line) and after (dashed line) left unilateral 300 Hz taVNS treatment. Time regions highlighted by grey windows indicate regions of significant (p < 0.05) difference between the pre- and post-treatment conditions.

**Figure 42.**
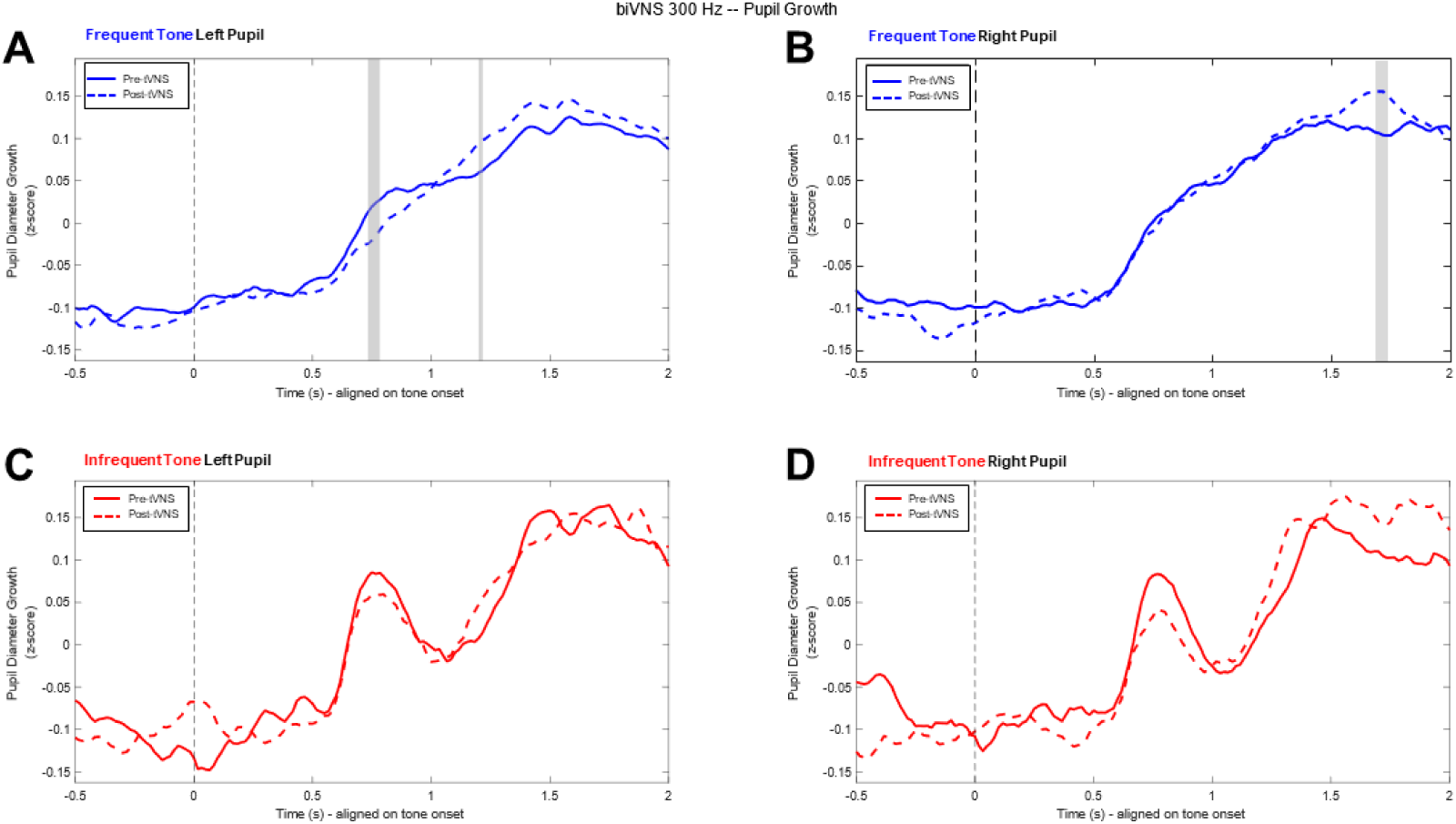
Influence of bilateral 300 Hz taVNS on pupil growth curves obtained during passive auditory oddball tasks. A) Left pupil growth curves shown as Z-scores from data obtained during passive auditory oddball listening tasks in response to frequent tones before (solid line) and after (dashed line) bilateral 300 Hz taVNS. B) Right pupil growth curves are similarly shown in response to frequent tones. C) Left and D) right pupil growth curves in response to infrequent tones are shown for data obtained before (solid line) and after (dashed line) bilateral 300 Hz taVNS treatment. Time regions highlighted by grey windows indicate regions of significant (p < 0.05) difference between the pre- and post-treatment conditions.

### Safety and Tolerability of taVNS

Besides assessing basic neuro- and psychophysiological outcomes in healthy human research subjects as described above, we also evaluated the acute safety and tolerability of taVNS in our performance efforts. Immediately following taVNS treatment and tasks, subjects respond to a short questionnaire in which they are asked to subjectively describe their level of comfort and/or discomfort produced by the treatment. Again 24 hours following the study, participants were asked to respond to a few short follow-up questions related to potential adverse reactions, such as skin irritation or occurrence of headaches. Statistical results from these investigations are summarized in Table 8 and data are illustrated in Figures 43-54.

**Table 8.**
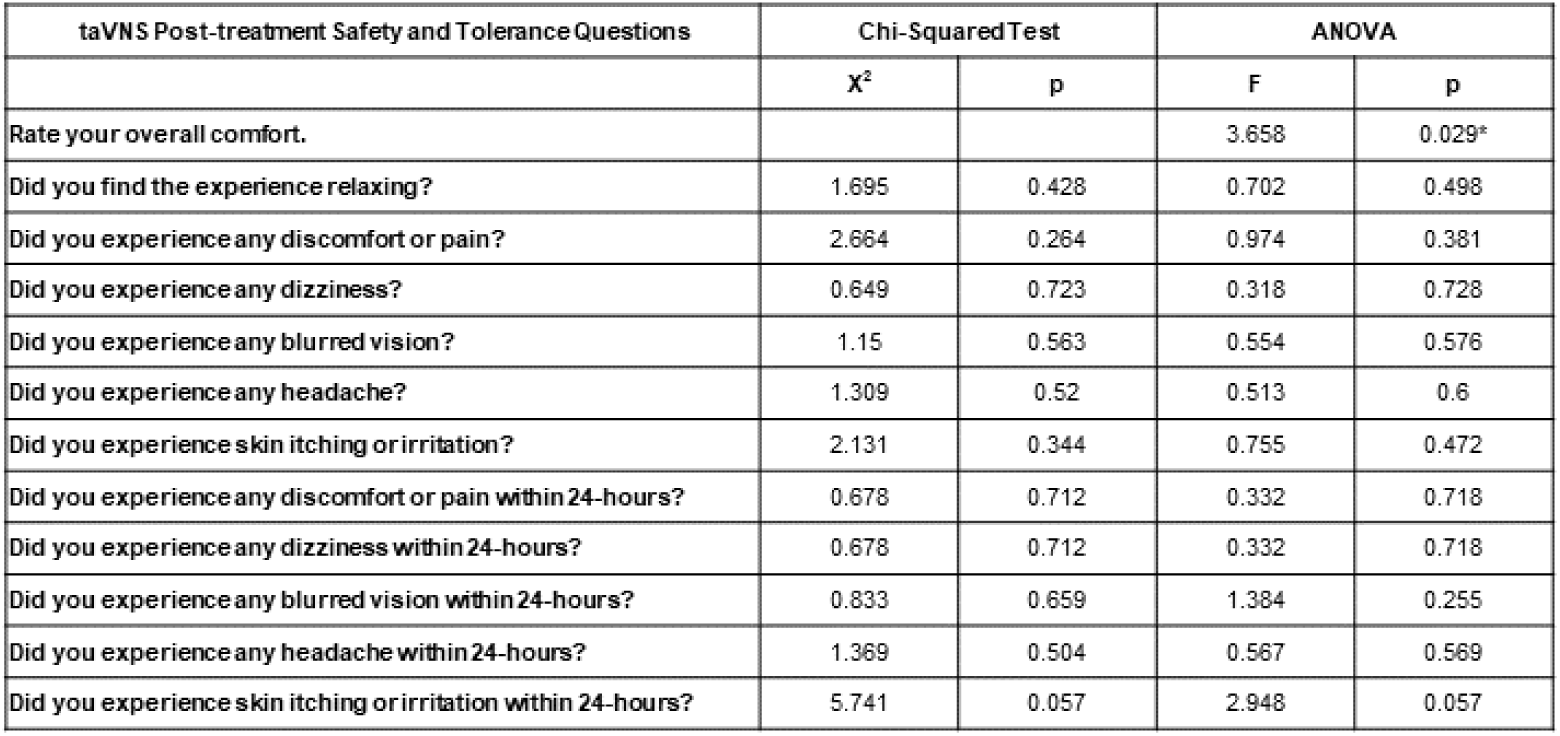
Statistical results table summarizing acute safety and tolerability outcomes comparing sham treatments to active taVNS.

**Figure 43.**
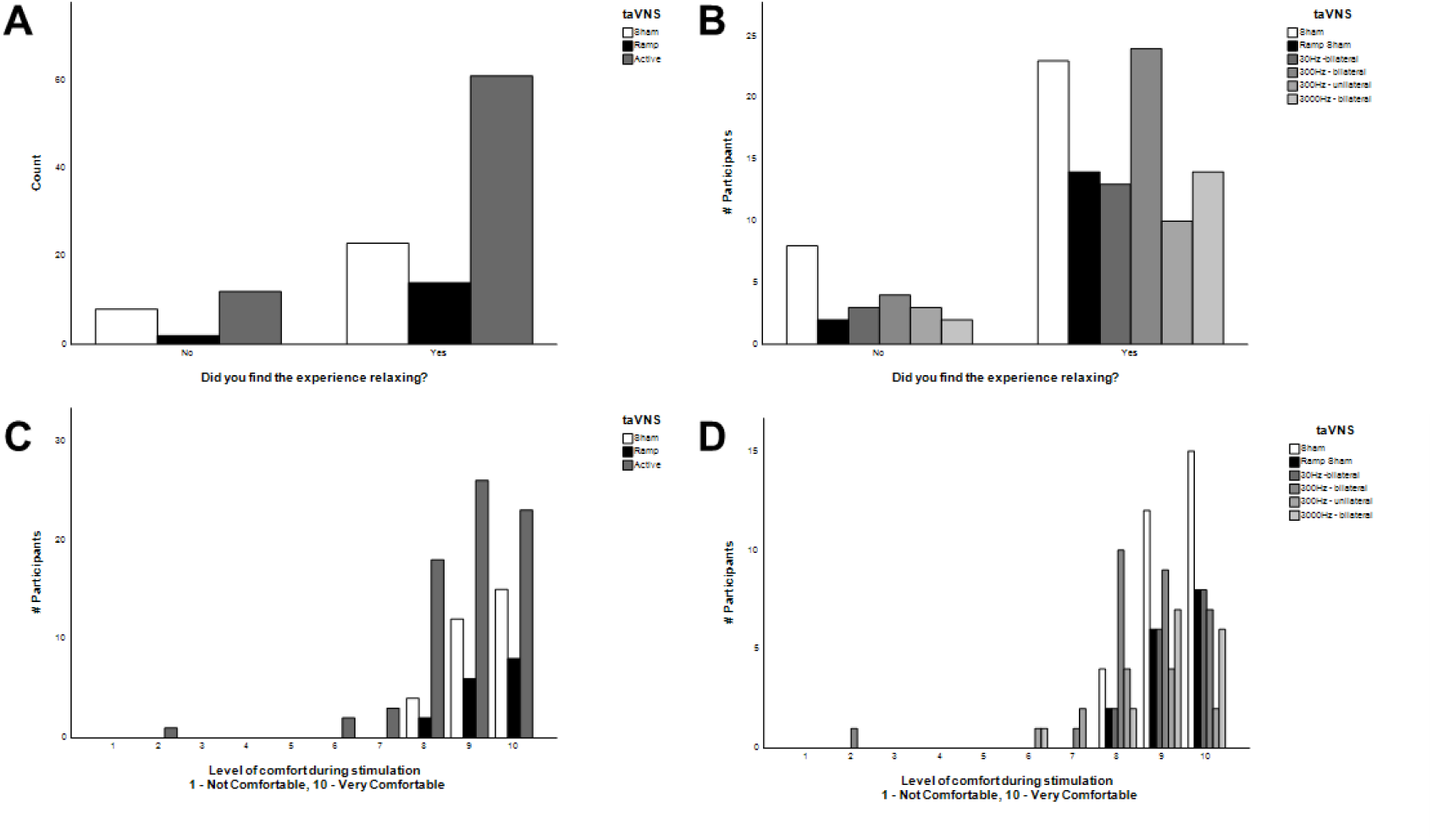
Influence of acute taVNS on relaxation and comfort. A) Subject responses are shown for question pertaining to relaxation for inactive sham, ramp sham, and all active taVNS groups collapsed. B) Data as shown in panel A but illustrated by individual taVNS treatment groups. C) Subject responses are shown for question pertaining to comfort for inactive sham, ramp sham, and all active taVNS groups collapsed. D) Data as shown in panel C but illustrated by individual taVNS treatment groups.

**Figure 44.**
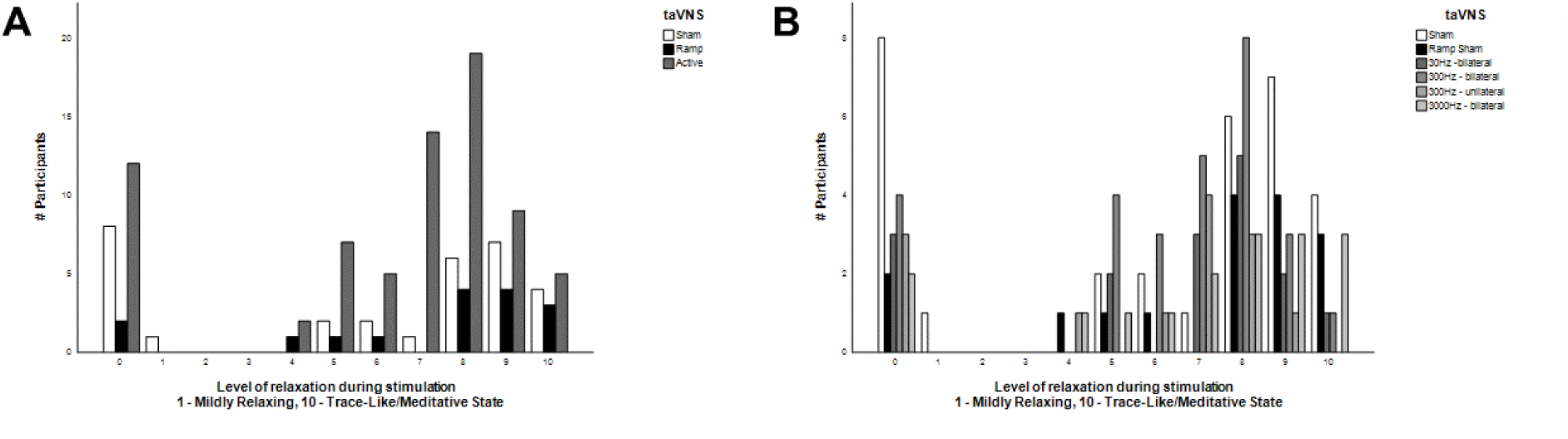
Influence of acute taVNS on level of relaxation. A) Subject responses are shown for question pertaining to level of relaxation for inactive sham, ramp sham, and all active taVNS groups collapsed. B) Data as shown in panel A but illustrated by individual taVNS treatment groups.

**Figure 45.**
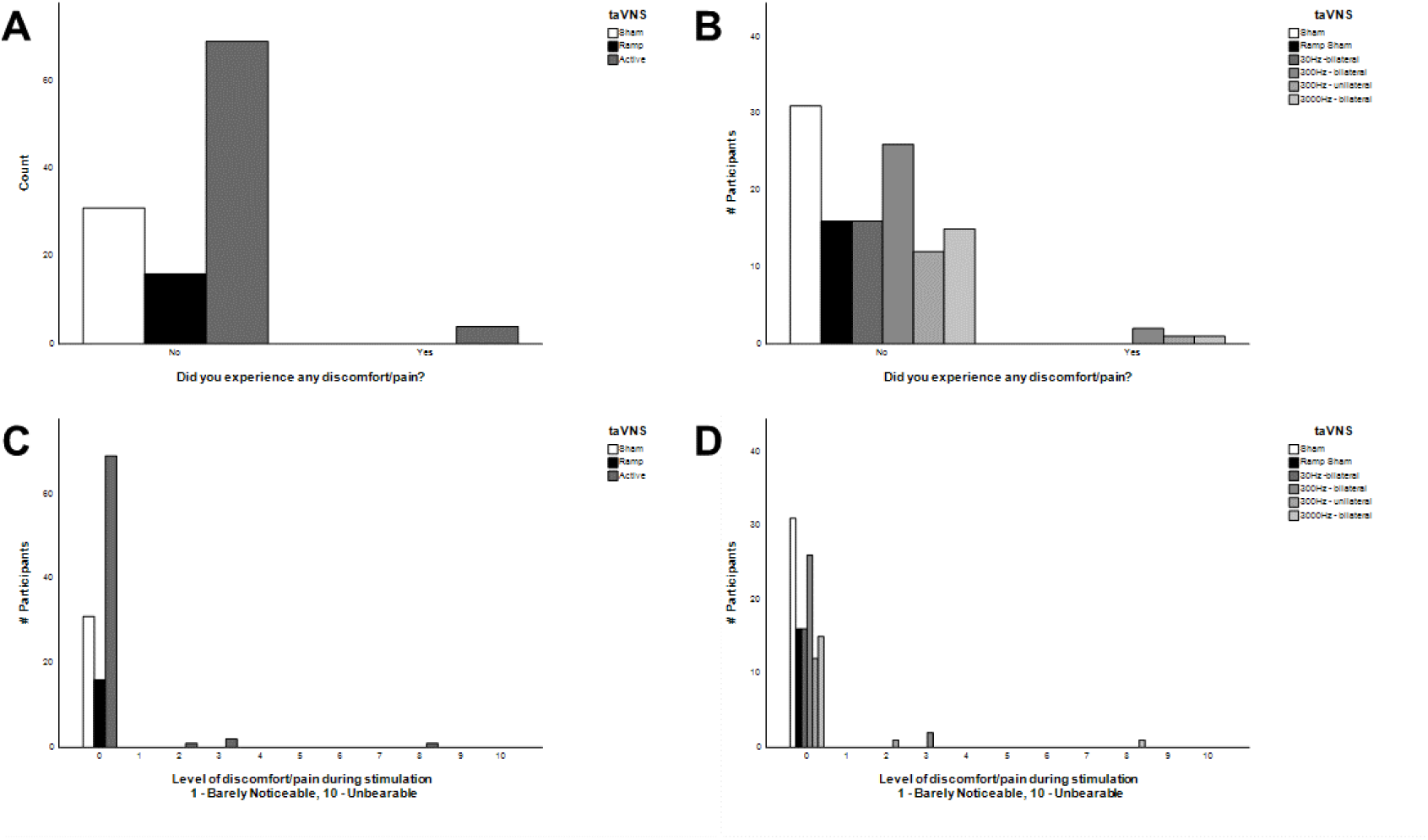
Influence of acute taVNS on discomfort. A) Subject responses are shown for question pertaining to discomfort for inactive sham, ramp sham, and all active taVNS groups collapsed. B) Data as shown in panel A but illustrated by individual taVNS treatment groups. C) Subject responses are shown for question pertaining to the level of discomfort experienced during stimulation for inactive sham, ramp sham, and all active taVNS groups collapsed. D) Data as shown in panel C but illustrated by individual taVNS treatment groups.

**Figure 46.**
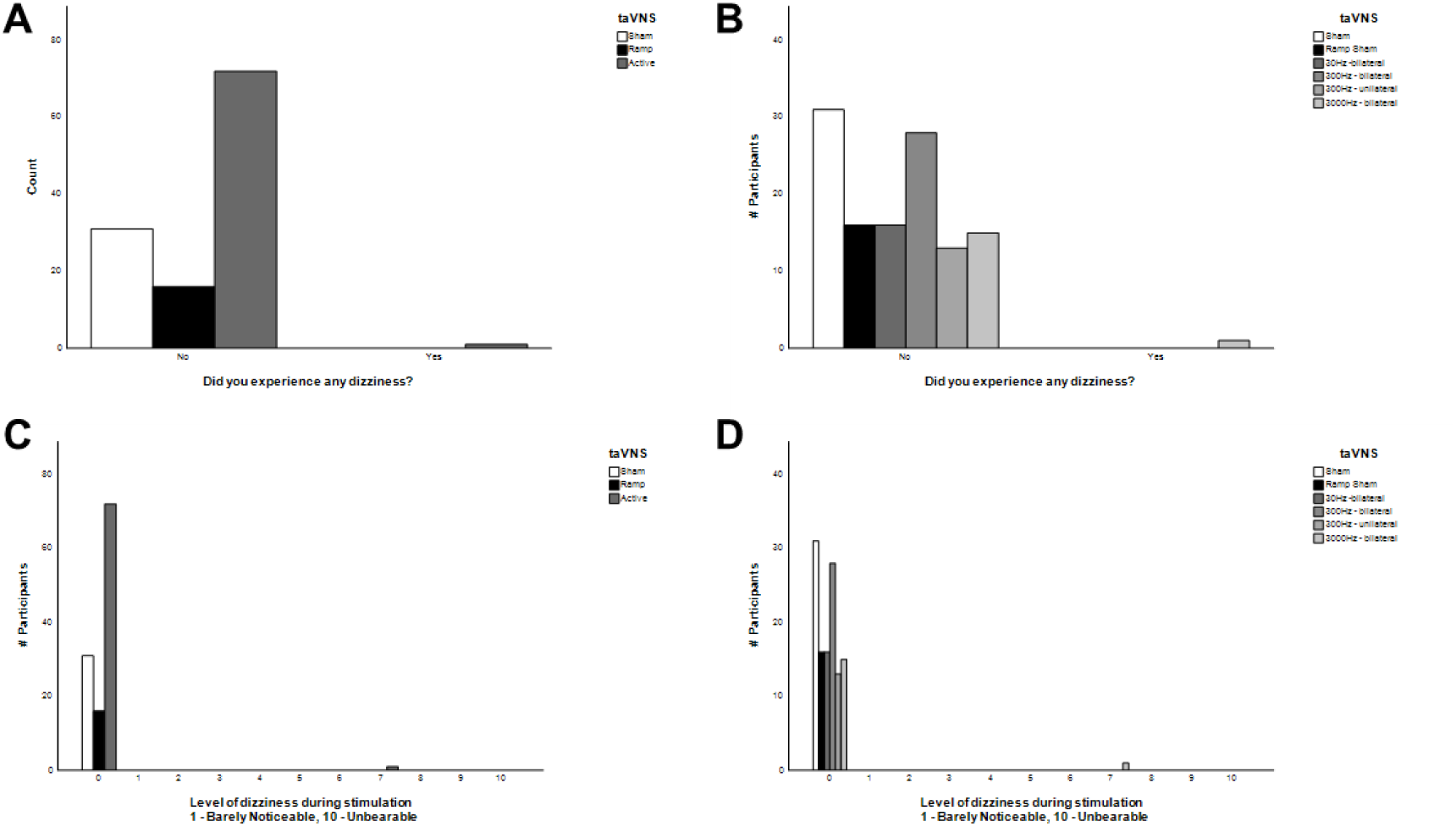
Influence of acute taVNS on dizziness. A) Subject responses are shown for question pertaining to dizziness for inactive sham, ramp sham, and all active taVNS groups collapsed. B) Data as shown in panel A but illustrated by individual taVNS treatment groups. C) Subject responses are shown for question pertaining to the level of dizziness experienced during stimulation for inactive sham, ramp sham, and all active taVNS groups collapsed. D) Data as shown in panel C but illustrated by individual taVNS treatment groups.

**Figure 47.**
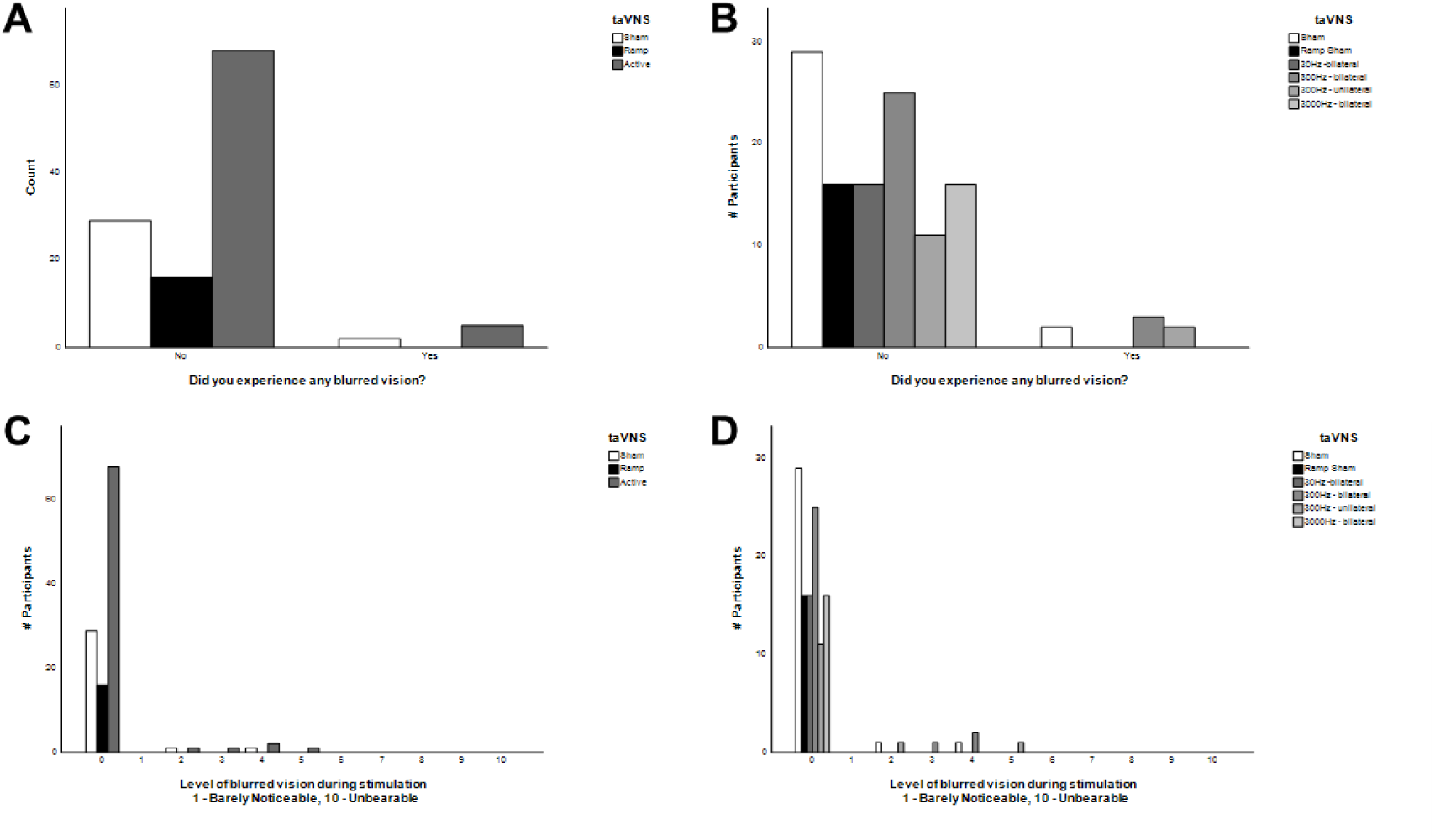
Influence of acute taVNS on blurred vision. A) Subject responses are shown for question pertaining to blurring of vision for inactive sham, ramp sham, and all active taVNS groups collapsed. B) Data as shown in panel A but illustrated by individual taVNS treatment groups. C) Subject responses are shown for question pertaining to the level of blurred vision experienced during stimulation for inactive sham, ramp sham, and all active taVNS groups collapsed. D) Data as shown in panel C but illustrated by individual taVNS treatment groups.

**Figure 48.**
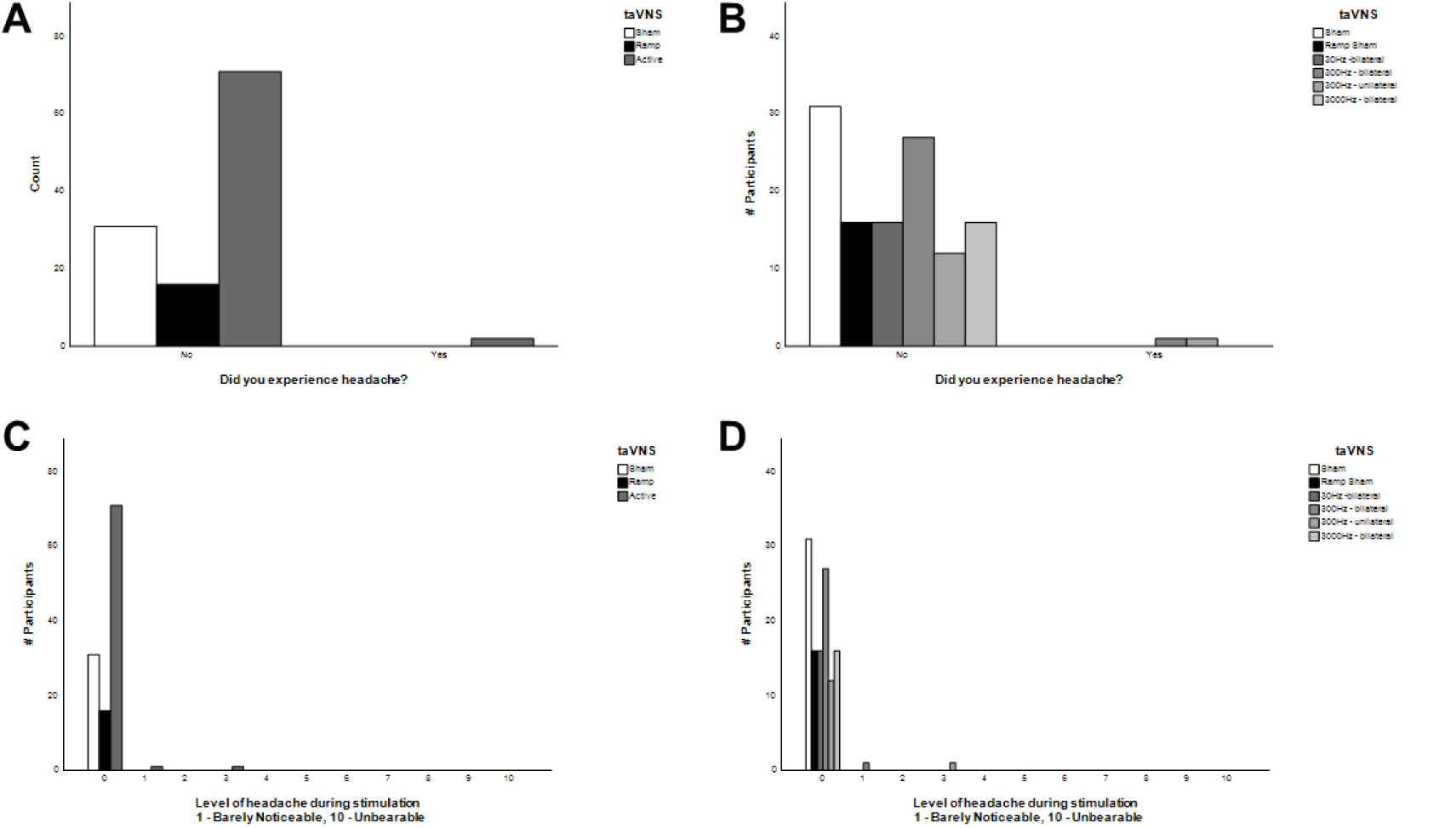
Influence of acute taVNS on headache. A) Subject responses are shown for question pertaining to the occurrence of a headache for inactive sham, ramp sham, and all active taVNS groups collapsed. B) Data as shown in panel A but illustrated by individual taVNS treatment groups. C) Subject responses are shown for question pertaining to the level of headache experienced during stimulation for inactive sham, ramp sham, and all active taVNS groups collapsed. D) Data as shown in panel C but illustrated by individual taVNS treatment groups.

**Figure 49.**
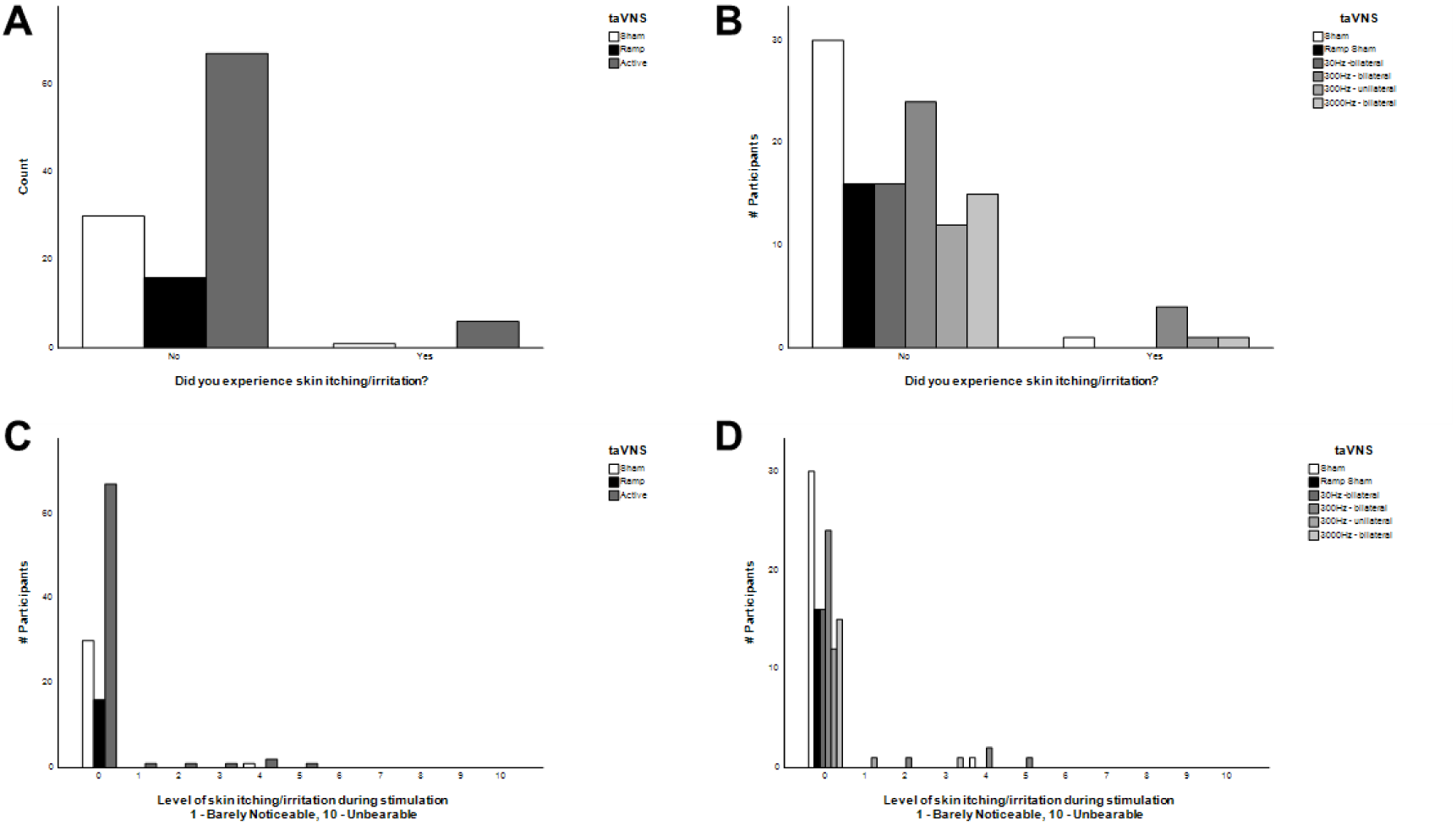
Influence of acute taVNS on skin itching and irritation. A) Subject responses are shown for question pertaining to the occurrence of skin itching or irritation for inactive sham, ramp sham, and all active taVNS groups collapsed. B) Data as shown in panel A but illustrated by individual taVNS treatment groups. C) Subject responses are shown for question pertaining to the level of skin itching or irritation experienced during stimulation for inactive sham, ramp sham, and all active taVNS groups collapsed. D) Data as shown in panel C but illustrated by individual taVNS treatment groups.

**Figure 50.**
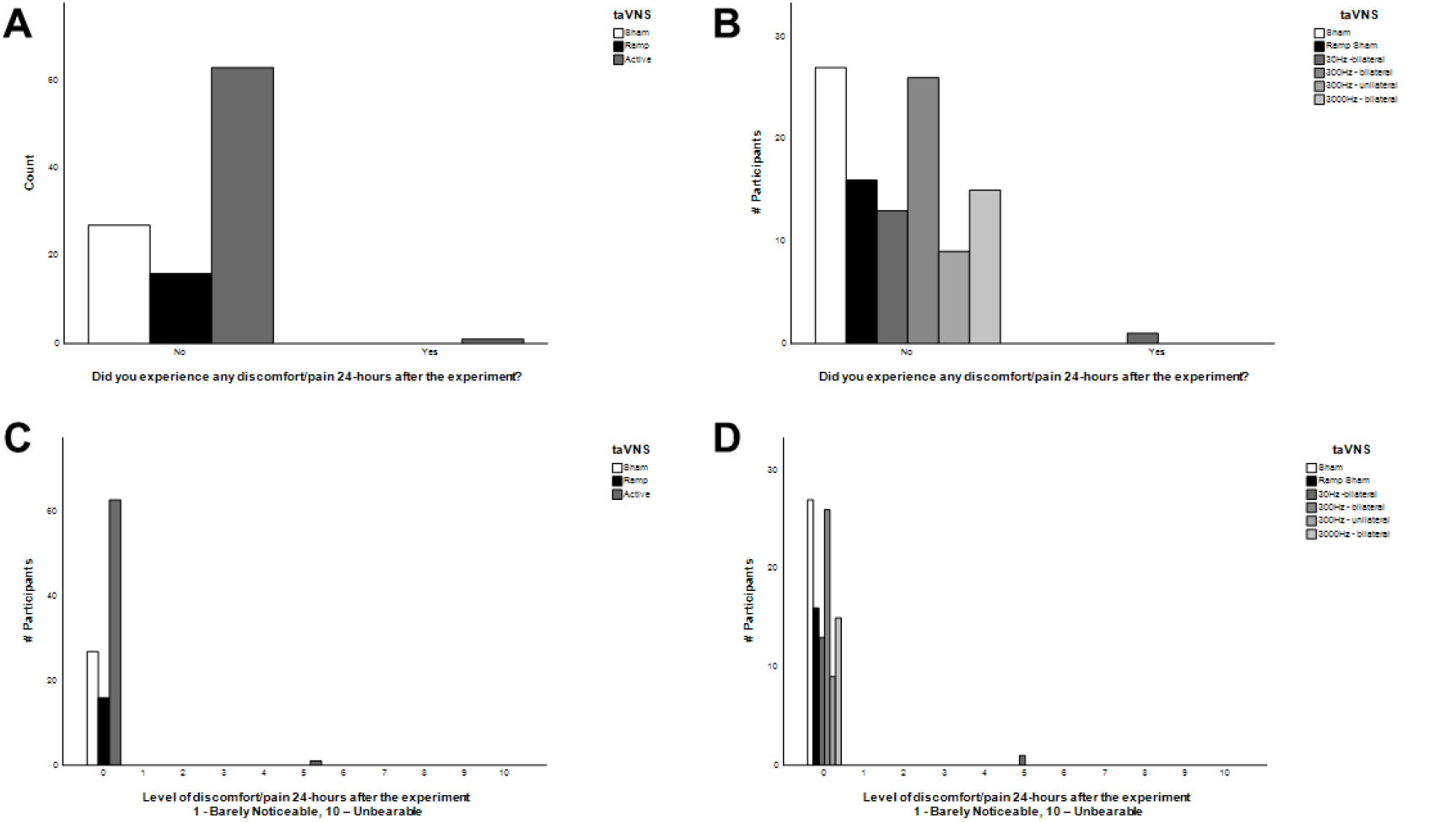
Influence of acute taVNS on discomfort 24 hours following treatment. A) Subject responses are shown for question pertaining to occurrence of discomfort within 24 hours of treatment for inactive sham, ramp sham, and all active taVNS groups collapsed. B) Data as shown in panel A but illustrated by individual taVNS treatment groups. C) Subject responses are shown for question pertaining to the level of discomfort experienced during the past 24 hours following stimulation for inactive sham, ramp sham, and all active taVNS groups collapsed. D) Data as shown in panel C but illustrated by individual taVNS treatment groups.

**Figure 51.**
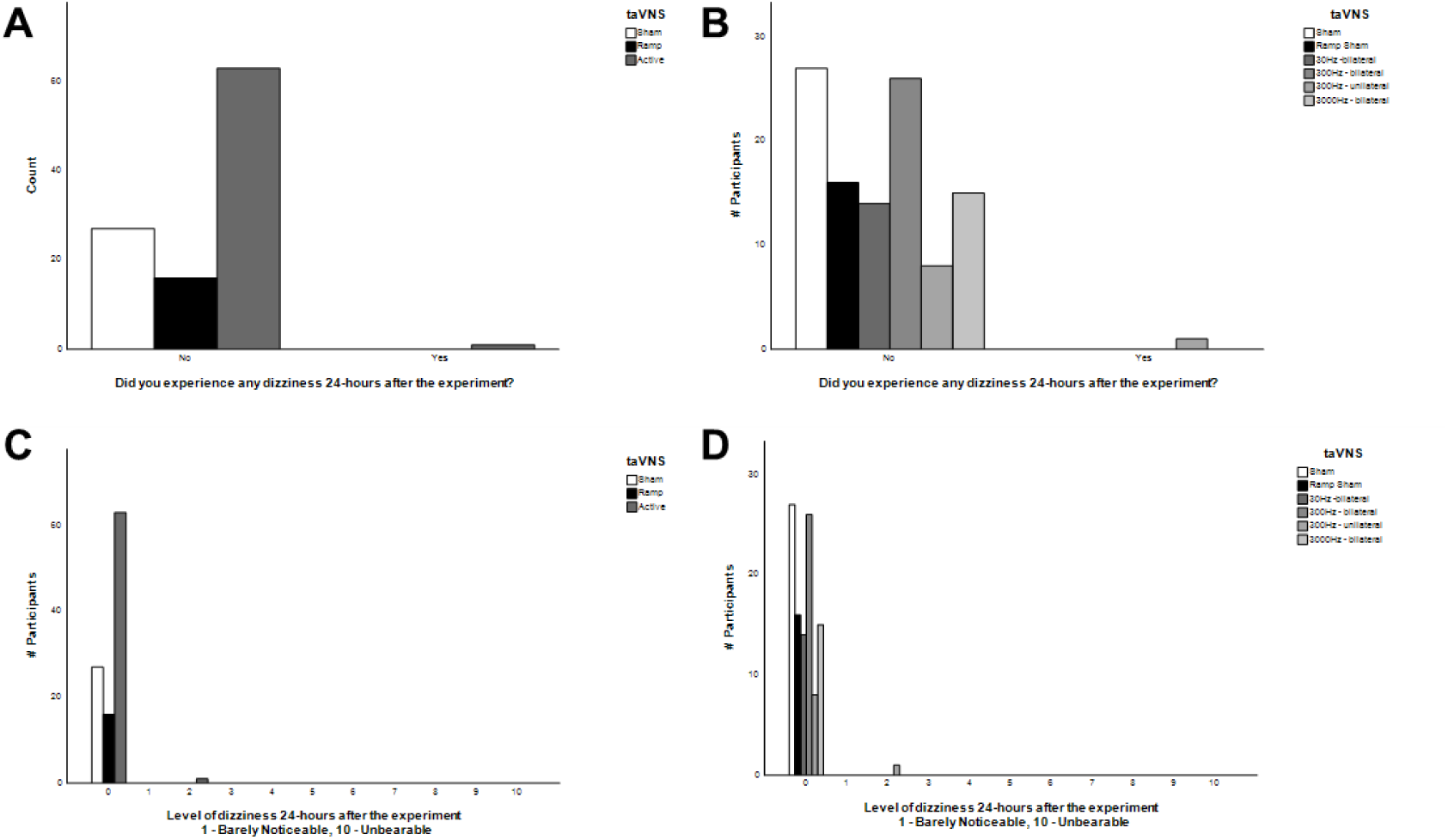
Influence of acute taVNS on dizziness 24 hours following treatment. A) Subject responses are shown for question pertaining to occurrence of dizziness within 24 hours of treatment for inactive sham, ramp sham, and all active taVNS groups collapsed. B) Data as shown in panel A but illustrated by individual taVNS treatment groups. C) Subject responses are shown for question pertaining to the level of dizziness experienced during the past 24 hours following stimulation for inactive sham, ramp sham, and all active taVNS groups collapsed. D) Data as shown in panel C but illustrated by individual taVNS treatment groups.

**Figure 52.**
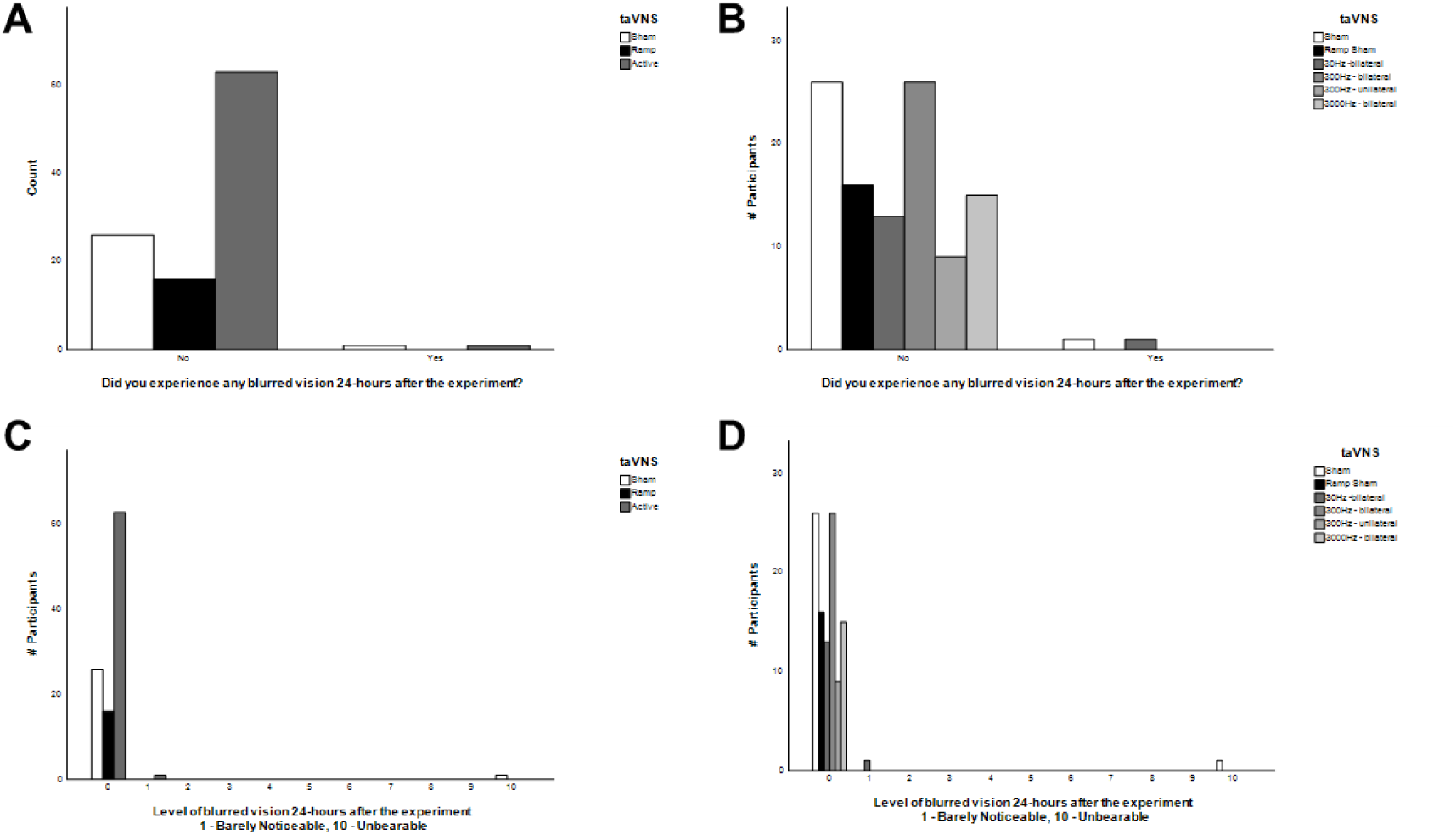
Influence of acute taVNS on vision 24 hours following treatment. A) Subject responses are shown for question pertaining to occurrence of blurred vision within 24 hours of treatment for inactive sham, ramp sham, and all active taVNS groups collapsed. B) Data as shown in panel A but illustrated by individual taVNS treatment groups. C) Subject responses are shown for question pertaining to the level of blurred vision experienced during the past 24 hours following stimulation for inactive sham, ramp sham, and all active taVNS groups collapsed. D) Data as shown in panel C but illustrated by individual taVNS treatment groups.

**Figure 53.**
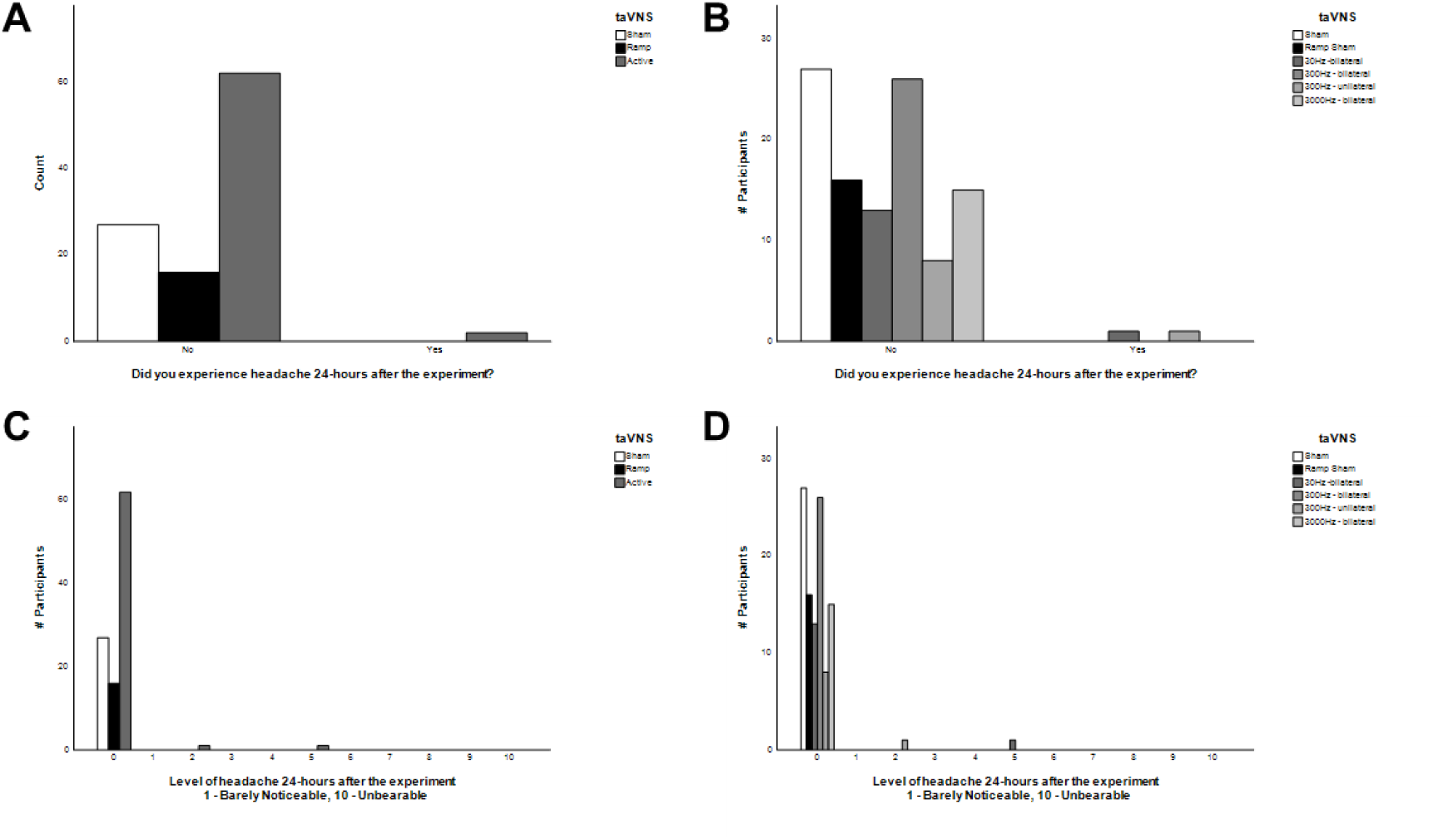
Influence of acute taVNS on headache 24 hours following treatment. A) Subject responses are shown for question pertaining to occurrence of headache within 24 hours of treatment for inactive sham, ramp sham, and all active taVNS groups collapsed. B) Data as shown in panel A but illustrated by individual taVNS treatment groups. C) Subject responses are shown for question pertaining to the level of headache experienced during the past 24 hours following stimulation for inactive sham, ramp sham, and all active taVNS groups collapsed. D) Data as shown in panel C but illustrated by individual taVNS treatment groups.

**Figure 54.**
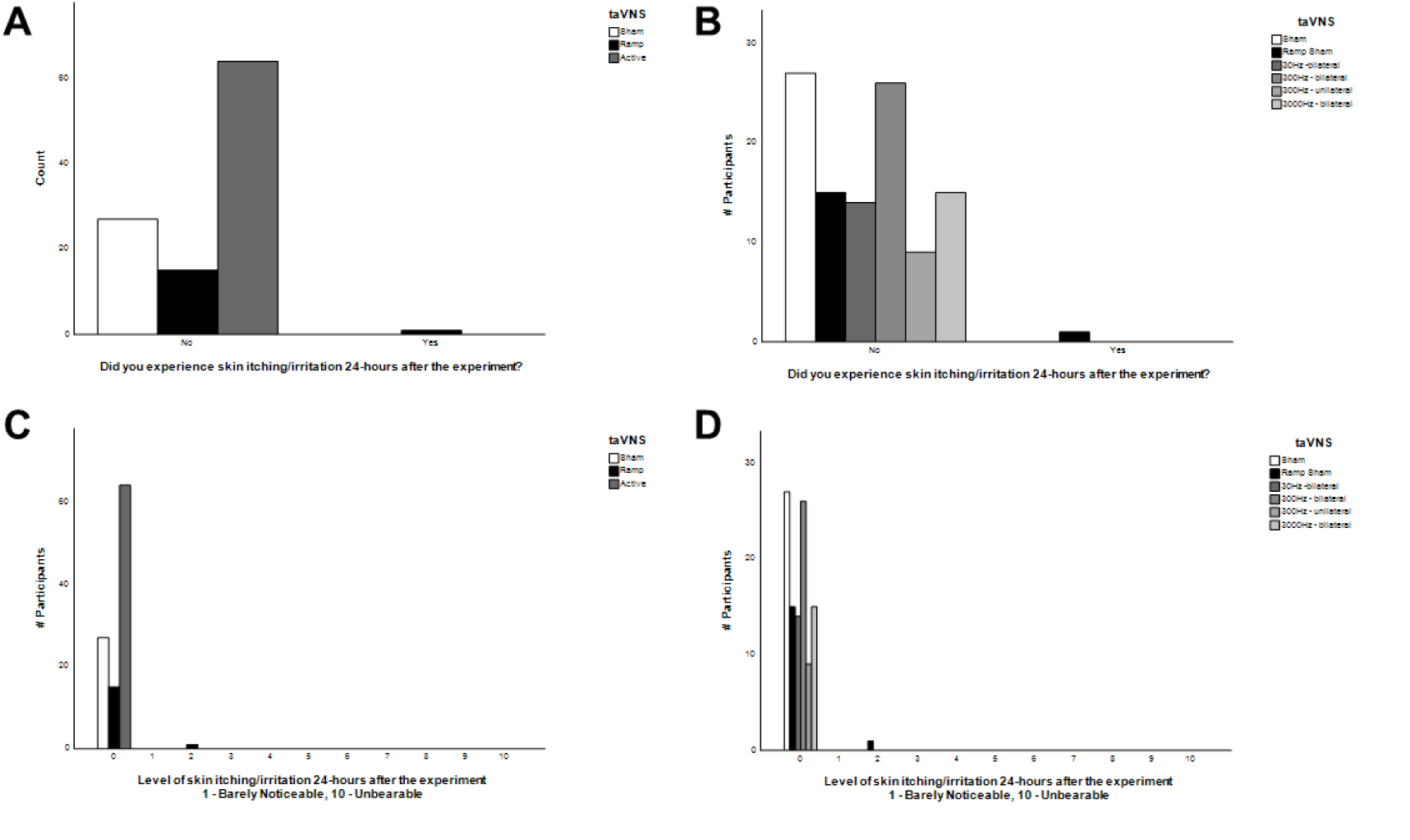
Influence of acute taVNS on skin itching and irritation 24 hours following treatment. A) Subject responses are shown for question pertaining to occurrence of itching and irritation within 24 hours of treatment for inactive sham, ramp sham, and all active taVNS groups collapsed. B) Data as shown in panel A but illustrated by individual taVNS treatment groups. C) Subject responses are shown for question pertaining to the level of itching and irritation experienced during the past 24 hours following stimulation for inactive sham, ramp sham, and all active taVNS groups collapsed. D) Data as shown in panel C but illustrated by individual taVNS treatment groups.

## Brief Discussion

Our work has shown that acute taVNS is safe as a noninvasive neuromodulation method across varied frequencies and pulse shapes. We also found that bilateral taVNS is as safe as unilateral taVNS. Future studies may wish to examine the safety and tolerability of repeated daily use for 1-4 weeks then expand to multi-month investigations where tracking primary safety and health outcomes is a effort designed into clinical investigations or post-market surveillance activities.

With respect to the effects of taVNS on autonomic physiology, it is difficult to determine if one of the taVNS stimulus waveforms is better than another. One of the major limitations in drawing such conclusions is that both the inactive sham and ramp sham treatment condition produced significant effects on autonomic physiology including HR, HRV, SDNN, and others. We hypothesize the robust sham effects on autonomic physiology can be ascribed to a natural relaxation of subjects who sit passively for the most part of an hour. This is a difficult situation to overcome. We have ongoing and planned analyses of the data that will provide finer temporal resolutions. We have analyzed mean data for the last 10 minutes of the baseline, stimulation, and post-stimulus time points. In other cranial nerve modulation experiments including VNS, we have routinely observed a transient 3-4% drop in HR that recovers over the course of about 30 seconds following the onset of stimulation. We plan to reanalyze the data looking at the first tens of seconds of taVNS treatment onset to determine if we can gain greater specificity over some autonomic dependent variables.

In our studies of auditory ERPs we observed weak or marginal effects on brain activity produced by taVNS. In our investigations of continuous EEG spectra and MMN potentials however, we did indeed observe more convincing evidence that taVNS does indeed modulate ongoing and stimulus evoked brain activity. It cannot be determined with any degree of certainty whether one stimulus approach is better than another. More work is required to uncover the rules and regulations for predictively altering brain network activity in response to taVNS and other cranial nerve modulation methods. Whether the influence of taVNS on brain activity is due to direct mechanisms and pathways versus a more general arousal effect also remains unknown.

We noted that there were differences in the current intensity selected by users across the two different types of electrodes. Users of the Nervana modified electrodes used much lower current intensities to reach sensory thresholds compared to those who used custom designed electrodes. This raises an important issue in that one of the main variables appears to be comfort. This point stresses the importance of human factors issues, as well as electrode design factors. Efforts designed to develop hardware for TNT applications should focus on materials that optimize the comfort and efficiency of current delivery to and across the skin.

Further identifying other variables that can lead to robust and predictable outcomes is necessary. Are there good responders and bad responders? How do we identify and screen these individuals? What physiological or psychological factors may contribute to outcomes produced by taVNS? These types of question may be difficult to address in the short-term, but by collecting large data sets we will be able to grasp them. Finally, we conclude that the basic earbud electrode designs described in this report are a comfortable, viable, scalable, rationale, scientifically-valid, and unique method of modulating auricular vagal nerve activity that satisfies several human form factors design requirements for optimizing extended use in real world contexts or environments.

## Disclosures

WJT and NH are inventors or co-inventors on issued and pending patents described herein and pertaining to cranial nerve modulation including vagal nerve modulation. WJT and NH are equity holding co-founders of neurotechnology companies.

## Acknowledgements

We would like to graciously thank Drs. Polly O’Rourke and Stefanie Kuchinsky of the Applied Research Laboratory for Intelligence and Security (ARLIS) at the University of Maryland for their patience, generous sharing of insights, and enduring collaborations. WJT would also like to thank Dr. Henk Haarmann for a decade’s worth of discussions on implementing various neuromodulation methods to enhance foreign language learning and human operator performance. The views, opinions, and/or findings contained in this material are those of the authors and should not be interpreted as representing endorsements, views, or policies or Drs. O’Rourke, Kuchinsky, Haarmann, ARLIS, or the University of Maryland. We simply thank them for their support.

### Funding/Support

This material is based in part upon work supported by the Naval Information Warfare Center (NIWC) and Defense Advanced Research Projects Agency (DARPA) under Cooperative Agreement No. N66001-17-2-4009. The views, opinions, and/or findings contained in this material are those of the authors and should not be interpreted as representing the official views or policies of the Department of Defense, the U.S. Government, or Arizona State University.

